# A lateral temporal network for transmodal combinatorial semantics: Convergent evidence from a multi-study investigation

**DOI:** 10.64898/2026.02.19.706785

**Authors:** Gina F. Humphreys, Matthew A. Lambon Ralph

## Abstract

This study integrates three literatures typically examined in isolation: single-concept semantics, combinatorial semantics, and theory of mind (ToM). We argue that these domains share overlapping computational principles and neuroanatomical networks. Here, we report three major investigations with converging methodological approaches: a meta-analysis of 410 neuroimaging studies, a large omnibus fMRI cross-study comparison (drawing on data from over 150 participants), and two targeted fMRI studies that integrate large language models with traditional psycholinguistic measures, allowing us to quantify combinatorial processing demands and predict brain activation. Across methods, convergent evidence identified a stable, bilateral ATL-STS-TPJ network supporting transmodal combinatorial semantic processing. In addition, the anterior temporal lobe (ATL) showed graded functional specialisation: ventral ATL responded equivalently to single-concept and combinatorial semantics, consistent with a domain-general semantic role, whereas lateral superior ATL was selectively recruited by combinatorial demands. Although ATL-STS-TPJ overlapped with ToM-related activation, targeted control analyses demonstrated that this overlap was eliminated when lexico-syntactic and semantic coherence demands were controlled. Together, these findings support a multimodal combinatorial semantic network centred on bilateral ATL-STS-TPJ with implications for theories of semantic and social cognition. On the basis of these results, we propose a unified theoretical framework of semantic processing.

## Introduction

The aim of this study was to bring together three vast literatures that are typically studied independently: (1) single concept semantics, (2) combinatorial semantics, and (3) theory of mind (ToM) processing. Despite these literatures being studied in isolation there are good reasons to assume some commonality in terms of 1) theoretical overlap, and 2) neuroanatomical overlap. In terms of computational processes, models of single-concept semantics already exist that integrate information across contexts over time (Jackson et al., 2021). With only minor computational adjustments, such approaches could also capture the combinatorial meaning of multiple stimuli from temporally unfolding input (Hoffman et al., 2018). Furthermore, foundational work on ToM, demonstrated that ToM tasks require constructing an integrated meaning from disparate elements, demanding a holistic, global perspective (U. Frith & Happé, 1994), a function akin to that required for combinatorial semantic processing. In terms of neuroanatomy, each domain engages a fronto-temporal network with theoretical importance placed on common areas, for example the anterior temporal lobe (ATL) (C. D. Frith & Frith, 2021; Lambon Ralph et al., 2017; Zaccarella et al., 2017). However, despite potential theoretical and neuroanatomical overlap, the relationship between single-concept semantics, combinatorial semantic processing, and ToM has yet to be explored. Here, we report three major investigations with converging methodological approaches: a meta-analysis of 410 neuroimaging studies, a large omnibus fMRI cross-study comparison (drawing on data from over 150 participants), and two targeted fMRI studies that integrate large language models with traditional psycholinguistic measures, allowing us to quantify combinatorial processing demands and predict brain activation. On the basis of these results, we propose a unified theoretical framework that can explain findings across the three literatures. Such a framework is important both for advancing neurocognitive theory as well as having clinical implications since certain disorders impair social behaviour (e.g., frontotemporal dementia, or traumatic brain injury). The remainder of the Introduction outlines the background for each literature in turn.

### Single concept semantic representation

The hub-and-spoke model proposes that each concept is supported by modality-specific “spoke” regions (for example, visual, auditory, motor) interfacing with a central “hub” anatomically located in the bilateral anterior temporal lobes (Lambon Ralph et al., 2017). Indeed, this is supported by evidence using a wide range of converging methodological approaches, including TMS, fMRI, ECoG, and computational modelling (Lambon Ralph et al., 2010, 2017; Pobric et al., 2007; Rogers et al., 2021; Shimotake et al., 2015). The semantic hub is thought to possess two critical properties that are essential to support long-term conceptual representation. First, it is transmodal, it can accrue knowledge across modalities, thereby embodying a single semantic system that encompasses and generalises across both verbal and non-verbal representations. Secondly, it is transtemporal, the hub draws together information from multiple modalities and experiences over time, thereby allowing for the integration of information across temporal contexts. For instance, one’s knowledge that “birds can fly” and “birds lay eggs” can coexist without experiencing both properties simultaneously. Indeed, this time-extended and context-independent abstraction has been formally demonstrated in computational architectures of semantic cognition (Jackson et al., 2021).

### Combinatorial semantic processing

Given that that the world is constantly changing, understanding how combinatorial semantic processing, that is the temporal integration of semantic information into a coherent representation, is achieved is essential for a complete model of semantic representation. Whilst single concept semantic and combinatorial semantics have typically been studied in separate literatures there are both computational and anatomical reasons to assume that a common underlining mechanism could explain both. First, in terms of computational overlap, whilst the hub-and-spoke model originated as a model single concept representations, hub and spoke computational models already possesses properties that allow it to generalise information across long-range temporal contexts (Jackson et al., 2021), and with only subtle adjustments the model is able to process ongoing temporal input. For instance, (Hoffman et al., 2018) offered a computational framework for combinatorial semantics, whereby a model takes a sequence of single words as input and learns by predicting both the upcoming words and each word’s sensory-motor properties. Through this learning process, the model acquires context-sensitive semantic representations that reflect the multimodal experiential knowledge for each concept while also accounting for co-occurrence and information integration over time (McClelland, St. John, et al., 1989). Furthermore, like in the single-concept semantic model, this combinatorial framework has been shown to apply transmodally, for instance predicting visual objects from their surrounding context (Sadeghi et al., 2015). Indeed, transmodal processing would presumably vitality important function for real world combinatorial semantics given the continuous co-occurrence of verbal and nonverbal input in the natural world.

Next, in terms of anatomical overlap, as described above, the hub-and-spoke model proposes that the ATL is acts as a transmodal and transtemporal hub for semantic representations. The ATL has also been implicated in combinatorial semantic processing. For instance, in the language domain, a meta-analysis of neuroimaging studies that compare sentences > word lists found that combinatorial processing in language also involves the ATL in addition to a more extensive network along the length of the superior temporal sulcus (STS) to the temporo-parietal junction (TPJ) (hence forth the ATL-STS-TPJ network) (Zaccarella et al., 2017). Furthermore, this same network appears to be transmodal, being also engaged by nonverbal paradigms such as movie watching with or without verbal input (Hasson et al., 2008; Lerner et al., 2011; Sueoka et al., 2024). Saying this, there are several reasons to be cautious when interpreting the combinatorial data: 1) there are comparatively few studies that investigate combinatorial semantics, particularly in the nonverbal domain, 2) nonverbal studies have tended to rely more on specific paradigms, such as movie watching (Hasson et al., 2008; Lerner et al., 2011; Sueoka et al., 2024), which may lack experimental control, and 3) importantly, little/no study has directly compared verbal and nonverbal combinatorial semantics in a within-subject controlled experiment.

### Theory of Mind processes

Theory of Mind (ToM) refers to the capacity to understand that other people have thoughts, feelings, and intentions that may differ from one’s own. ToM and semantic domains have been studied almost entirely separately to date. Despite this separation there are several reasons to suspect meaningful overlap between them. First, in terms of neuroanatomical convergence, meta-analyses of ToM studies have consistently shown recruitment of the STS–TPJ network, which is also strongly associated with combinatorial semantics (Schurz et al., 2021). Moreover, the limited work that does exist has shown that the semantic memory and ToM processes engage overlapping networks in meta-analyses (Balgova et al., 2024). Indeed, it is intuitive that ToM must draw on the semantic memory system to some degree: for instance, integrating knowledge about identity, roles and context is vital to recognise social cues and generate socially appropriate responses (Balgova et al., 2024). Going further, foundational theoretical accounts of ToM proposed that a key, central component involves integrating diverse information into a coherent whole, requiring a global, holistic perspective rather than simple additive processing (U. Frith & Happé, 1994). Contemporary models similarly suggest that the anterior temporal lobe (ATL) contributes by providing access to “social scripts”, that is context-rich templates for typical social interactions (C. D. Frith & Frith, 2021). These ideas closely mirror principles from hub-and-spoke and sentence gestalt models of semantic integration (Hoffman et al., 2018; McClelland, St. John M, et al., 1989).

### Is there likely to be a one-to-one mapping across these three domains?

Therefore, given the apparent computational and anatomical overlaps, it would be surprising if there were no relationship between the mechanisms engaged by single-concept semantics and combinatorial semantics. That being said, it is unlikely that there is a simplistic one-to-one mapping between the two processes. Mechanistically, combinatorial processing involves the integration of continuously unfolding inputs and therefore depends more heavily on online integrative mechanisms. Indeed some dorsal posterior temporal/parietal regions seem to function as temporary buffers for sequential information (Humphreys et al., 2021; Humphreys & Lambon Ralph, 2015), and computational models indicate that, whilst the ventral pathway integrates semantic information without maintaining a veridical, reproducible code of input order, the dorsal stream learns to represent the sequential ordering of inputs (Ueno et al., 2011). Anatomically, whilst there have been few/no direct comparisons across domains, combinatorial semantics appears to be more typically associated with a more superior-lateral region of the ATL, and also engages a broader ATL-STS-TPJ network (Hasson et al., 2008; Lerner et al., 2011; Sueoka et al., 2024; Zaccarella et al., 2017) consistent with a greater reliance on a dorsal processing route. In contrast, in single-concept semantics particular emphasises has been placed on the ventrolateral ATL, as being centre point for a graded representational hub region (Lambon Ralph et al., 2017) (Binney et al., 2010; Sato et al., 2021). However, the absence of ventral ATL activation for combinatorial semantic may have several possible explanations: 1) there may be modular segregation within the ATL, such that single-concept and combinatorial semantics engage distinct subregions; 2) the ATL might function as a graded semantic hub, with some regions being more temporally sensitive (e.g., superior ATL) and others, such as ventral ATL, supporting more stable, long-term representations; 3) the apparent dissociation may reflect methodological limitations, particularly the lack of ventral ATL signal coverage in most standard fMRI protocols due to magnetic susceptibility artefacts and lack of an active baseline control task. Indeed, multi-echo fMRI methods with enhanced signal coverage and sue of active baseline tasks have demonstrated reliable ventral ATL activation for single-concept semantics (Halai et al., 2014, 2025; Humphreys et al., 2015)(Halai et al., 2014), but similar optimised approaches have not yet been applied to investigate vATL engagement in combinatorial studies.

As with single-concept semantics, there is also reason to assume that there may not be a one-to-one mapping between combinatorial semantics and ToM processing. Indeed, several authors have argued that language, semantics and ToM rely on distinct specialised networks (Olson et al., 2013; Saxe & Powell, 2006; Shain et al., 2023). Apparent neuroanatomical convergence between them could arise for several reasons: 1) neighbouring but distinct systems – ToM and semantic processing may depend on adjacent but separate networks. The limited number of direct ToM–semantic comparisons, and the reliance on meta-analyses (which lack the anatomical precision of within-subject designs), may obscure clear boundaries; 2) content confounds – Many combinatorial semantic studies use human- or socially themed material, which could bias results toward overlap with ToM regions; 3) task demands – ToM tasks may inherently place greater demands on the combinatorial semantic system, as they require the continuous integration of semantic information over time to understand complex, inferential social events. Indeed, evidence suggests that ToM effects within the language network can disappear when linguistic variables are appropriately controlled (Shain et al., 2023). Disentangling these alternatives will require direct, within-subject comparisons using carefully controlled experimental manipulations.

### Keys questions and aims

The overarching aim of the present study was to unite the literature on single-concept semantics, combinatorial semantics, and Theory of Mind (ToM) by delineating the neural architecture supporting combinatorial semantic processing and its relationship to single concept semantics and social-cognitive inference. Specifically, we sought to determine whether 1) combinatorial semantics engages a stable and reliable neural network, 2) the combinatorial semantic network is multimodal, 3) to what extent the combinatorial semantic network overlaps with single concept semantics, particularly in terms of ATL engagement, and 4) how combinatorial semantics relates to ToM processing: are the combinatorial semantic and ToM networks overlapping or functionally adjacent, and if they do overlap, to what extent any apparent overlap can be explained by the combinatorial demands inherent in ToM tasks.

These questions were addressed through a convergent combination of three major investigations, each with distinct methods. Using multiple methods makes it possible to leverage the strengths of each technique while minimising their individual limitations, thereby providing a basis for powerful converging evidence. First, we used ALE meta-analyses to determine the similarities and differences between the networks engaged by single-concept compared, combinatorial semantics processing, and verbal and nonverbal ToM task. Meta-analyses provide a powerful means of identifying robust patterns that are consistent across studies and independent of study-specific idiosyncrasies. However, because they rely on published data, they are susceptible to methodological or sampling biases, such as the predominance of verbal over non-verbal combinatorial tasks, as well as limitations in imaging protocol in terms of ATL signal detection (Halai et al., 2014; Visser et al., 2010), a major limitation given the theoretical importance of the ATL to the research question. Combining meta-analytic results with data from within-subject fMRI studies thus provides a more comprehensive understanding of the underlying neural architecture.

Second, an omnibus fMRI analysis combined 17 tasks from eight studies and 151 participants, all using fMRI protocols that enhance ATL signal detection (Halai et al., 2014, 2025). Here we directly compared the networks engaged by combinatorial versus single-item tasks, semantic versus non-semantic combinatorial processing, and the interaction of the network with tasks involving social cognition. The inclusion of fMRI studies that have good ATL signal coverage is critically important to the question of whether the same region of ATL engaged by single-concept compared to combinatorial semantic task e.g. the role of ventral vs. superior ATL. This approach is advantageous over ALE meta-analyses as it allows direct task comparisons, provides full spatial resolution across all voxels (rather than just the point of peak activation) and incorporates data from a large number of participants and paradigms. Whilst this approach provides higher spatial resolution than meta-analysis, it is still constrained by cross-study variability, uneven task types, and paradigms not specifically designed to maximally challenge the combinatorial semantic system. These limitations were addressed by the third approach.

Third, we conducted two targeted and well-controlled within-subject fMRI studies. The first experiment employed a 2 × 2 combinatorial semantic manipulation, comparing verbal and non-verbal coherent compared to scrambled narratives. In the second experiment we compared activation for ToM vs. non-ToM narratives, and examined the extent to which network engagement could be explained by increased demands on the combinatorial semantic system by systematically varying computational combinatorial load using measures derived from large language models alongside traditional psycholinguistic metrics.

Together, these three complementary approaches allowed us to rigorously address our key objectives: (1) to establish converging evidence of the existence of a robust combinatorial semantic network, (2) to determine its multimodal nature, and to clarify its relationship with 3) single concept semantics 4) and ToM processing, in order to determine the extent of genuine mechanistic overlap.

## Section 1: ALE meta-analysis

An ALE meta-analysis of neuroimaging studies was conducted with the key aim to examine to what extent studies investigating combinatorial semantics overlap with single-concept semantics, and ToM processing. For semantic tasks we chose to focus only verbal semantics given the scarcity of nonverbal task particularly in terms of combinatorial semantic paradigms. Using GingerAle 3.0.2 (Eickhoff et al., 2009; Laird et al., 2005) we calculated the activation likelihood estimate (ALE) for studies from each cognitive domain: 1) combinatorial semantic tasks (133 fMRI or PET studies involving sentence-level semantic manipulations), 2) single-word semantic manipulations (93 studies), and 3) studies including some form of ToM manipulation (113 studies; separated into 71 verbal studies, and 42 nonverbal studies) applying voxel-level thresholding at a p-value of 0.001 and cluster-level FWE-correction with a p-value of 0.05 over 10,000 permutations. The results can be visualised in Figure 1 (left panel), and for a detailed description of the methods and results refer to Supplementary 1.

**Figure 1.**
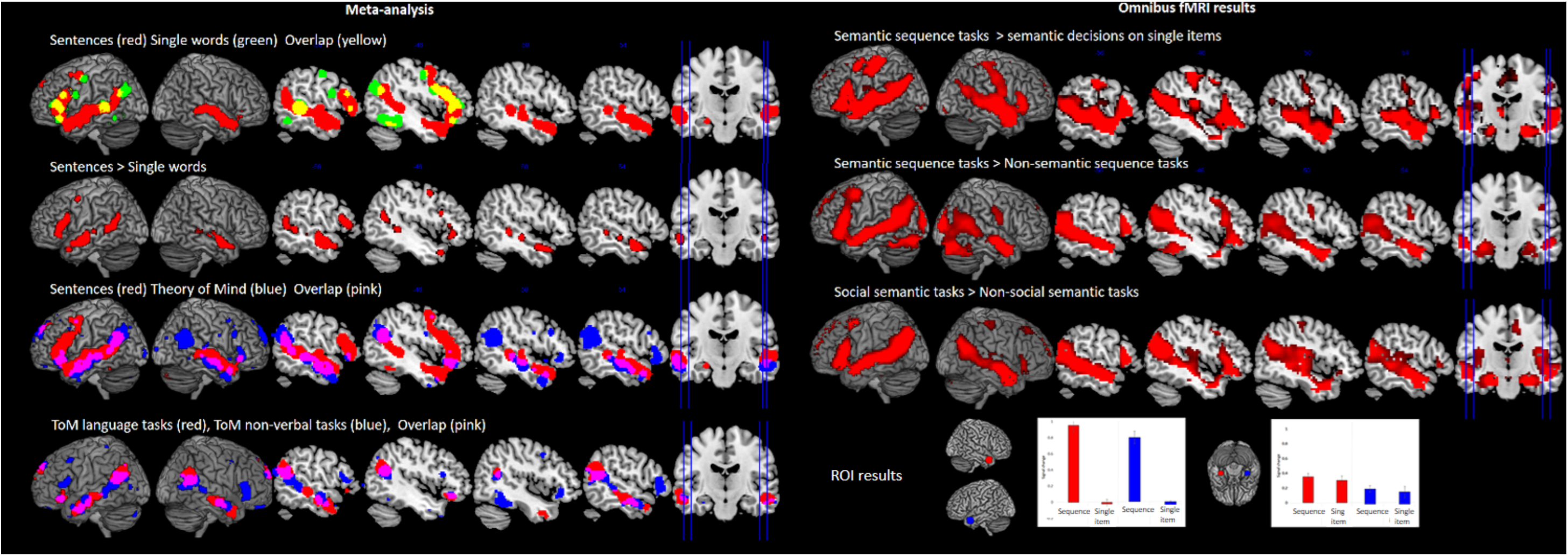
Left panel: The results from the ALE meta-analysis (voxel-level thresholding at a p-value of 0.001 and cluster-level FWE-correction with a p-value of 0.05 over 10,000 permutation). Right top panel: The results from the omnibus fMRI study. Group analyses were conducted using standard voxel height threshold p < .001, cluster corrected using FWE p < .05. Right bottom panel: ROI analysis showing the % signal change from rest for the semantic sequence tasks (combinatorial semantics) compared to the single item semantic tasks from Temporal Pole and vATL.

### Sentence-level semantic tasks compared to single-word tasks

Sentence-level semantic task were found to reliably engage the length of the ATL-STS-TPJ network, from the anterior temporal pole until the temporo-parietal junction (TPJ). Recruitment was bilateral, although extending more posteriorly in the left hemisphere. Within frontal cortex, reliable activation was found in left frontal areas within precentral gyrus, inferior frontal gyrus (IFG, BA44, BA45), and supplementary motor area. In contrast, single-word semantic studies were found to reliably elicit a more restrictive network and was limited to the left hemisphere. This included clusters with IFG, premotor cortex, posterior middle temporal gyrus, and angular gyrus, but not the more anterior portion of the temporal lobe or the TPJ. The two networks overlapped in LIFG, pMTG, and the anterior potion of the AG. Contrast analyses revealed that combinatorial semantic studies were statistically more likely to engage the bilateral ATL-STS-TPJ network, as well as LIFG (see Figure 1: left panel). This effect remained when controlling for task input modality (visual vs. auditory stimulus type).

### Sentence tasks and Theory of mind tasks

Next we compared the networks engaged by sentence-level semantic tasks to those involving some form of ToM manipulation. There was a striking overlap between both domains along the length of the ATL-STS-TPJ network bilaterally, as well as the LIFG. Given that it is possible that the high degree of sentence-level semantic and ToM overlap could be driven by the inclusion linguistic task in the ToM analysis. We therefore performed an ALE analysis but this time separated the verbal ToM paradigms (e.g. false belief narratives) and non-verbal ToM paradigms (e.g. abstract shape animations). The resultant ALE maps were largely overlapping across modalities, with no statistically significant difference between the two (see Figure 1, left panel). Therefore, the overlap between sentence-level semantics and ToM manipulations is unlikely explained by input modality

## Section 2: fMRI omnibus study

Next, we conducted a large omnibus fMRI analysis using 17 tasks from eight studies (N = 151) all of which used fMRI protocols that enhance ATL signal detection. The primary aims of omnibus fMRI study was to replicate and extend to findings from Section 1. Specifically, 1) we examined the network engaged by combinatorial semantic tasks, such as narrative comprehension or narrative speech production, as compared to single-item semantics, such as picture naming. This was done at both a whole-brain level as using an ROI approach to investigate potential differences in the vATL vs. superior temporal polar region. 2) Next we sought to determine the extent to which the ATL-STS-TPJ network is engaged primarily for combinatorial semantics, or whether it is also engaged by non-semantic tasks that require combinatorial integration over time i.e. a general combinatorial processing network (e.g. listening to unfamiliar classical music, or processing numerical sequences). Lastly, 3) we examined the extent to which combinatorial semantics engaged a similar network to social semantic processing, as defined as a task where the stimuli contain human participants as opposed to objects or non-human animals.

To address these three questions, the 17 tasks were subdivided into subtypes: combinatorial tasks (with or without semantic content), single concept semantic tasks, and social vs. non-social semantic tasks (see Table 1). These tasks were entered into a flexible factorial ANOVA in SPM12 included factors for subject (non-independent, equal variance), study (8 level), and task (17 levels). This structure accounted for repeated measures within subjects and heterogeneity across datasets. Planned contrasts then tested the comparison between combinatorial semantic tasks > single-item semantic tasks, combinatorial semantic tasks vs. non-semantic sequential tasks, and social semantic > non-social semantic tasks. Group analyses were conducted using standard voxel height threshold p < .001, cluster corrected using FWE p < .05. For detailed description of the tasks and analyses see Supplementary 2.

**Table 1.**
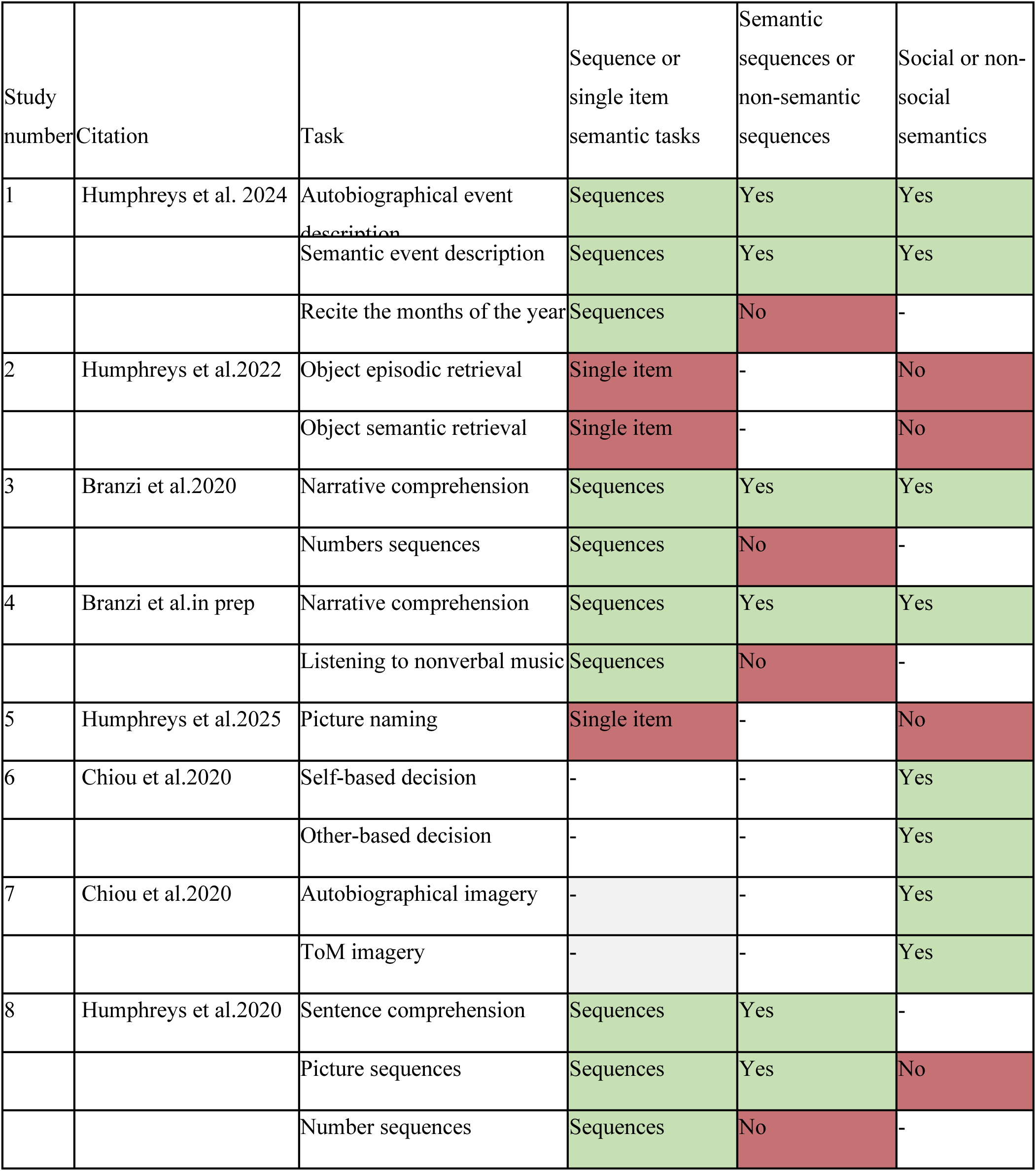
A summary of the studies included in the omnibus fMRI analysis.

| Study number | Citation | Task | Sequence or single item semantic tasks | Semantic sequences or non-semantic sequences | Social or non-social semantics |
| --- | --- | --- | --- | --- | --- |
| 1 | Humphreys et al. 2024 | Autobiographical event description | Sequences | Yes | Yes |
|  |  | Semantic event description | Sequences | Yes | Yes |
|  |  | Recite the months of the year | Sequences | No | - |
| 2 | Humphreys et al.2022 | Object episodic retrieval | Single item | - | No |
|  |  | Object semantic retrieval | Single item | - | No |
| 3 | Branzi et al.2020 | Narrative comprehension | Sequences | Yes | Yes |
|  |  | Numbers sequences | Sequences | No | - |
| 4 | Branzi et al.in prep | Narrative comprehension | Sequences | Yes | Yes |
|  |  | Listening to nonverbal music | Sequences | No | - |
| 5 | Humphreys et al.2025 | Picture naming | Single item | - | No |
| 6 | Chiou et al.2020 | Self-based decision | - | - | Yes |
|  |  | Other-based decision | - | - | Yes |
| 7 | Chiou et al.2020 | Autobiographical imagery | - | - | Yes |
|  |  | ToM imagery | - | - | Yes |
| 8 | Humphreys et al.2020 | Sentence comprehension | Sequences | Yes | - |
|  |  | Picture sequences | Sequences | Yes | No |
|  |  | Number sequences | Sequences | No | - |

## Section 2: Results

### Whole-brain analyses

#### Combinatorial semantic tasks > single-item semantic tasks

Replicating the results from the meta-analysis, combinatorial semantic tasks more strongly engaged bilateral activation extended the full length of the ATL-STS-TPJ network compared to single-concept semantics, as well as frontal activation including lateral frontal areas within bilateral IFG and left premotor cortex, as well as medial frontal areas within SMA.

#### Combinatorial semantic tasks > non-semantic sequences

Again, this contrast revealed bilateral activation extended the full length of the ATL-STS-TPJ network. In the frontal cortex, activation included the LIFG and premotor cortex. Therefore, it appears the ATL-STS-TPJ network is specific to combinatorial semantic processing rather than combinatorial processing in general (see Figure 1, right panel).

#### Social tasks > non-social semantic tasks

Again, this contrast revealed activation along the entire bilateral ATL-STS-TPJ network, as well as bilateral IFG and premotor cortex. Hence, the results could equally be explained by the semantic gestalt as well as a ToM-specific, or more general “social semantic”, network prediction (see Figure 1, right panel).

### ROI analysis: The contribution of the ventral ATL compared to the superior Temporal Pole

A large body of work has highlighted to the importance of the ventral ATL to single concept semantics and particular prominence is given to vATL in neurocomputational models, yet existing evidence tends to highlight activation for combinatorial semantics in the superior-lateral temporal pole. It is unclear at this stage whether this reflects 1) a true difference in vATL vs. superior-lateral temporal polar response 2) a methodological artefact that has arisen due to signal loss in vATL that is associated with standard fMRI. Therefore, here utilising paradigms with enhanced ATL signal detection we directly compared activation for the single-concept vs. combinatorial semantic tasks in *a priori* defined bilateral vATL (Binney et al., 2010) and bilateral superior lateral temporal pole ROIs taken from peak coordinates from the meta-analysis in Section 1.

A two-way ANOVA with showed a significant ROI x condition interaction in the left-hemisphere (F(149) = 41.72, p < .001) and right-hemisphere (F(149) = 33.84, p < .001). In the temporal pole, independent samples t-tests tests assuming unequal variance showed significantly stronger activation for the combinatorial semantic compared to single concepts semantic tasks in both the left - (t(148.80) = 9.38, p < .001) and right-hemisphere (t(134.37) = 8.25, p < .001). Activation was significantly above rest for the combinatorial semantic condition (all ts > 10.12 all ps < .001) but did not differ from rest for single concept semantics (all ts < -.86, all ps > .20) (see Figure 1, lower right panel). In contrast, independent Samples t-tests assuming unequal variance showed that the vATL responded equally strongly to the combinatorial semantic and single concept semantic tasks in both the left- (t(94.50) = .66, p = .55) and right-hemisphere (t(68.98) = .46, p = .65), with the activation significantly above rest in all conditions (all ts > 2.16, all ps > .01). These results support the notion that vATL is involved in *all* forms of semantic processing, whereas the superior temporal pole appears specifically involved in combinatorial semantic tasks.

### Summary

There were two key findings from the omnibus fMRI study. Firstly, the results replicated and extended the results from the meta-analyses, showing the importance of bilateral ATL-STS-TPJ network in combinatorial semantic processing and/or social-semantic processing/ToM. Secondly, the ROI analyses demonstrate the vATL and superior lateral surface within the temporal pole respond differently to combinatorial semantics tasks compared to single concept semantics: Whereas the vATL responds equally strongly regardless of task types, supporting its more general role in all forms of semantic processing, the lateral temporal polar area responds selectively to tasks involving combinatorial semantics. These results will be clarified and extended in the next section.

The next section will go on to replicate and extend the current results

## Section 3. Targeted fMRI studies

The third approach involved two targeted and well-controlled within-subject fMRI studies that replicate, extend, and clarify the results to section 1 and section 2 using an fMRI protocol with enhanced ATL signal coverage. For a detailed description of the tasks and results see Supplementary 3.

### Experiment 1

The aims of Experiment 1 were as follows: 1) To directly compare the networks engaged by combinatorial coherent items that elicit a semantic gestalt compared to combinatorial incoherent items in a well-controlled within-subject experiment 2) To determine the transmodal nature of this network; is the same network engaged by combinatorial stimuli in both the verbal and nonverbal stimuli 3) To examine whether responses to coherence across the ATL, specifically within the vATL as compared to the superior temporal pole.

Experiment 1 employed a 2 × 2 combinatorial semantic manipulation, comparing verbal and non-verbal (picture) stories which either followed a coherent or semantically scrambled content. In the verbal task, the participants were visually presented with short three sentence narratives (e.g., “Betty’s flight just arrived at the airport. She walks toward the baggage claim area and picks up her suitcase. After taking a cab home, she rushes to her room and gets into bed”). The scrambled verbal stories were created using a separate but psycholinguistically matched set of items whereby the three sentences were pseudo-randomly swapped across narratives (sentence 1 with sentence 1, sentence 2 with sentence 2, and sentence 3 with sentence 3) such that the story no longer form a coherent meaningful narrative (e.g. “Earlier today, Kevin played in the snow with his best friend Joe. Arriving at the station, she runs toward the platform. Then, for the next hour, he lifts weights and does crunches”).

The non-verbal stories were made up of comic book like sequences. Each story was made of nine picture sequence event with at least one human character. Each of the nine-picture sequences were subdivided into three three-picture sequences. The three-picture events were then presented in either their usual order to create a clear coherent semantic narrative (coherent picture condition) or using a separate set of pictures they were pseudo-randomly shuffled across items in order to create an incoherent narrative (scrambled picture condition).

### Whole-brain analyses contrasting against rest

In order to examine the neural networks engaged by each condition, and the overlap between conditions, we first contrasted each condition relative to rest. Group analyses were conducted using standard voxel height threshold p < .001, cluster corrected using FWE p < .05 (see Figure 2).

**Figure 2.**
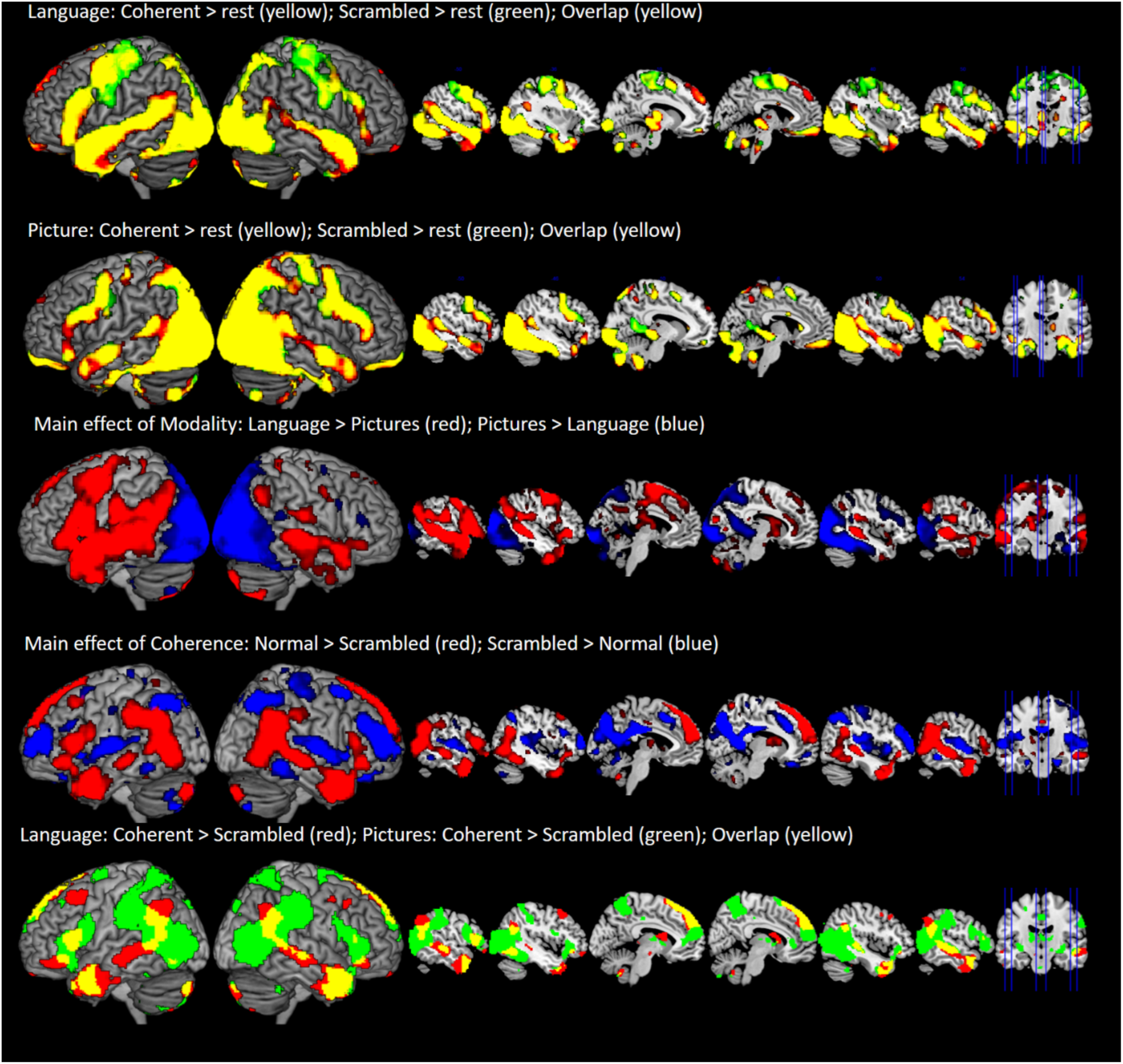
Experiment 1 whole-brain fMRI results. Group-level analyses employed a voxel-wise threshold of p < .001, with cluster-level family-wise error correction at p < .05.

#### Language tasks

Relative to rest, coherent stories engaged a fully bilateral fronto-temporo-parietal network, as well as visual processing areas. Specifically, in the temporal lobe, activation spanned the length of the ATL-STS-TPJ, including the entirety of the temporal pole, as well as the posterior-anterior fusiform gyrus. Frontal activation included lateral frontal cortex (IFG, middle frontal, premotor cortex) as well as medial frontal areas (SMA, vMPFC, dMPFC, orbito-frontal cortex), and parietal activation included SMG, anterior AG, as well IPS. This activation pattern overlaps with the known language network, as well as the MD system. Interestingly, the scrambled language condition > rest engaged an entirely overlapping network, the only visual difference being a slightly reduced volume of activation in the temporal lobe but mildly more extensive in fronto-parietal areas. This suggests that any significant difference between conditions will be quantitative rather than reflecting a qualitatively different pattern of recruitment.

#### Picture tasks

Relative to rest, the coherent picture condition engaged a strikingly similar neural network as the language task, engaging a largely overlapping bilateral fronto-temporo-parietal network. This included the length of the ATL-STS-TPJ, including the entirety of the temporal pole, as well as the posterior-anterior fusiform gyrus. Frontal activation included lateral frontal cortex (IFG, middle frontal, premotor cortex) as well as medial frontal areas (SMA, vMPFC, dMPFC, orbito-frontal cortex), and parietal activation included SMG, anterior AG, as well IPS. In addition to this, as expected the visual task also engaged an extensive occipito-parietal activation, spanning the entirety of the occipital lobe, and extending dorsally into superior parietal cortex, as well as the posterior AG and IPS. As in the language task, the scrambled picture condition engaged a fully overlapping network to the coherent picture condition, with only moderate visual differences in cluster size. Together these results provide evidence that both the language and picture task engage a common multi-modal semantic network.

### Factorial ANOVA

In order to conduct direct task comparisons, the data were analysed using a 2 x 2 (Language vs. Pictures) factorial ANOVA testing for a main effect of Coherence (Coherent vs. Scrambled) and Modality (Language vs. Picture-cartoons), as well as the interaction. Group analyses were conducted using standard voxel height threshold p < .001, cluster corrected using FWE p < .05 (see Figure 2). Despite the high degree of overlap across networks, the results from the factorial ANOVA indicate a significant main effect of modality, coherence, as well as a modality x coherence interaction.

#### Modality

In terms of modality, relative to the picture task, the language conditions showed stronger engagement of the bilateral fronto-temporal system that has been commonly associated with language processing elsewhere (refs), whereas as expected, the picture task more strongly engaged posterior occipito-parietal areas, as well as the ventral temporal lobe in areas associated with visual recognition.

#### Coherence

Consistent with our prior results, the Coherent > Scrambled contrast showed bilateral recruitment of the length of the ATL-STS-TPJ network with activation including the full extent of the temporal pole, and a large TPJ/pMTG cluster. In addition to this, the bilateral IFG and medial dorsal prefrontal cortex were also recruited more strongly for the coherent compared to the scrambled conditions. In contrast, the Scrambled > Coherent conditions revealed a bilateral fronto-parietal network that is commonly associated with executive processing and overlapped with the MD system. This included bilateral middle frontal gyrus, premotor cortex, SMA, the IPS, and posterior ITG

#### Interaction

A significant interaction was revealed in the left IFG, left ATL-STS-TPJ network, as well as large areas of bilateral parietal cortex, including TPJ, SMG, and the SPL. Direct task comparisons showed that this pattern reflected a larger coherence effect in the left STS for the Language compared to the Picture task, but a stronger coherence effect for the Picture task in bilateral parietal cortex compared to the Language task.

#### Summary

The key findings were that 1) a common fronto-temporo-parietal network responds to all language and picture tasks above rest, providing evidence of a multimodal network for semantic processing. In the temporal lobe, activation included the ventral ATL, as well as the entire bilateral ATL-STS-TPJ network. 2) The entire bilateral ATL-STS-TPJ network it is sensitive to coherence in both the picture and language domain – it responds more strongly coherent compared to scrambled conditions in both tasks. Note that this is in contrast to the vATL which is actively engaged by all tasks equally. 3) Within the ATL-STS-TPJ network there are relative modality differences: there is a larger coherence effect in the mid portion of the left STS Language compared to Picture task, but a stronger coherence effect for the Picture task in bilateral parietal cortex compared to the Language task. This is contrast to the bilateral temporal pole which shows a comparable coherence effect across tasks. These results are further explored in targeted ATL-STS-TPJ ROI analyses below.

### ROI analyses

#### ATL-STS-TPJ vector ROIs

First, in order to investigate our specific hypotheses with regard the ATL-STS-TPJ network, bilateral vector ROIs were defined stretching from the anterior temporal pole along the length of STS to the TPJ, Beginning with the ATL-STS-TPJ vector ROIs, the mean signal for each subject was entered into a 2 x 2 x 6 within-subject ANOVA, where we tested for the main effects and interactions between Coherence, Modality, and ROI anterior-posterior location. To simplify the interpretation of results we initially tested for Coherence, Modality, and ROI effects within the left- and right-hemisphere separately, before conducting cross-hemisphere comparisons (See Figure 3). The key results were as follows (full results are reported in Supplementary 3). The full in depth ANOVA is reported in Supplementary but the principle findings were:

1. All tasks activated all ROIs: Regardless of coherence or modality, task activation was statistically above rest for all tasks in every ROI (all ts > 3.51, p < .001) (although only marginally for the right STS4 for the scrambled picture task (t = 2.11, p = .02)). The only regions to show no above rest activation were within the left STS3 for the coherent and scrambled picture condition, and the right STS3 for the scrambled picture condition in right STS3.
2. All areas showed a significant effect of coherence: There was a main effect of coherence in both hemispheres (left: F(42) = 68.79, p < .001; right (F(42) = 93.75, p < .001), with pairwise comparisons showing significant increased activation for the coherent vs. scrambled condition in all ROIs (all ts > 3.76, p < .001, with the exception of LSTS2 which showed a marginal difference t(42) = 2.67, p = .005). The exception being no significant effect in right TPJ for the language task (t(42) = .24, p =.4).
3. There were graded modality differences: There was a significant modality x ROI interaction in both the left and right hemispheres (left: F(42) = 71.72, p < .001; right: (F(42) = 38.69, p < .001). This reflected a larger Language > Picture difference in the middle portions of STS closest to auditory cortex bilaterally (for STS3 and STS4) compared to all other regions (all ts > 4.91, all ps < .001, with the exception of only a marginal difference with LSTS5 (t(42) = 2.70), and in the right hemisphere, a larger Picture > Language effect in TPJ compared to all regions (all ts > -5.22, all ps < .001). In contrast, both the left- and right-hemispheres the modality difference was smallest in the most anterior ROIs (left STS1: all ts > 3,60, all ps < .001; right: STS1: all ts > -3.27, all ps < .002); and STS2 all ts > -3.96, all ps < .001).
4. There were graded laterality differences: 1) There was a main effect of hemisphere (F(42) = 5.37, p = .03) reflecting stronger overall activation in the left compared to right hemisphere across tasks (t(42) = 2.32, p = .01), 2) a significant effect of modality (F(42) = 79.26, p < .001) reflecting overall stronger activation for the language task compared to the picture task across both hemispheres (t(42) = 8.90, p =, .001), and 3) a significant hemisphere x modality interaction (F(42) = 103.37, p < .001) reflecting greater language activation in the left-hemisphere compared to right-hemisphere (t42) = 6.31, p < .001), and greater picture activation in the right-hemisphere compared to the left-hemisphere (t42) = 5.00, p < .001).

### Ventral ATL vs. Lateral Temporal ROIs

In order to examine whether the response in vATL differed to that of the lateral temporal polar region, the coherent vs. scrambled ROIs analyses were conducted using the same bilateral vATL and superior temporal pole ROIs as used in Section 2 (see Figure 3). A 2 x 2 ANOVA comparing the size of the coherence effect for both tasks in vATL and superior-lateral temporal pole revealed a significant coherence effect x ROI interaction in both the left-hemisphere (F(42) = 14.36, p < .001) and right-hemisphere (F(42) = 10.32, p < .005). Follow-up t-test showed that the temporal pole showed a significant coherence effect for both the language and picture tasks, bilaterally (all ts > = 6.12 p < .001). In contrast, the vATL was found to respond equally strongly to the coherent and scrambled conditions in both the language and picture tasks, in both hemispheres (all ts < .84, all ps = .1). Together this supports the earlier finding that within the ATL, the superior-lateral temporal appears to be more selectively recruited for combinatorial process, whereas the vATL responds equally to all forms of semantic stimuli task.

### Experiment 2

The primary aim of fMRI Experiment 2 was to clarify the relationship between the ATL-STS-TPJ network and ToM processing. Specifically, we investigated to what extent can the increased engagement of the ATL-STS-TPJ network by ToM paradigms be explained by the increased combinatorial demands inherent in ToM tasks. This was achieved by systematically varying computational combinatorial load of each item using large language model–derived measures of semantic combinatorial demand alongside traditional psycholinguistic metrics.

**Figure 3.**
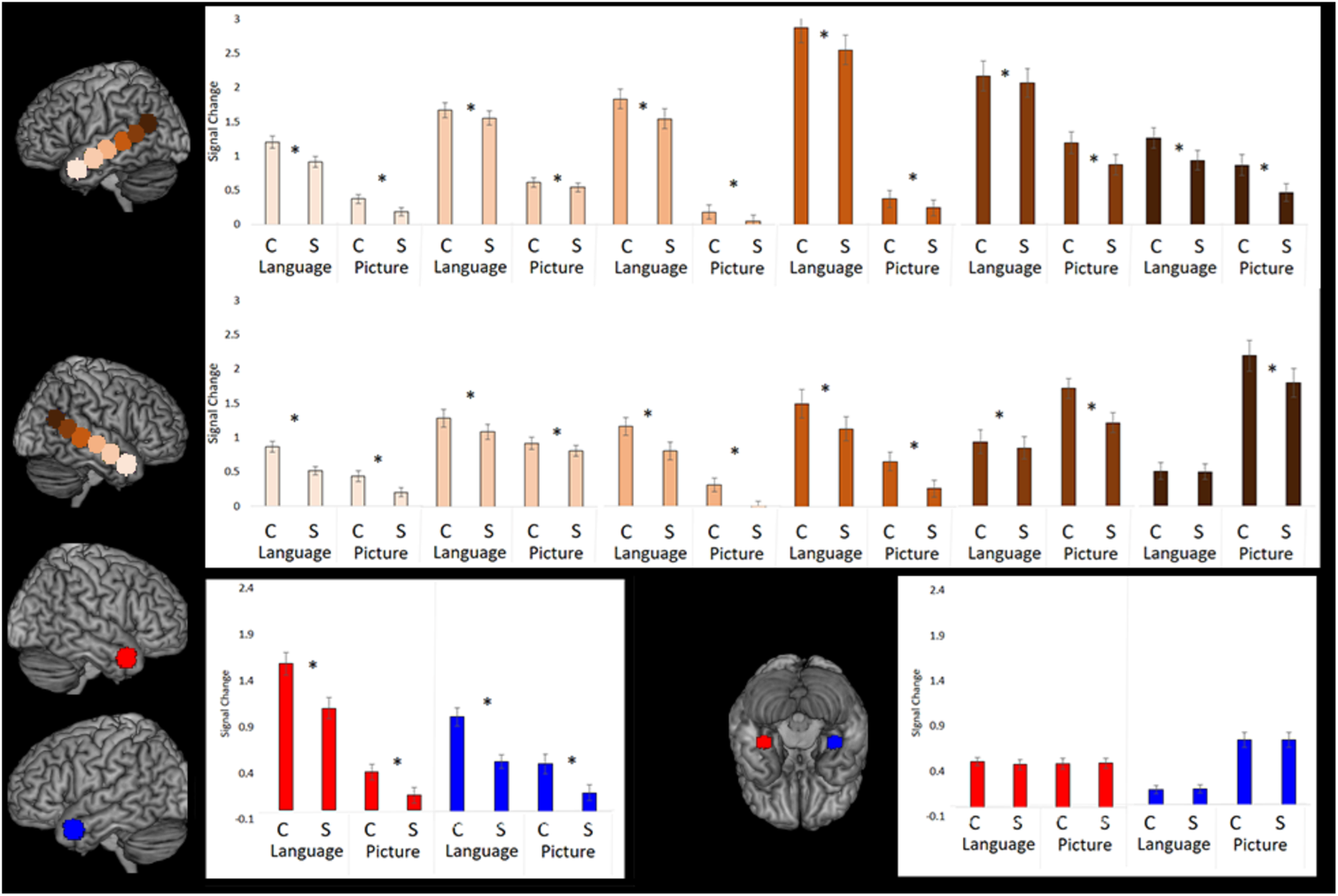
Experiment 1. ROI results for the coherent and scrambled conditions from the language and picture task.

#### ToM and nToM task activation

This experiment compared Theory of Mind (ToM) vs. non-ToM written narratives pooled from two previous studies which found a robust ToM > non-ToM effect (Deen et al., 2015; Dodell-Feder et al., 2011). At the individual level, each task condition was modelled separately against rest, as well as the direct contrast between ToM > nToM items. The contrasts were then analysed at the group-level using one sample t-tests. Group analyses were conducted using standard voxel height threshold p < .001, cluster corrected using FWE p < .05. Compared to rest, the ToM narratives and nToM narratives activated an entirely overlapping network of regions, nearly identical to that engaged by the coherent narratives in study 1. That is a fully bilateral fronto-temporo-parietal network, as well as visual processing areas. Specifically, in the temporal lobe, activation spanned the length of the ATL-STS-TPJ, including the entirety of the temporal pole, as well as the posterior-anterior fusiform gyrus. Frontal activation included lateral frontal cortex (IFG, middle frontal, premotor cortex) as well as medial frontal areas (SMA, vMPFC, dMPFC, orbito-frontal cortex), and parietal activation included SMG, anterior AG, as well IPS. Despite the overlap across conditions, the ToM > nToM contrast revealed that the ToM task elicited significantly stronger activation throughout the entirety of the fronto-temporo-parietal network (with the exception of primary motor cortex, dorsal parietal areas, and visual cortex which did not differ between conditions). The bilateral precuneus was the only area to be revealed by the ToM > nToM contrast that was not present when contrasting task > rest (See Figure 4).

**Figure 4.**
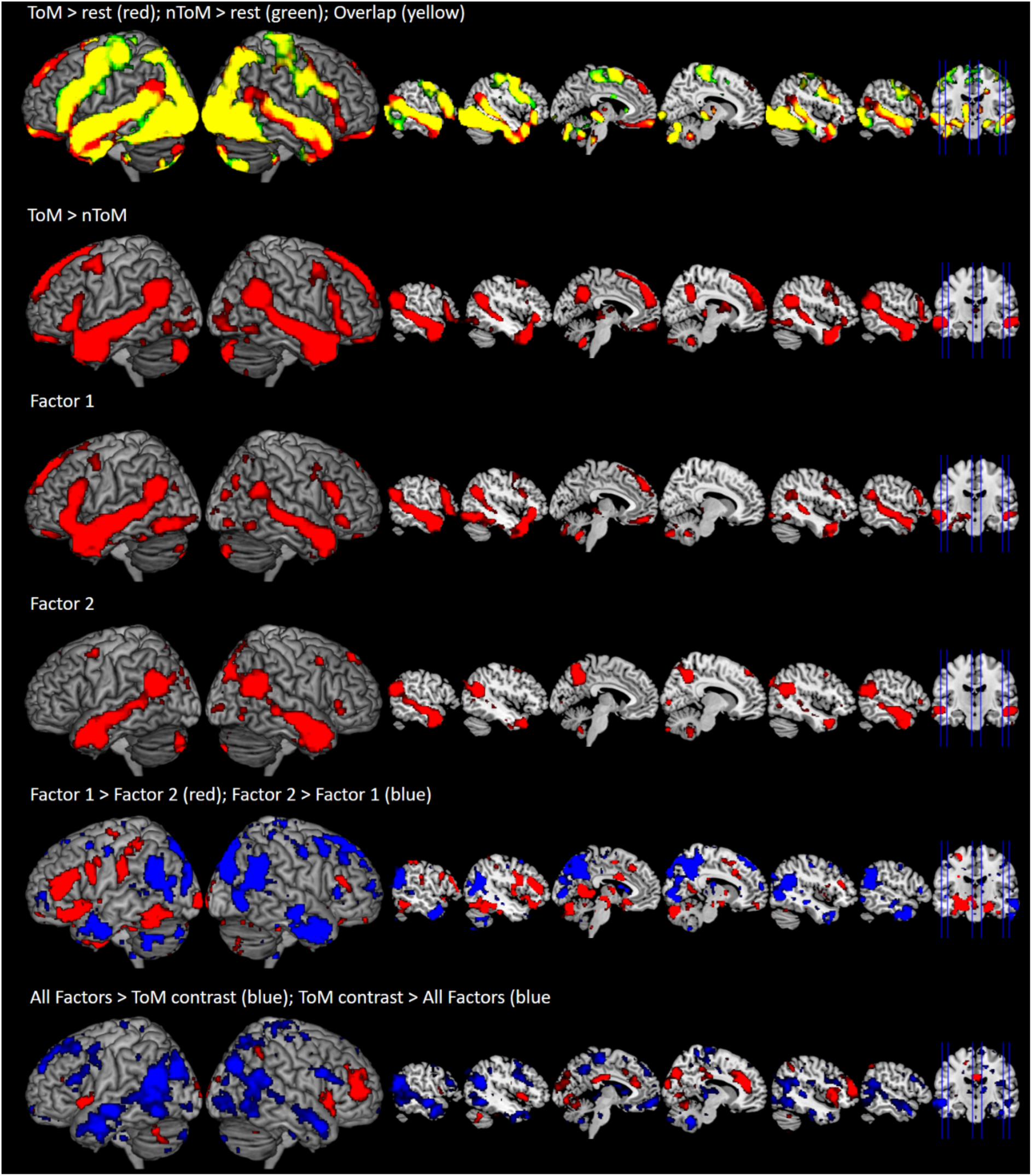
fMRI Experiment 2. Whole-brain fMRI results. Group-level analyses employed a voxel-wise threshold of p < .001, with cluster-level family-wise error correction at p < .05

#### Regression analyses

To determine the extent to which the ToM effect could be explained by linguistic and semantic factors rather than ToM per se, we explored a number of psycholinguistic properties that may influence combinatorial processing, and determined their influence on the ATL-STS-TPJ network and the ToM effect. To do so, for each item we collected a large number of lexical and syntactic probabilistic characteristics that have been shown to influence processing difficulty (Deen et al., 2015) (see Supplementary 3 for full details). In addition to these linguistic variables, the large language Sentence-BERT model (SBERT; model all-mpnet-base-v2; instructions: from https://sbert.net/index.html and https://huggingface.co/sentence-transformers/all-mpnet-base-v2) was used to derive a semantic predictability metric for each narrative stimulus using a model that measures the semantic similarity across the sentences in a narrative. The SBERT- cosine similarity metric provides a computational proxy for the ease of semantic integration within narrative contexts, distinguishing between sequences that support linear, additive integration and those that require more complex, cumulative processing. A principal component analysis with varimax rotation was performed to reduce the large set of correlated variables into a smaller number of underlying factors. This approach addresses multicollinearity among variables and summarises shared variance into orthogonal, interpretable components. Factors with eigenvalues greater than one were retained, yielding four factors that together explained 76.01% of the variance (Table 2 shows the factors and loading scores for each variable). These factors were labelled as: 1) lexico-syntactic complexity (31.81% variance explained), 2) semantic predictability (20.70% variance explained). Factors 3 and 4 were labelled as working-memory-related factors (12.57% and 10.99% variance explained, respectively).

**Table 2.**
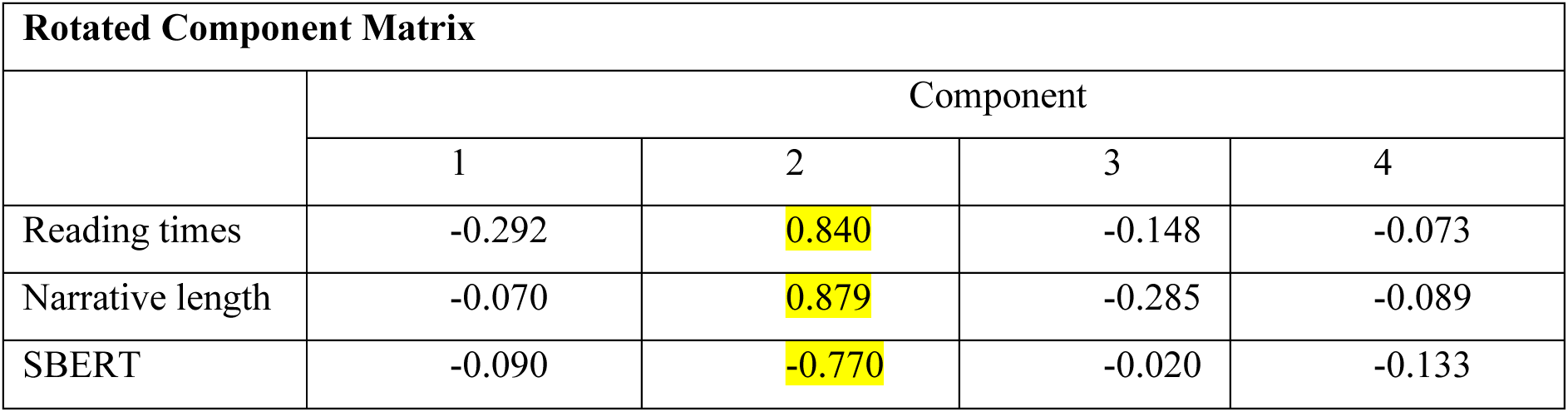
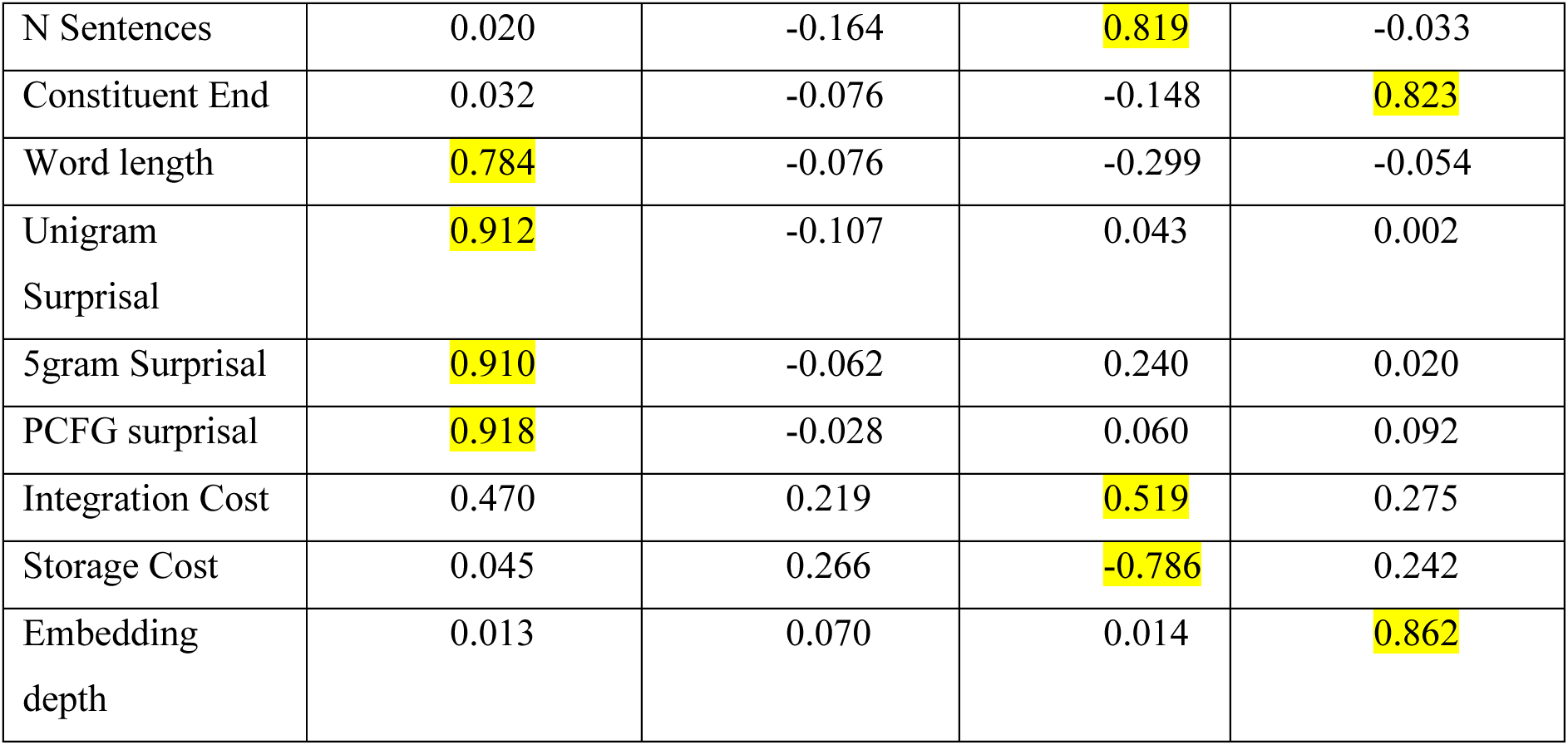
PCA factor loadings.

### Whole-brain results

To determine whether the Theory of Mind (ToM) effect (ToM > nToM) could be explained by the four linguistic and semantic factors rather than ToM per se, the GLM was repeated including the ToM contrast alongside the four factors as covariates. Group-level analyses employed a voxel-wise threshold of p < .001, with cluster-level family-wise error correction at p < .05, See Figure 4.

Lexico-syntactic complexity (Factor 1): Factor 1 was associated with activation spanned the length of the ATL-STS-TPJ, including the entirety of the temporal pole, as well as the posterior-anterior fusiform gyrus. Frontal activation included lateral frontal cortex (IFG, middle frontal, premotor cortex) as well as medial frontal areas (SMA, vMPFC, dMPFC, orbito-frontal cortex). This pattern directly mirrored that of the ToM > nToM contrast, with the exception of the precuneus which was not significantly modulated by Factor 1.

Semantic predictability (Factor 2): Factor 2 was specifically associated with activation along the length of the ATL-STS-TPJ network bilaterally, as well as the bilateral precuneus. Factor 3 and 4 were not significantly associated with any neural activation.

Direct contrasts of Factor 2 > 1 showed that activation of the bilateral anterior temporal lobe, TPJ, and precuneus was more strongly related to the semantic predictability compared to lexico-syntactic complexity of the stimuli, whereas lexico-syntactic complexity (Factor 1 > 2) was more strongly related to activation in left lateral prefrontal cortex, as well as left posterior fusiform gyrus.

All Factors > ToM contrast: In order to determine to what extent task activation could be explained by the PCA factors over and above the ToM > contrast we directly compared the activation for all Factors combined with the ToM contrast (ToM > nToM), and vice versa. All Factors combined were associated with significantly greater activation than the ToM contrast within the bilateral entire ATL-STS-TPJ network, as well as bilateral middle frontal gyrus, and midline areas within anterior medial prefrontal cortex and precuneus. Nevertheless, not all ToM contrast activation could be explained by the PCA Factors. Indeed, the ToM contrast revealed significantly greater activation compared to the combined Factors within bilateral insular and orbitofrontal cortex, bilateral anterior cingulate, and right frontal pole.

#### Summary

To summarise the results from fMRI Experiment 2, 1) The ToM and nToM narrative engaged a common network of fully bilateral fronto-temporo-parietal network, similar to that found in Experiment 1. Specifically, in the temporal lobe, activation spanned the length of the ATL-STS-TPJ, including the entirety of the temporal pole, as well as the posterior-anterior fusiform gyrus. Frontal activation included lateral frontal cortex as well as medial frontal areas. 2) Despite the overlapping engagement, as expected the ToM narratives more strongly engaged a fronto-temporal network including the entire bilateral ATL-STS-TPJ network, as well as lateral and medial frontal areas. 3) Significantly, however, this effect was almost entirely eliminated when controlling for lexico-syntactic, and the semantic coherence of the stimuli (with the exception of a few small clusters including in the bilateral insular, orbito-frontal cortex, and anterior cingulate). 4) Interestingly, a direct comparison on the lexico-syntactic factor (Factor 1) and the semantic coherence factor (Factor 2) revealed that semantic coherence more strongly modulated activation of the bilateral temporal pole and TPJ, whereas lexico-syntactic properties more strongly engaged the left lateral frontal and posterior MTG, a network previously linked to executive processing in semantic tasks (the semantic control network).

## Discussion

The overarching aim of the present study was to delineate the neural architecture supporting combinatorial semantic processing and its relationship to single-concept semantics and ToM processing with the goal of uniting thee related yet separate research literatures. Specifically, we aimed to 1) determine the extent to which combinatorial semantics engages a stable and reliable neural network, and if so, 2) to what extent this is similar or different to that engaged by single-concept semantics, particularly with regard ATL engagement. 3) The extent to which the combinatorial semantic processing network is transmodal in nature, and 4) how do the neurocomputations that underpin combinatorial semantic processing interact with ToM processing, for instance, are the networks overlapping or functionally adjacent, and 5) to what extent any apparent overlap can be explained by the combinatorial demands inherent in ToM tasks. These five aims were investigated using a combination of three different methodological approaches. Establishing convergent results across these very different methods provides powerful evidence for stable and reliable findings.

First, we found highly convergent evidence across the three approaches to suggest that the bilateral ATL-STS-TPJ is a stable and reliable network engaged by combinatorial semantics (Aim 1). Specifically, 1) the ALE meta-analysis (Section 1) showed that this network is more reliably engaged by studies involving the semantic judgments based on sentences compared to single words, and 2) the omnibus fMRI study results (Section 2) demonstrated that the same network is more engaged by tasks involving semantic sequences compared non-semantic sequences or single-item semantic judgements. Finally, 3) the two targeted and well-controlled within-subject fMRI experiments that directly manipulated combinatorial processing (section 3) revealed that same ATL-STS-TPJ network is multimodal in nature and responds more strongly to coherent vs. scrambled narratives in both the language and picture domain (Aim 3).

In terms of Aim 2: given the prominence of the ATL to single concept semantics, what role does the ATL play in combinatorial semantic processing? Across studies, the results suggested that 1) the ATL is engaged by both single-concept and combinatorial semantics, but 2) there are graded differences in the response across the ATL whereby the vATL is equally engaged by all semantic tasks showing equal engagement for by single concept semantics and combinatorial semantics in the omnibus fMRI study, and the intact combinatorial tasks as compared to the scrambled items (Section 3 - fMRI Study 1). In contrast to the response from the vATL which responded equally to all semantic tasks, the lateral superior ATL appears to be more selectively involved in combinatorial semantic tasks showing i) a greater response to tasks that engage combinatorial semantic processing as compared to single concept semantics which did not differ from rest, and ii) a stronger response to the intact combinatorial tasks as compared to the scrambled items (Section 3 - fMRI Study 1). Together these results suggest a more general role of the vATL in all forms of semantic cognition, whereas the lateral superior ATL may only become engaged for tasks that require some form of combinatorial semantic integration.

With regard Aims 3 and 4: the findings from the meta-analysis and omnibus fMRI study showed that the same overlapping network appears to be engaged by verbal and nonverbal ToM tasks, as well as tasks involving social stimuli. There are at least two possible explanations for this effect 1) the results could be explained by a social confound in the stimuli: Indeed, many combinatorial semantic studies use human- or socially-themed material, which could bias results towards overlap with ToM regions. Or 2) ToM tasks may inherently place greater demands on the combinatorial semantic system, as they require the continuous integration of semantic information over time to understand complex, inferential social events. Therefore, to address Aim 5; the former possibility was firmly rejected by the findings from fMRI Study 2 (Section 3) which demonstrated that the ToM effect in the ATL-STS-TPJ network could be eliminated entirely when controlling for the lexico-syntactic features and semantic coherence of the stimuli. Overall, the results provide clear and convergent evidence for the extensive of a multimodal combinatorial network in the bilateral ATL-STS-TPJ.

### A unified neurocomputational framework for single-concept and combinatorial semantics

The hub-and-spoke model provides a mechanistic account of how conceptual knowledge is both distributed and integrated within the semantic system (Lambon Ralph et al., 2017). In this framework, modality-specific cortical regions encode sensory-motor and linguistic features of concepts, while a central, transmodal hub located in the bilateral ATL integrates these inputs into coherent conceptual representations. Two properties are central to this account. First, the hub is transmodal, allowing information derived from different modalities to be unified within a single representational space. Second, it is transtemporal, such that conceptual knowledge is shaped by accumulated experience over time. This allows for the formation of stable conceptual representations that are robust to varying context. Computational instantiations of the hub-and-spoke architecture demonstrate that these properties are sufficient to support time-extended abstraction, in which multiple features of a concept can be learned and maintained without requiring their simultaneous activation (Jackson et al., 2021).

Our fMRI findings provide direct neural support for this unified account. We show that combinatorial semantic processing engages the same ATL regions for both verbal and nonverbal tasks, overlapping with areas implicated in single-concept semantic representation. In terms of computational models, the same computational principles that support single-concept semantics also provide a natural basis for combinatorial semantic processing (Hoffman et al., 2018). Rather than treating combinatorial semantics as a qualitatively distinct process, this computational model showed how temporal integration can arise naturally from the same hub-based architecture whereby a model takes a sequence of single words as input and learns by predicting both the upcoming words and each word’s sensory-motor properties. Through this learning process, the model acquires context-sensitive semantic representations that reflect the multimodal experiential knowledge for each concept while also accounting for co-occurrence and information integration over time (McClelland, St. John, et al., 1989), and the same architecture has shown to operate for nonverbal semantics (pictures of objects in a context) (Sadeghi et al., 2015). In this way, single-concept semantics and combinatorial semantics are explained by a common neural and computational mechanism centred on ATL hub.

### A Graded Transmodal Semantic Network

We found that the combinatorial semantic processing network was entirely overlapping for the picture and language tasks, consistent with a transmodal hub that integrates information across different modalities. Within this largely overlapping network, there were subtle asymmetries. Language stimuli produced stronger activation in the left hemisphere, particularly nearby auditory cortex, while picture stimuli produced slightly stronger activation in the right hemisphere, especially in posterior occipito-parietal regions. Correspondingly, the magnitude of the coherence effect was larger for language in auditory-related areas and larger for pictures in visual-spatial regions. This is consistent with the hub-and-spoke whereby graded modality biases arise from differential connectivity to modality-specific spokes (Lambon Ralph et al., 2017). Regions of the temporal lobe more strongly connected to auditory and language areas are expected to respond more to language stimuli, whereas regions with stronger connections to visual and spatial areas respond more to pictures. This provides a natural explanation for the subtle asymmetries we observed: left-hemisphere language dominance near auditory cortex, right-hemisphere picture dominance in posterior occipito-parietal regions.

### A Graded ATL hub

With regard functional importance of the ATL our results go one step-further by revealing graded functional differentiation within the ATL. Ventral bilateral ATL were engaged across all semantic tasks, irrespective of whether they involved single concepts or combinatorial meaning, consistent with its proposed role as the core transmodal semantic hub. In contrast, bilateral superior lateral ATL around the polar region were additionally recruited by tasks that required the integration of multiple semantic elements into a coherent gestalt or the drawing of associative semantic inferences. This finding was consistent across approaches in Section 2 and Section3, and aligns with the proposal that sequential information is continually integrated into a semantic gestalt by a ventral pathway terminating at the ATL. Connectionist models of sentence comprehension demonstrate that sequential inputs are incrementally and continuously integrated over time (McClelland, St. John M, et al., 1989).

In contrast to the lateral ATL, the vATL appears to be commonly engaged by all forms of semantic task. It may be that vATL is universally active because it occupies the centre of the semantic representational machinery and receives convergent input from all semantic sources, whereas the dorsal-lateral ATL is more biased towards auditory and verbal materials, which are inherently time-varying stimuli (Visser & Lambon Ralph, 2011). Additionally, due to the vATLs falling at the apex of the ventral pathway, it may be that the vATL region marries temporally-varying information from the STS/TJP network with more discretely segmented and/or spatially bounded individuated representations, such as those experienced during visual object processing. As shown computationally by Hoffman et al (2018), this merger would support the formation of stable, context-general conceptual representations, consistent with ventral ATL engagement across all semantic conditions.

The finding that combinatorial semantic processing especially engages the temporal pole has important implications for research on semantic dementia, a condition in which this region is severely atrophied. While it is well established that semantic dementia patients show profound multimodal semantic deficits (Jefferies & Lambon Ralph, 2006), the current findings imply that the patients might also be particularly impaired in combinatorial or sequential meaning processing. Exploring this hypothesis could reveal new insights into the nature of semantic deficits in these patients and help distinguish between general loss of semantic knowledge and impaired semantic integration. Furthermore, this perspective could guide the development of targeted behavioural assessments and interventions aimed at probing and supporting combinatorial semantic processing, offering a more fine-grained understanding of semantic dementia.

### The ATL-STS-TPJ network is modulated by combinatorial semantics rather than Theory of Mind processes

Our finding from Section 3 fMRI Study 2 showed that the apparent overlap between combinatorial semantic processing and the Theory of Mind (ToM) network is eliminated when controlling for linguistic and semantic integration demand provides direct evidence supporting the view that previous reports of convergence may reflect task- or content-driven confounds rather than a shared cognitive mechanism. While meta-analyses have suggested anatomical overlap in regions such as the STS–TPJ (Schurz et al., 2021), and classical theory predicts that ToM draws on semantic memory to interpret social cues and construct coherent event representations (C. D. Frith & Frith, 2021), our results indicate that this engagement is not intrinsic to ToM *per se*. Instead, combinatorial semantic activation within ToM-associated regions appears to be modulated by the language and semantic integration requirements of the stimuli. This aligns with prior work showing that ToM effects within language networks diminish when linguistic variables are controlled (Shain et al., 2023) and emphasises the importance of distinguishing domain-general semantic integration from domain-specific social cognition.

The results from Section 3 Experiments 1 and 2 also highlight an additional interesting contrast: we found that activation within the combinatorial semantic network was parametrically modulated by increasing demands on semantic integration systems (fMRI Experiment 2), yet reduced activation when the input is entirely scrambled and cannot be integrated at all (fMRI Experiment 1). Clearly, this result is incompatible with a simple “combinatorial effort” account, in which greater difficulty should monotonically increase neural response. Instead, it seems like that with scrambled materials the semantic combinatorial mechanisms fail entirely and instead the multi-demand executive system is engaged, presumably to initiate more general executive mechanisms for solving novel problems.

Although controlling for linguistic and semantic demand eliminated the apparent overlap between combinatorial semantic processing and the canonical ToM network, we found that ToM selectively modulated activation in bilateral insular and orbitofrontal cortex, bilateral anterior cingulate, and the right frontal pole. These regions have been repeatedly implicated in social cognition and mental state inference independent of language. The insula and anterior cingulate cortex are core components of the brain’s salience and social evaluative systems, contributing to emotion processing, empathy, and the evaluation of others’ motivations and social outcomes, functions that underpin ToM and broader social reasoning (Apps et al., 2016). The orbitofrontal cortex has been associated with social evaluation, value representation in social decision-making, and adjusting behaviour based on social context, with lesions or atrophy here producing deficits in adaptive social behaviour (Rouse et al., 2024). These regions are also prominently affected in behavioural variant frontotemporal dementia (bvFTD), where atrophy of orbitofrontal, medial frontal, insula, and cingulate cortices is linked to profound impairments in ToM, social conduct, and social context processing. Together, this pattern supports the view that ToM relies on a distinct fronto-insula social-cognitive network specialised for evaluating others’ internal states and guiding socially appropriate behaviour, separate from the transmodal semantic integration network, and that perturbations of these regions in clinical populations underpins characteristic social cognitive deficits.

## Notes

### Competing Interest Statement

The authors have declared no competing interest.

## References

Apps, M. A. J., Rushworth, M. F. S., & Chang, S. W. C. (2016). The Anterior Cingulate Gyrus and Social Cognition: Tracking the Motivation of Others. Neuron, 90(4), 692–707. 10.1016/j.neuron.2016.04.018

Balgova, E., Diveica, V., Jackson, R. L., & Binney, R. J. (2024). Overlapping neural correlates underpin theory of mind and semantic cognition: Evidence from a meta-analysis of 344 functional neuroimaging studies. Neuropsychologia, 200, 108904. 10.1016/j.neuropsychologia.2024.108904

Binney, R. J., Embleton, K. V., Jefferies, E., Parker, G. J. M., & Lambon Ralph, M. A. (2010). The Ventral and Inferolateral Aspects of the Anterior Temporal Lobe Are Crucial in Semantic Memory: Evidence from a Novel Direct Comparison of Distortion-Corrected fMRI, rTMS, and Semantic Dementia. Cerebral Cortex, 20(11), 2728–2738. 10.1093/cercor/bhq019

Deen, B., Koldewyn, K., Kanwisher, N., & Saxe, R. (2015). Functional Organization of Social Perception and Cognition in the Superior Temporal Sulcus. Cerebral Cortex, 25(11), 4596–4609. 10.1093/cercor/bhv111

Dodell-Feder, D., Koster-Hale, J., Bedny, M., & Saxe, R. (2011). fMRI item analysis in a theory of mind task. NeuroImage, 55(2), 705–712. 10.1016/j.neuroimage.2010.12.040

Frith, C. D., & Frith, U. (2021). Mapping Mentalising in the Brain. In M. Gilead & K. N. Ochsner (Eds), The Neural Basis of Mentalizing (pp. 17–45). Springer International Publishing. 10.1007/978-3-030-51890-5_2

Frith, U., & Happé, F. (1994). Autism: Beyond “theory of mind”. Cognition, 50(1), 115–132. 10.1016/0010-0277(94)90024-8

Halai, A. D., Henson, R. N., Finoia, P., & Correia, M. M. (2025). Comparing the effect of multi-gradient echo and multi-band fMRI during a semantic task. Imaging Neuroscience, 3, IMAG.a.1043. 10.1162/IMAG.a.1043

Halai, A. D., Welbourne, S. R., Embleton, K., & Parkes, L. M. (2014). A comparison of dual gradient-echo and spin-echo fMRI of the inferior temporal lobe. Human Brain Mapping, 35(8), 4118–4128. 10.1002/hbm.22463

Hasson, U., Yang, E., Vallines, I., Heeger, D. J., & Rubin, N. (2008). A hierarchy of temporal receptive windows in human cortex. J Neurosci, 28(10), 2539–2550. 10.1523/JNEUROSCI.5487-07.2008

Hoffman, P., McClelland, J. L., & Lambon Ralph, M. A. (2018). Concepts, control, and context: A connectionist account of normal and disordered semantic cognition. Psychological Review, 125(3), 293–328. 10.1037/rev0000094

Humphreys, G. F., Hoffman, P., Visser, M., Binney, R. J., & Ralph, M. A. L. (2015). Establishing task- and modality-dependent dissociations between the semantic and default mode networks. Proceedings of the National Academy of Sciences of the United States of America, 112(25), 7857–7862. 10.1073/pnas.1422760112

Humphreys, G. F., & Lambon Ralph, M. A. (2015). Fusion and fission of cognitive functions in the human parietal cortex. Cerebral Cortex, 25, 3547–3560.

Humphreys, G. F., Lambon Ralph, M. A., & Simons, J. S. (2021). A Unifying Account of Angular Gyrus Contributions to Episodic and Semantic Cognition. Trends in Neurosciences, 44(6), 452–463. 10.1016/j.tins.2021.01.006

Jackson, R. L., Rogers, T. T., & Lambon Ralph, M. A. (2021). Reverse-engineering the cortical architecture for controlled semantic cognition. Nature Human Behaviour, 5(6), 774–786. 10.1038/s41562-020-01034-z

Jefferies, E., & Lambon Ralph, M. A. (2006). Semantic impairment in stroke aphasia versus semantic dementia: A case-series comparison. Brain, 129, 2132–2147. Doi 10.1093/Brain/Awl153

Lambon Ralph, M. A., Jefferies, E., Patterson, K., & Rogers, T. T. (2017). The neural and computational bases of semantic cognition. Nat Rev Neurosci, 18(1), 42–55. 10.1038/nrn.2016.150

Lambon Ralph, M. A., Sage, K., Jones, R. W., & Mayberry, E. J. (2010). Coherent concepts are computed in the anterior temporal lobes. Proceedings of the National Academy of Sciences of the United States of America, 107(6), 2717–2722. DOI 10.1073/pnas.0907307107

Lerner, Y., Honey, C. J., Silbert, L. J., & Hasson, U. (2011). Topographic Mapping of a Hierarchy of Temporal Receptive Windows Using a Narrated Story. Journal of Neuroscience, 31(8), 2906–2915. 10.1523/JNEUROSCI.3684-10.2011

McClelland, J. L., St. John, M., & Taraban, R. (1989). Sentence comprehension: A parallel distributed processing approach. Language and Cognitive Processes, 4(3–4), SI287–SI335. 10.1080/01690968908406371

McClelland, J. L., St. John M, & Taraban, R. (1989). Sentence comprehension: A parallel distributed processing approach. Language and Cognitive Processes, 4, 287–335.

Olson, I. R., McCoy, D., Klobusicky, E., & Ross, L. A. (2013). Social cognition and the anterior temporal lobes: A review and theoretical framework. Social Cognitive and Affective Neuroscience, 8(2), 123–133. 10.1093/scan/nss119

Pobric, G., Jefferies, E., & Ralph, M. A. L. (2007). Anterior temporal lobes mediate semantic representation: Mimicking semantic dementia by using rTMS in normal participants. Proceedings of the National Academy of Sciences, 104(50), 20137–20141. 10.1073/pnas.0707383104

Rogers, T. T., Cox, C. R., Lu, Q., Shimotake, A., Kikuchi, T., Kunieda, T., Miyamoto, S., Takahashi, R., Ikeda, A., Matsumoto, R., & Lambon Ralph, M. A. (2021). Evidence for a deep, distributed and dynamic code for animacy in human ventral anterior temporal cortex. eLife, 10, e66276. 10.7554/eLife.66276

Rouse, M. A., Binney, R. J., Patterson, K., Rowe, J. B., & Lambon Ralph, M. A. (2024). A neuroanatomical and cognitive model of impaired social behaviour in frontotemporal dementia. Brain: A Journal of Neurology, 147(6), 1953–1966. 10.1093/brain/awae040

Sadeghi, Z., McClelland, J. L., & Hoffman, P. (2015). You shall know an object by the company it keeps: An investigation of semantic representations derived from object co-occurrence in visual scenes. Special Issue: Semantic Cognition, 76, 52–61. 10.1016/j.neuropsychologia.2014.08.031

Sato, N., Matsumoto, R., Shimotake, A., Matsuhashi, M., Otani, M., Kikuchi, T., Kunieda, T., Mizuhara, H., Miyamoto, S., Takahashi, R., & Ikeda, A. (2021). Frequency-Dependent Cortical Interactions during Semantic Processing: An Electrocorticogram Cross-spectrum Analysis Using a Semantic Space Model. Cerebral Cortex, 31(9), 4329–4339. 10.1093/cercor/bhab089

Saxe, R., & Powell, L., J. (2006). It’s the Thought That Counts: Specific Brain Regions for One Component of Theory of Mind. Psychological Science, 17(8), 692–699. 10.1111/j.1467-9280.2006.01768.x

Schurz, M., Radua, J., Tholen, M. G., Maliske, L., Margulies, D. S., Mars, R. B., Sallet, J., & Kanske, P. (2021). Toward a hierarchical model of social cognition: A neuroimaging meta-analysis and integrative review of empathy and theory of mind. Psychological Bulletin, 147(3), 293–327. 10.1037/bul0000303

Shain, C., Paunov, A., Chen, X., Lipkin, B., & Fedorenko, E. (2023). No evidence of theory of mind reasoning in the human language network. Cerebral Cortex, 33(10), 6299–6319. 10.1093/cercor/bhac505

Shimotake, A., Matsumoto, R., Ueno, T., Kunieda, T., Saito, S., Hoffman, P., Kikuchi, T., Fukuyama, H., Miyamoto, S., Takahashi, R., Ikeda, A., & Lambon Ralph, M. A. (2015). Direct Exploration of the Role of the Ventral Anterior Temporal Lobe in Semantic Memory: Cortical Stimulation and Local Field Potential Evidence From Subdural Grid Electrodes. Cerebral Cortex, 25(10), 3802–3817. 10.1093/cercor/bhu262

Sueoka, Y., Paunov, A., Tanner, A., Blank, I. A., Ivanova, A., & Fedorenko, E. (2024). The Language Network Reliably “Tracks” Naturalistic Meaningful Nonverbal Stimuli. Neurobiology of Language, 5(2), 385–408. 10.1162/nol_a_00135

Ueno, T., Saito, S., Rogers, T. T., & Lambon Ralph, M. A. (2011). Lichtheim 2: Synthesizing Aphasia and the Neural Basis of Language in a Neurocomputational Model of the Dual Dorsal-Ventral Language Pathways. Neuron, 72(2), 385–396. DOI 10.1016/j.neuron.2011.09.013

Visser, M., Jefferies, E., & Ralph, M. A. L. (2010). Semantic Processing in the Anterior Temporal Lobes: A Meta-analysis of the Functional Neuroimaging Literature. Journal of Cognitive Neuroscience, 22(6), 1083–1094.

Visser, M., & Lambon Ralph, M. A. (2011). Differential Contributions of Bilateral Ventral Anterior Temporal Lobe and Left Anterior Superior Temporal Gyrus to Semantic Processes. Journal of Cognitive Neuroscience, 23(10), 3121–3131. 10.1162/jocn_a_00007

Zaccarella, E., Schell, M., & Friederici, A. D. (2017). Reviewing the functional basis of the syntactic Merge mechanism for language: A coordinate-based activation likelihood estimation meta-analysis. Neuroscience & Biobehavioral Reviews, 80, 646–656. 10.1016/j.neubiorev.2017.06.011

