## Supplementary Materials for "A lateral temporal network for transmodal combinatorial semantics: Convergent evidence from a multi-study investigation"

**Supplementary 1**

**Section 1: Meta-analysis**

**Methods**

*Study selection:* The meta-analysis comprised 133 PET and fMRI experiments that included a sentence comprehension paradigm (e.g. sentences > word strings (Kuperburg et al., 2000), auditory sentences > speech envelope sound (von Kriegstein et al., 2003), sentences > nonwords (Baumgaertener et al., 2002) etc.), and a 93 experiments that included a semantic tasks using single-word level studies compared to a control condition or resting baseline (e.g. word > nonword (Garbin et al., 2012; Vignali et al., 2019), semantic > phonological judgement (Price et al. 1997), semantic > perceptual judgment (Seghier et al., 2010) etc.) (see Table S1 for a list of included studies). In addition we included 113 experiments that involved a theory of mind task of some form, 71 from the verbal domain (e.g. false belief narrative > false photo narrative (Aichhorn et al., 2008), ToM narrative > non-social narrative (Cheung et al., 2012) etc), and 42 experiments from the nonverbal domain (e.g. shape animations: social judgment > visuospatial judgment (Martin & Weisberg 2003; McAdams & Krawczyk, 2011; cartoons with ToM > non-ToM decisions (Roser et al., 2012; Saft et al., 2013; Samson et al., 2008). Our inclusion criteria were any fMRI/PET study that reported peak activation in standard space (Talairach or MNI) based on whole-brain statistical comparisons. Studies were selected from existing comprehensive meta-analyses (Hodgson et al., 2023; Jackson, 2021).

*ALE analyses*: For each domain (sentence processing, word processing, ToM processing) ALE analyses were carried out using GingerAle 3.0.2 (Eickhoff et al., 2009;(Humphreys et al., 2024) Laird et al., 2005). All activation peaks were converted to MNI standard space using the built-in Talairach to MNI (SPM) GingerALE toolbox. Analyses were performed with voxel-level thresholding at a p-value of 0.001 and cluster-level FWE-correction with a p-value of 0.05 over 10,000 permutations. To directly compare ALE maps for 1) sentence-level and word-level studies, and 2) verbal vs. non-verbal ToM tasks we conducted contrast analyses following the procedure of Laird et al. (2005). In this approach, experiments from both domains are combined and repeatedly randomized to generate a null distribution of ALE score differences, against which a Z-value map of the observed difference is computed. For the present analyses, 10,000 permutations were performed, and an uncorrected threshold of $P < 0.001$ was applied.

*Study selection:* The meta-analysis comprised 133 PET and fMRI experiments that included a sentence comprehension paradigm (e.g. sentences > word strings (Kuperburg et al., 2000), auditory sentences > speech envelope sound (von Kriegstein et al., 2003), sentences > nonwords (Baumgaertener et al., 2002) etc.), and a 93 experiments that included a semantic tasks using single-word level studies compared to a control condition or resting baseline (e.g. word > nonword (Garbin et al., 2012; Vignali et al., 2019), semantic > phonological judgement (Price et al. 1997), semantic > perceptual judgment (Seghier et al., 2010) etc.) (see Table S1 for a list of included studies). In addition we included 113 experiments that involved a theory of mind task of some form, 71 from the verbal domain (e.g. false belief narrative > false photo narrative (Aichhorn et al., 2008), ToM narrative > non-social narrative (Cheung et al., 2012) etc), and 42 experiments from the nonverbal domain (e.g. shape animations: social judgment > visuospatial judgment (Martin & Weisberg 2003; McAdams & Krawczyk, 2011; cartoons with ToM > non-ToM decisions (Roser et al., 2012; Saft et al., 2013; Samson et al., 2008). Our inclusion criteria were any fMRI/PET study that reported peak activation in standard space (Talairach or MNI) based on whole-brain statistical comparisons. Studies were selected from existing comprehensive meta-analyses (Hodgson et al., 2023; Jackson, 2021).

*ALE analyses*: For each domain (sentence processing, word processing, ToM processing) ALE analyses were carried out using GingerAle 3.0.2 (Eickhoff et al., 2009;(Humphreys et al., 2024) Laird et al., 2005). All activation peaks were converted to MNI standard space using the built-in Talairach to MNI (SPM) GingerALE toolbox. Analyses were performed with voxel-level thresholding at a p-value of 0.001 and cluster-level FWE-correction with a p-value of 0.05 over 10,000 permutations. To directly compare ALE maps for 1) sentence-level and word-level studies, and 2) verbal vs. non-verbal ToM tasks we conducted contrast analyses following the procedure of Laird et al. (2005). In this approach, experiments from both domains are combined and repeatedly randomized to generate a null distribution of ALE score differences, against which a Z-value map of the observed difference is computed. For the present analyses, 10,000 permutations were performed, and an uncorrected threshold of $P < 0.001$ was applied.

### Results

*Sentence tasks and sing-word tasks:* The results from the ALE analysis of sentence studies revealed a fronto-temporo-parietal network. Reliable activation was found to extend the length of the superior and middle temporal gyrus, from the anterior temporal pole until the temporo-parietal junction (TPJ). Recruitment was bilateral, although extending more posteriorly in the left hemisphere. Within frontal cortex, sentences reliably engaged left frontal areas within precentral gyrus, inferior frontal gyrus (IFG, BA44, BA45), and supplementary motor area. In contrast, single word studies were found to reliably elicit a more restrictive network, and was limited to the left hemisphere. This included clusters with IFG, premotor cortex, posterior middle temporal gyrus, and angular gyrus, but not the more anterior portion of the temporal lobe or the TPJ. The two networks overlapped in LIFG, pMTG, and the anterior potion of the AG. Contrast analyses revealed that sentence studies were statistically more likely than single word studies to recruit the length of the left superior and middle temporal gyrus, the right anterior superior and middle temporal gyrus, and the LIFG. In order to check that this difference was driven by there being a greater number of auditory compared to visual studies for sentences compared to single words the analysis was repeated using only Sentence tasks from the visual domain. The visual-only sentence network was found to be largely identical to the all-sentence network, although slightly more restrictive in the right hemisphere, thereby suggesting the sentence vs. word differences are not driven by modality.

*Sentence tasks and Theory of mind tasks:* There was found to be striking overlap in the network recruited by ToM and Sentence tasks, particularly along the length of left STG/MTG/TPJ as well as the LIFG. In the right hemisphere, the two tasks overlapped across the right STG/MTG, but the right TPJ/AG and RIFG were observed only in the ToM tasks. Contrast analyses revealed that despite the overlap Sentence tasks were more likely to recruit the left STG/MTG, as well as LIFG, whereas the right TPJ/AG were more reliably recruited by ToM tasks. The ToM tasks included in the analysis were a combination of task paradigms, some utilising linguistic tasks where the participants comprehend verbal stories, and others using abstract shape cartoons. Thus, it is possible that the high degree of Sentence –ToM overlap could be driven by the inclusion linguistic task in the ToM analysis. To test this, we repeated the analysis but this time separating the ToM tasks into those that used linguistic stimuli compared to those that used non-verbal cartoons. The resultant ALE maps were found to be largely overlapping across modalities, with no statistically significant difference between the two. Therefore, the Sentence –ToM overlap cannot be explained by the inclusion of verbal ToM tasks.

**Supplementary 2**

**Section 2: fMRI omnibus study**

**Methods**

*Task design and procedure*

For the fMRI omnibus study we combined the data from eight fMRI studies, totalling 17 task paradigms, and X participants. Below is a brief description of each study (for a detailed methodological description for each study please refer to the original publication).

1) Humphreys et al. (2024): A receptive speech production task whereby participants were given a cue topic followed by a prompt in which they verbal described their response to that topic for 14 seconds. and asked to speak about the Autobiographical condition. There were three conditions: autobiographical event description, semantic event description, and a control condition (recite the months of the year). In the autobiographical event description condition the participants were presented with a variety of questions probing autobiographical memories of events (e.g., describe the last time you had a cup of tea). In the semantic event description condition, the probe questions had a similar content to the autobiographical condition but were phrased to refer to a generalised semantic event rather than a personal autobiographical experience (e.g., describe how you would typically make a cup of tea). For the control condition, the participants were asked to recite the months of the year, repeating if necessary for the full 14 seconds.

2) Branzi et al. 2020: This involved a semantic comprehension task and numerical sequence task. In the semantic comprehension task, a narrative comprehension condition whereby participants read narratives describing events that included human participants. In the numerical sequence task, the participants read strings of 4-digit numbers that followed a simple numerical pattern.

3) Branzi et al (in prep): There were two conditions of interest, a narrative comprehension condition whereby the participants listened to short auditory narrative describing human event, and a music condition whereby the participants listened to unfamiliar nonverbal classical music.

4) Humphreys et al. (2020): The participants completed three experimental tasks: sentence task, picture task, and number task. The sentences described events some involving human participants and some involving non-human participants (e.g. *He loosened the tie around his neck;* or *The cow was on the farm*). Picture task*:* a series of four colour pictures depicted the occurrence of real-life nonhuman everyday events with a clear causal structure, that is, the events could not plausibly occur in a different order (e.g., a banana being peeled, a house being built, etc.). Number task: a series of four numbers involving low-digit multiplication (e.g., 2 4 6 8) or addition (e.g., 1 2 3 4).

5) (Humphreys et al., 2022): There were 3 conditions of interest: object episodic retrieval; object semantic retrieval, and a control task. In the object episodic retrieval task, the participants selected the feature that best matched the target items. The target items were verbal labels that corresponded to colour photographs that were viewed prior to the scan (e.g. bucket: green or red). In

the object semantic task, the participants indicated which item was semantically related to the target (e.g. dart; sharp or soft). The control task involved a visual decision as to whether a scrambled image had moved to the left or right of fixation.

6) (Humphreys & Lambon Ralph, 2025): This consisted of a picture-naming task whereby the participants named coloured photographs of living (non-human) or non-living items allowed. In the control task, the participants said “OK” in response to a scrambled image.

7) (Chiou et al., 2020): In this study there were three condition self-based decisions, other-based decisions, and a control task. In the self-based decision, the participants read single adjectives describing various personality traits and assessed whether the words suitably describe the characteristics of themselves, similar in the other-based decision task the participants decided whether the depictions suit another individual’s personality (in this case the Queen Elizabeth II). In the control task, the participants made visuospatial decisions on meaningless patterns.

8) (Chiou et al., 2020): In this study there were three conditions—autobiographical memory, theory of mind (ToM), visuospatial control. For the autobiographical memory and ToM task the participants were presented of a picture of event and asked questions that related to the type of event depicted. The questions were either autobiographical “Remember the time you learnt the outcome of Brexit referendum. How did you feel? How did you respond to it?” or ToM “Imagine what the girl who’s holding the Christmas cracker is thinking and feeling. Also imagine how her grandpa would respond”). In the visuospatial control condition the participants judged whether a small triangle was present or absent amongst a scrambled meaningless image.

*Task acquisition parameters:*

With the exception of study 1 and study 4, all studies acquired images using a 3T Philips Achieva scanner using a dual gradient-echo sequence, which has improved signal relative to conventional techniques, especially in areas associated with signal loss (Halai AD et al. 2014). 31 axial slices were collected using a TR = 2.8 seconds, TE = 12 and 35ms, flip angle = 95°, 80 x 79 matrix, with resolution 3 x 3 mm, slice thickness 4mm. The data from study 1 and study 4 MRI data were collected using a Siemens 3 T PRISMA system. T1-weighted images were acquired using a 3D Magnetization Prepared RApid Gradient Echo (MPRAGE) sequence [repetition time (TR) = 2250ms; echo time (TE) = 3.02 ms; inversion time (TI) = 900 ms; 230 Hz per pixel; flip angle = 9°; field of view (FOV) 256 × 256 × 192 mm; GRAPPA acceleration factor 2]. Functional data were acquired using a multi-echo multi-band (MEMB) blood oxygenation level dependent (BOLD)-weighted echo-planar imaging (EPI) pulse sequence. The MEMB sequence had the following parameters: (TR = 1792 ms; TEs = 13, 25.85, 38.7, and 51.55 ms; flip angle = 75°; FOV = 192 mm × 192 mm, MB factor = 2, in-plane acceleration = 3, partial Fourier = 7/8). Each EPI volume consisted of 46 axial slices in descending order (3 x 3mm) covering the whole brain (FOV = 240 x 240 x 138 mm).

*Task data analysis*

The primary question of interest here were: 1) To determine the network engaged by tasks with a sequential nature compared to single-item tasks; 2) To determine whether this effect was driven by tasks that included semantic sequential items or was also true for non-semantic sequential, and 3) To determine to what extent activation of this network was primarily driven by tasks involving a social component (human stimuli) compared to non-social tasks using non-social stimuli. To address this question the tasks were subdivided into subtypes: sequential tasks (with or without semantic content), single item tasks, and social vs. non-social tasks:

Sequential tasks: There were 6 semantic sequential processing tasks: two involving narrative speech production: autobiographical event description (e.g. *Describe the last time you had a cup of tea/coffee)*, semantic event description (e.g. *Describe how you would typically make a cup of coffee)*; two narrative comprehension tasks including human participants; one non-social sentence comprehension task, and one non-social picture sequence comprehension task. There were 4 non-semantic sequential tasks processing tasks: two tasks involving comprehending visual number sequences, one task involving listening to nonverbal classical music, and one task where the participants recited the months of the year.

Single item tasks: The single item stimuli consisted of a mixture of tasks with and without semantic content. To determine the general sequential processing network the semantic and non-semantic sequential tasks were contrasted with tasks involving single-item decision. These of these had a semantic component: object naming, object episodic retrieval, and object semantic retrieval, and four were non-semantic control tasks: three involving some form visuospatial decision, and one where the participant produced a meaningless word to a cue.

Social vs. non-social tasks: Finally, in order to examine the neural network engaged by social compared to non-social semantic processing we compared tasks involving human or self-related decisions vs. non-human semantic decisions. For social tasks we included the autobiographical event description, semantic event description; the two narrative comprehension tasks including human participants, already discussed above. In addition, we included a self-related decision task, an "other person" decision task, and Study 8) an autobiographical imagery task, and a ToM imagery task. These were contrasted with non-social semantic tasks, including non-social sentence comprehension, non-social picture sequence comprehension, object naming, object episodic retrieval, and object semantic retrieval, already referred to above.

*Preprocessing*: The data were analysed using SPM12, implemented in MATLAB. All functional images were corrected for differences in slice acquisition time (slice timing correction), followed by motion correction using a 6-parameter rigid-body transformation and co-registration to each participant's T1-weighted anatomical image using normalized mutual information. Normalisation to standard template space was achieved using DARTEL (Ashburner, 2007), warping all images to MNI space based on the MNI ICBM 152 (non-linear 6th generation) template. Normalized functional images were resampled to 3 mm3 resolution using 4th-degree B-spline interpolation. Spatial smoothing was applied using a 6 mm3 full-width at half maximum (FWHM) isotropic Gaussian kernel.

*General Linear Modelling (GLM)*: The data were analysed using a general linear model using SPM12. Data were high-pass filtered at 128 seconds to remove low-frequency drifts. Each condition for each task was modelled with a separate regressor, and event-related responses were convolved with SPM's canonical hemodynamic response function. At the individual subject level, each condition was modelled against rest. At the first level, statistical analysis was conducted using the general linear model (GLM).

In the second-level analysis, to compare the two task sets, we implemented a second-level flexible factorial model in SPM12. At the second level, a flexible factorial model in SPM12 included factors for Subject (non-independent, equal variance), Study, and Task (17 levels). This structure accounted for repeated measures within subjects and heterogeneity across datasets. Planned contrasts then tested the comparison between sequential tasks > single-item task, semantic sequential tasks vs. non-semantic sequential tasks, and social semantic > non-social semantic tasks. Group analyses were conducted using standard voxel height threshold $p < .001$, cluster corrected using FWE $p < .05$.

**Results**

Sequential tasks > single-item tasks: This contrast revealed a bilateral fronto-temporo-parietal network. The temporo-parietal cortex, bilateral activation extended the full length of the STS from the temporal pole to TPJ. Frontal activation included lateral frontal areas within bilateral IFG and left premotor cortex, as well as medial frontal areas within SMA.

Semantic sequences > non-semantic sequences: This contrast revealed a similar network to the sequential > single-item contrast. Again, a bilateral fronto-temporo-parietal network, including the full length of the STS from the temporal pole to TPJ, but this time frontal activation was limited to the left IFG and premotor cortex.

Social tasks > non-social tasks: Again, this contrast revealed a similar network to the sequential > single-item contrast, and semantic sequences > non-semantic sequences. The contrast revealed activation of a bilateral fronto-temporo-parietal network, including the full length of the STS from the temporal pole to TPJ, as well as bilateral IFG and premotor cortex.

Taken together these results provide evidence that bilateral STS/TPJ responds sequential tasks that include a semantic component, in addition to social semantic tasks. These results could equally be explained by the semantic gestalt as well as the ToM hypothesis.

**ROI results**

In order to examine whether the response in vATL differed to that of the Temporal Pole for the combinatorial semantic tasks compared to the tasks engaging single concept semantics, independent

ROIs were defined for the vATL and Temporal Pole in the left and right hemisphere using Independent Samples t-tests assuming unequal variance.

vATL: The vATL responded equally strongly to the combinatorial semantic and single concept semantic tasks in both the left- ($t(94.50) = .66$, $p = .55$) and right-hemisphere ($t(68.98) = .46$, $p = .65$), with the activation significantly above rest in all conditions (all ts > 2.16, all ps > .01).

Temporal Pole: The temporal pole showed significantly stronger activation for the combinatorial semantic compared to single concepts semantic tasks in both the left - ($t(148.80) = 9.38$, $p < .001$) and right-hemisphere ($t(134.37) = 8.25$, $p < .001$). Activation was significantly above rest for the combinatorial semantic condition (all ts > 10.12 all ps < .001) but did not differ from rest for single concept semantics (all ts < -.86, all ps > .20).

### Section 3. Targeted fMRI studies

**Supplementary 3**

**Methods**

*Participants*

Forty-five participants took part in the fMRI study (average age = 28.24 years, SD = 16.18; N female = 32). Two participants were removed from further analyses due to technical issues with data collection, resulting in a dataset of 43 individuals. All participants were native English speakers with no history of neurological or psychiatric disorders and normal or corrected-to-normal vision. The fMRI experiment was approved by the local ethics committee. Written informed consent was obtained from all participants.

*Task design and procedures*

Experiment 1:

Design and procedure: This experiment used a 2 × 2 design: verbal vs. non-verbal stories, and coherent vs. scrambled structure:

Verbal stories: The coherent condition included 24 three-sentence narratives from Deen et al. (Deen et al., 2015). Each story described a simple event involving a human participant (e.g., "Betty's flight just arrived at the airport. She walks toward the baggage claim area and picks up her suitcase. After taking a cab home, she rushes to her room and gets into bed"). The scrambled verbal stories were built from a separate set of 24 narratives from the same stimulus set, matched on numerous psycholinguistic properties (number of sentences, number of words, mean syllables per word, Flesch reading ease, number of noun phrases, number of modifiers, number of higher level constituents, number of words before the first verb, number of negations, and mean semantic frequency (log Celex frequency). To create scrambled versions, the three sentences were pseudo-randomly swapped across narratives: sentence 1 with sentence 1, sentence 2 with sentence 2, and sentence 3 with sentence 3. This produced a narrative-like sequence nut without a coherent meaning.

Non-verbal stories: The coherent condition included 24 comic strips, each made of nine pictures depicting a clear, unambiguous event with at least one human character. The images were taken from still frames from freely available non-verbal cartoons on YouTube. Stories were chosen from a larger database based on a rating study. Ten independent participants rated each story on a 5-point scale ("How easy is the story to understand?" 1 = very easy, 5 = very hard). They were instructed to use the full scale and to reserve ratings of 1 or 2 for stories that were entirely unambiguous. From these ratings, the 24 easiest stories were selected (M = 1.5, SD = 0.43). The scrambled non-verbal condition was created from a different set of 24 stories. Each nine-picture sequence was divided into three three-picture events, which were then pseudo-randomly shuffled across items using the same procedure as for the verbal stories.

Procedure: Each trial began with a fixation cross for 500 ms. For verbal stories, sentences were shown one at a time across three screens using a word-by-word sliding window paradigm (350 ms per word), with one sentence per screen. The words remained on the screen until the entire sentence was displayed. Average trial duration = 12.55 s (SD = 1.03). For the non-verbal stories: the 9 picture stories were presented with three pictures per screen across three screens. Following fixation, the pictures were revealed one at a time every 1.5 seconds and remained on the screen until the next screen was presented. The total trial duration for the picture trials was 13.5 s. For both tasks, each item was followed by a True/False comprehension question for 2.5 s, answered via button press. Trials were treated as blocks, with ordering optimized using Optseq (http://www.freesurfer.net/optseq). Six 12 s rest blocks with a fixation cross were interspersed. The experiment was divided into 4 runs, each including 6 items from each condition. Each run lasted 403.2 s, and run order was counterbalanced across participants.

Experiment 2

Design: This experiment compared Theory of Mind (ToM) vs. non-ToM narratives. Each condition included 32 items pooled from two previous studies (Deen et al., 2015; Dodell-Feder et al., 2011), shown to elicit a robust ToM > non-ToM effect in the target neural network. A 32-item scrambled control condition was also included but not analysed further.

Procedure: The paradigm was identical to that used for verbal items in Experiment 1. Narratives were presented one sentence at a time across three screens using a word-by-word sliding window paradigm (350 ms per word). Average trial duration = 14.15 s (SD = 1.98). Each trial ended with a 2.5 s True/False comprehension question. As in Experiment 1, trials were treated as blocks, with ordering optimized using Optseq. Nine 12 s rest blocks with a fixation cross were included. The experiment consisted of 4 runs, each with 9 items from each condition. Each run lasted 537.6 s, and run order was counterbalanced across participants.

<u>fMRI acquisition</u>

MRI data were collected using a Siemens 3 T PRISMA system. T1-weighted images were acquired using a 3D Magnetization Prepared RApid Gradient Echo (MPRAGE) sequence [repetition time (TR) = 2250ms; echo time (TE) = 3.02 ms; inversion time (TI) = 900 ms; 230 Hz per pixel; flip angle = 9°; field of view (FOV) 256 × 256 × 192 mm; GRAPPA acceleration factor 2]. Functional data were acquired using a multi-echo multi-band (MEMB) blood oxygenation level dependent (BOLD)-weighted echo-planar imaging (EPI) pulse sequence. The MEMB sequence had the following parameters: (TR = 1792 ms; TEs = 13, 25.85, 38.7, and 51.55 ms; flip angle = 75°; FOV = 192 mm × 192 mm, MB factor = 2, in-plane acceleration = 3, partial Fourier = 7/8). Each EPI volume consisted of 46 axial slices in descending order (3 x 3mm) covering the whole brain (FOV = 240 x 240 x 138 mm). A total 225 volumes were acquired per run for Experiment 1, and 300 volumes were acquired per run for Experiment 2.

Preprocessing

All raw DICOM data were converted to nifti format using dcm2niix. The T1 data were processed using FSL (v5.0.11) (Jenkinson, Beckmann, Behrens, Woolrich, & Smith, 2012; Smith et al., 2004; Woolrich et al., 2009) and submitted to the 'fsl_anat' function. This tool provides a general processing pipeline for anatomical images and involves the following steps (in order): 1) reorient images to standard space ('fslreorient2std'), 2) automatically crop image ('robustfov'), 3) bias-field correction ('fast'), 4) registration to MNI space ('flirt' then 'fnirt'), 5) brain extraction (using fnirt warps) and 6) tissue-type segmentation ('fast'). All images warped to MNI space were visually inspected for accuracy. The functional MEMB data were pre-processed using a combination of tools in FSL, AFNI (v18.3.03) (Cox, 1996) and a python package to perform TE-dependent analysis (DuPre et al., 2020; Kundu et al., 2013; Kundu, Inati, Evans, Luh, & Bandettini, 2012). Despiked (3dDespike), slice time corrected (3dTshift, to the middle slice), and realigned (3dvolreg) images were submitted to the "tedana" toolbox (max iterations = 100, max restarts = 10), which takes the time series from all the collected TEs, decomposes the resulting data into components that can be classified as BOLD or non-BOLD based on their TE-dependence, and then combines the echoes for each component, weighted by the estimated T2* in each voxel, and projects the noise components from the data (Kundu et al., 2012; Poser, Versluis, Hoogduin, & Norris, 2006). The resulting denoised images were then averaged, and the mean image coregistered to T1 (flirt), warped to MNI space (using fnirt warps and flirt transform), and smoothed with 6 mm FHWM Gaussian kernel.

*General Linear Modelling (GLM)*: The data from experiment 1 and experiment 2 were analysed using a general linear model using SPM12. At the individual subject level, each condition was modelled with a separate regressor, and events were convolved with the canonical hemodynamic response function.

Experiment 1:

At the individual level, each task condition was modelled separately against rest. In order to examine the neural networks engaged by each condition, and the overlap between conditions, we first contrasted each condition relative to rest. Next, direct task comparisons, the data were analysed using a 2 x 2 (Language vs. Pictures) factorial ANOVA, where we tested for the main effect of Coherence (Coherent vs. Scrambled) and Modality (Language vs. Picture-cartoons), as well as the interaction. Group analyses were conducted using standard voxel height threshold $p < .001$, cluster corrected using FWE $p < .05$.

*ROI analyses:* In order to investigate our specific hypotheses with regard the STS/TPJ network six 10mm spherical vector ROIs were defined stretching from the anterior temporal pole along the length of STS to the TPJ separately for the left and right hemisphere, based on the *a priori* defined network identified in the fMRI omnibus study. The signal from these ROIs were entered into a 2 x 2 x 6 ANOVA, where we tested for the main effects and interactions between Coherence, Modality, and ROI.

Experiment 2:

At the individual level, each task condition was modelled separately against rest, as well as the direct contrast between ToM > nToM items. The contrasts were then analysed at the group-level using one sample t-tests. Group analyses were conducted using standard voxel height threshold $p < .001$, cluster corrected using FWE $p < .05$.

*Regression Analyses:* We sought to examine the extent to which the ToM effect (ToM > nTOM difference) could be explained by linguistic and semantic factors, rather than ToM per se.

Linguistic variables: To do so we included a number of linguistic variables that have been shown elsewhere to influence processing difficulty. We followed the approach used elsewhere (Shain et al., 2023) whereby a set of linguistic variables was included to capture structural and probabilistic characteristics of each item, and have been shown elsewhere to influence processing difficulty. Importantly, using an identical stimulus set to the current study, within a localiser defined language network the apparent ToM effect is eliminated once controlling for a number of linguistic factors (although it is unclear to what extent this applies elsewhere, beyond the restricted ROIs). The full list of linguistic variables were: 1) Number of Words and 2) Number of Sentences quantified item length in terms of words and sentences, respectively, as language network activity has been shown to vary with linguistic span length (Pallier et al., 2011; Fedorenko et al., 2016) and sentence-level integration processes (Just & Carpenter, 1980; Rayner et al., 2000). 3) Constituent End denoted whether a word concluded a syntactic constituent, allowing examination of potential boundary-related effects on processing (Nelson et al., 2017). 4) Integration Cost, derived from the Dependency Locality Theory (DLT; Gibson, 2000), indexed working memory retrieval demands and followed the variant previously linked to language network activation (Shain et al., 2022). Three complementary measures reflected word predictability: 5) Unigram Surprisal, indexing lexical frequency (Heafield et al., 2013; Graff et al., 2007; Schuster et al., 2016); 6) 5-gram Surprisal, indexing contextual predictability from a 5-gram model (Heafield et al., 2013; Graff et al., 2007; Lopopolo et al., 2017; Shain et al., 2020); and 7) PCFG Surprisal, indexing syntactic predictability from a probabilistic context-free grammar parser (van Schijndel et al., 2013; Nguyen et al., 2012; Marcus et al., 1993; Shain et al., 2020). For an in-depth description of all linguistic variables please refer to Shain et al (2023).

SBERT: In addition to the linguistic variables, a large language model was used to derive a semantic predictability metric for each narrative stimulus. Specifically, Sentence-BERT (SBERT; all-mpnet-base-v2; Reimers & Gurevych, 2019; https://sbert.net, https://huggingface.co/sentence-transformers/all-mpnet-base-v2) was employed to compute semantic similarity between sentences. SBERT is a transformer-based model extending the BERT architecture (Devlin et al., 2019) via a pooling operation and fine-tuning on large-scale natural language inference and semantic textual similarity datasets, generating fixed-length sentence embeddings that capture global semantic content.

For each stimulus (three consecutive sentences), embeddings were obtained for individual sentences and for concatenated context sentences (e.g., S1, S2, S3, and S1+S2). Pairwise cosine similarity was calculated between consecutive or context-augmented sentences (S1–S2, S2–S3, S1–S3, and S1+S2–S3) to quantify semantic relatedness. Higher similarity values indicate that sentences occupy nearby regions in embedding space, suggesting smooth, approximately additive integration into prior context. Lower similarity values indicate a larger semantic shift, suggesting that comprehension depends on supra-additive or cumulative integration, where the meaning of a sentence emerges from the combination of multiple preceding sentences. As these similarity measures were highly correlated, the global average was used as the primary variable of interest (Table S1).

Thus, the SBERT-based cosine similarity metric provides a computational proxy for the ease of semantic integration within narrative contexts, distinguishing between sequences that support linear, additive integration and those that require more complex, cumulative processing. In addition to the linguistic variables, a large language model was used to derive a semantic predictability metric for each item. Specifically, Sentence-BERT (SBERT; model all-mpnet-base-v2; instructions: from https://sbert.net/index.html and https://huggingface.co/sentence-transformers/all-mpnet-base-v2) was employed to compute semantic similarity between sentences. SBERT is a transformer-based model that generates fixed-length vector embeddings representing sentence meaning (Reimers & Gurevych, 2019), extending the BERT architecture (Devlin et al., 2019) via a pooling operation and fine-tuning on natural language inference and semantic textual similarity datasets to produce semantically meaningful sentence-level representations. Pairwise cosine similarity between SBERT embeddings was calculated to quantify semantic relatedness between sentences or between a sentence and its broader context, with higher values indicating greater semantic overlap and lower values indicating reduced coherence. Semantic similarity was computed across multiple sentence pairs (Sentence1–S2, S2–S3, S1–S3, and S1andS2–S3). As these measures were highly correlated, the global average was used as the primary variable of interest (Table S1).

Table S1. SBERT correlations.

| **Correlations** | | | | | |
|---|---|---|---|---|---|
| | S1andS2-to-S3 | S1-to-S2 | S1-to-S3 | S2-to-S3 | Global average |
| S1andS2-to-S3 | 1 | .357** | .830** | .951** | .901** |
| S1-to-S2 | .357** | 1 | .419** | .392** | .630** |

| | | | | | |
|---|---|---|---|---|---|
| S1-to-S3 | .830** | .419** | 1 | .850** | .922** |
| S2-to-S3 | .951** | .392** | .850** | 1 | .939** |
| Global average | .901** | .630** | .922** | .939** | 1 |
| **. Pearson Correlation is significant at the 0.01 level (2-tailed). | | | | | |

Principal Component Analysis (PCA): A principal component analysis with varimax rotation was performed to reduce the large set of correlated variables into a smaller number of underlying factors. This approach addresses multicollinearity among variables and summarises shared variance into orthogonal, interpretable components. Factors with eigenvalues greater than one were retained, yielding four factors that together explained 76.01% of the variance (Table S2 shows the factors and loading scores for each variable). These factors were labelled as: 1) lexico-syntactic complexity (31.81% variance explained), 2) semantic predictability (20.70% variance explained). Factors 3 and 4 were labelled as working-memory-related factors (12.57% and 10.99% variance explained, respectively).

GLM analyses with covariates:

To assess the extent to which task activation (all narratives > rest) was related to the four factors, each factor was entered as a separate covariate in a first-level GLM. At the group level, one-sample t-tests were used to identify activation associated with each factor individually, as well as contrasts between factors. To determine whether the Theory of Mind (ToM) effect (ToM > nToM) could be explained by the four linguistic and semantic factors rather than ToM per se, the GLM was repeated including the ToM contrast alongside the four factors as covariates. Group-level analyses employed a voxel-wise threshold of $p < .001$, with cluster-level family-wise error correction at $p < .05$.

Table S2. PCA factor loadings.

| **Rotated Component Matrix** | | | | |
|---|---|---|---|---|
| | Component | | | |
| | 1 | 2 | 3 | 4 |
| Reading times | -0.292 | 0.840 | -0.148 | -0.073 |
| Narrative length | -0.070 | 0.879 | -0.285 | -0.089 |
| SBERT | -0.090 | -0.770 | -0.020 | -0.133 |
| N Sentences | 0.020 | -0.164 | 0.819 | -0.033 |
| Constituent End | 0.032 | -0.076 | -0.148 | 0.823 |
| Word length | 0.784 | -0.076 | -0.299 | -0.054 |
| Unigram Surprisal | 0.912 | -0.107 | 0.043 | 0.002 |
| 5gram Surprisal | 0.910 | -0.062 | 0.240 | 0.020 |

| | | | | |
|---|---|---|---|---|
| PCFG surprisal | 0.918 | -0.028 | 0.060 | 0.092 |
| Integration Cost | 0.470 | 0.219 | 0.519 | 0.275 |
| Storage Cost | 0.045 | 0.266 | -0.786 | 0.242 |
| Embedding depth | 0.013 | 0.070 | 0.014 | 0.862 |
| Extraction Method: Principal Component Analysis. Rotation Method: Varimax with Kaiser Normalization. | | | | |

fMRI study

Experiment 1

*Whole-brain analyses: task similarities*

Language > rest: Relative to rest, coherent language engaged a fully bilateral fronto-temporo-parietal network, as well as visual processing areas. Specifically, in the temporal lobe, activation spanned the length of the STS/TPJ, including the entirety of the temporal pole, as well as the posterior-anterior fusiform gyrus. Frontal activation included lateral frontal cortex (IFG, middle frontal, premotor cortex) as well as medial frontal areas (SMA, vMPFC, dMPFC, orbito-frontal cortex), and parietal activation included SMG, anterior AG, as well IPS. This activation pattern overlaps with the known language network, as well as the MD system. Interestingly, the scrambled language condition engaged an entirely overlapping network, the only visual difference being a slightly reduced volume of activation in the temporal lobe but mildly more extensive in fronto-parietal areas. This suggests that any significant difference between conditions will be quantitative rather than reflecting a qualitatively different pattern of recruitment. All peak MNI coordinates are reported in Table S3.

Pictures > rest: Relative to rest, the coherent picture condition engaged a strikingly similar neural network as the language task, engaging a largely overlapping bilateral fronto-temporo-parietal network. This included the length of the STS/TPJ, including the entirety of the temporal pole, as well as the posterior-anterior fusiform gyrus. Frontal activation included lateral frontal cortex (IFG, middle frontal, premotor cortex) as well as medial frontal areas (SMA, vMPFC, dMPFC, orbito-frontal cortex), and parietal activation included SMG, anterior AG, as well IPS. In addition to this, as expected the visual task also engaged an extensive occipito-parietal activation, spanning the entirety of the occipital lobe, and extending dorsally into superior parietal cortex, as well as the posterior AG and IPS. As in the language task, the scrambled picture condition engaged a fully overlapping network to the coherent picture condition, with only moderate visual differences in cluster size. Together these results provide evidence that both the language and picture task engage a common multi-modal semantic network. All peak MNI coordinates are reported in Table S3.

*Whole-brain analyses: task differences:*

Despite the high degree of overlap across networks, the results from the factorial ANOVA indicate a significant main effect of modality, coherence, as well as a modality x coherence interaction. All peak MNI coordinates are reported in Table S3.

Modality: In terms of modality, relative to the picture task, the language conditions showed stronger engagement of the bilateral fronto-temporal system that has been commonly associated with language processing elsewhere (refs), whereas as expected, the picture task more strongly engaged posterior occipito-parietal areas, as well as the ventral temporal lobe in areas associated with visual recognition.

Coherence: As predicted, the Coherent > Scrambled contrast showed bilateral recruitment of the length of the STS/TPJ network with activation including the full extent of the temporal pole, and a large TPJ/pMTG cluster. In addition to this, the bilateral IFG and medial dorsal prefrontal cortex were also recruited more strongly for the coherent compared to the scrambled conditions. In contrast, the Scrambled > Coherent conditions revealed a bilateral fronto-parietal network that is commonly associated with executive processing and overlapped with the MD system. This included bilateral middle frontal gyrus, premotor cortex, SMA, the IPS, and posterior ITG. In addition to this, a bilateral cluster of activation was revealed in mid-STG for Scrambled > Coherent contrast which was absent when examining the activation compared to rest, suggesting this reflects a pattern of differential deactivation, thereby making it hard to interpret.

Interaction: A significant interaction was revealed in the left IFG, left STS/TPJ network, as well as large areas of bilateral parietal cortex, including TPJ, SMG, and the SPL. Direct task comparisons showed that this pattern reflected a larger coherence effect in the left STS Language compared to Picture task, but a stronger coherence effect for the Picture task in bilateral parietal cortex compared to the Language task.

*ROI analyses*

In order to investigate our specific hypotheses with regard the STS/TPJ network bilateral vector ROIs were defined stretching from the anterior temporal pole along the length of STS to the TPJ, as well as the dorsal-ventral axis of the temporal pole. Beginning with the STS/TPJ vector ROIs, the mean signal for each subject was entered into a 2 x 2 x 6 within-subject ANOVA, where we tested for the main effects and interactions between Coherence, Modality, and ROI anterior-posterior location. To simplify the interpretation of results we initially tested for Coherence, Modality, and ROI effects within the left- and right-hemisphere separately, before conducting cross-hemisphere comparisons.

ATL-STS-TPJ vector ROIs:

Left hemisphere: Starting with the left hemisphere, this revealed a significant effect of Coherence ($F(42) = 68.79$, $p < .001$), Modality ($F(42) = 219.09$, $p < .001$), and ROI ($F(42) = 18.66$, p

< .001). In addition, there was a significant Coherence x ROI interaction (F(42) = 10.49, p < .001), as well as a Modality x ROI interaction (F(42) = 38.69, p < .001), and a significant Coherence x Modality x ROI interaction (F(42) = 7.55, p < .001). Interestingly, no evidence was found for a significant Coherence x Modality interaction (F(42) = 0.07, p = .49). Pair-wise comparisons applying a Bonferroni correction show that the size of the coherence effect was significantly greater in the left TPJ compared to all other left STS ROIs (all ts > 3.01, all ps < .001), in addition STS2 showed a significantly smaller effect than all other regions (all ts > 3.01, all ps < .001) (see stats LSTG1_diff – LSTG2_diff etc). In terms of Modality, the size of the modality effect was significantly smaller in the LTPJ compared to all other LSTS regions (all ts > 3,60, all ps < .001) except STS1 (t(42) – 2.13, p = .04). In contrast, LSTS4 showed a significantly larger modality difference compared to all other regions (stronger language activation compared to picture activation) (all ts > 4.91, all ps < .001), and LSTS3 showing a significantly greater modality effect to LSTS1, LSTS2, and LTPJ (all ts > 6.42, all ps < .001) but only a marginal difference with LSTS5 (t(42) = 2.70, p =.01). No other comparisons were significantly difference. Finally, in order to unpack the Coherence x Modality x ROI interaction, pairwise comparisons showed that the size of the coherence effect was found to be significantly greater for the picture task compared to the language task within the RSTS5 compared to all other regions (all ts > 4.31, all ps < .001; all ts > 5.58, all ps < .001), with the exception of LTPJ which did not differ significantly (t(42) = 1.95, p =.06), as well as the LPTJ showing a larger picture coherence effect compared language coherence effect compared to the picture coherence effect compared language coherence effect LSTS4 (t(42) = 3.42, p <.001). No other comparisons were significantly difference.

Right hemisphere: For the right hemisphere STS-TPJ ROIs, a 2 x 2 x 6 within-subject ANOVA revealed a significant effect of Coherence (F(42) = 93.75, p < .001), and ROI (F(42) = 13.05, p < .001). But no significant main effect of Modality (F(42) = 0.31, p = .58). In addition there was a significant Coherence x ROI interaction (F(42) = 9.21, p < .001), a Modality x ROI interaction (F(42) = 71.72, p < .001), a marginal Coherence x Modality interaction (F(42) = 4.19, p = .05), and a significant Coherence x Modality x ROI interaction (F(42) = 26.75, p < .001). IN order to interpret the Coherence x ROI interaction, pair-wise comparisons showed that coherence effect in RPTJ was significantly smaller compared to RSTS3, RSTS4, and RSTS5 (all ts > 2.88, all ps < .006) and marginally less than RSTS1 (t(42) = 2.15, p = .05), this is in contrast to the left hemisphere where LTPJ showed the largest effect across ROIs. The coherence effect was also significantly reduced in the RSTS2 compared to all regions (all ts > 3.57, all ps < .001) except the RTPJ where there was no significant difference (t(42) = 1.25, p = .22). In order to understand the modality x ROI interaction, pairwise comparisons revealed that all regions showed a significantly greater modality effect compared to RSTS1 (all ts > -3.27, all ps < .002) and STS2 (all ts > -3.96, all ps < .001), whilst STS1 and STS2 did not differ significantly (t(42) = .56, p = .58). In contrast, the largest modality effect was found in STS3 (all ts > 5.00, all ps < .001) and STS4 (all ts > -3.27, all ps < .002), which did not differ significantly from one-another (t(42) = .06, p = .95). Additionally, whilst the modality effect in RSTS5 was smaller than STS3 and STS4, RSTS5 did show a significantly larger modality effect compared to RSTS1, RSTS2, and RTPJ (all ts > 5.23, all ps < .001). Pairwise comparisons were also conducted to examine the marginal Coherence x Modality interaction, however there was only a marginally significant difference in the average size of the coherence effect across regions, with a marginally larger effect for the picture task compared to language task (t(42) = .2.05, p = .05). Finally, in order to unpack the Coherence x Modality x ROI interaction, pairwise comparisons showed that the size of the coherence effect was found to be significantly greater for the picture task compared to the language task within the RTPJ and RSTS5 compared to all other regions (all ts > 4.31, all ps < .001; all ts > 5.58, all ps < .001), whilst RTPJ and RSTS5 did not differ significantly from one another (t(42) = .57, p = .58). No other comparisons were significantly difference.

Hemispheric differences: To examine the influence the extent to which activation differed across hemispheres, we averaged the condition activity across the STS/TPJ ROIs and entered this into a 2 (hemisphere) x 2 (modality) x 2 (coherence) within-subject ANOVA. This revealed a significant effect of hemisphere (F(42) = 5.37, p = .03) reflecting stronger overall activation in the left compared to right hemisphere across tasks, a significant effect of modality (F(42) = 79.26, p < .001) reflecting overall stronger activation for the language task compared to the picture task across both hemispheres, a significant hemisphere x modality interaction (F(42) = 103.37, p < .001), as well as a hemisphere x coherence interaction (F(42) = 6.49, p < .001). Pairwise comparisons showed that the hemisphere x modality reflected greater language activation in the left-hemisphere compared to right-hemisphere (t42) = 6.31, p < .001), and greater picture activation in the right-hemisphere compared to the left-hemisphere (t42) = 5.00, p < .001). The hemisphere x coherence interaction can be explained by marginally greater activation for the scrambled conditions in the left- vs. right-hemisphere (t42) = 2.62, p = .01), but no significant cross hemisphere difference in the coherent condition (t42) = 1.97, p = .06).

Temporal Pole ROIs:

The six temporal pole ROIs were entered into a 2 (coherence) x 2 (modality) x 3 (ROI) x 2 (hemisphere) ANOVA. This revealed a significant main effect of hemisphere (F(42) = 20.83, p <.001), a significant main effect of ROI (F(42) = 117.58, p <.001), a significant main effect of modality ((F(42) = 83.03, p <.001), and a significant main effect of coherence (F(42) = 92.20, p <.001). In addition there was a significant hemisphere x modality interaction (F(42) = 25.60, p <.001), a significant ROI x modality interaction (F(42) = 15.70, p <.001), a significant hemisphere x ROI x modality interaction ((F(42) = 9.70, p <.001), a significant hemisphere x coherence interaction ((F(42) = 4.96, p = .03), a significant ROI x coherence interaction ((F(42) = 14.79, p <.001) reflecting a stronger coherence effect in the superior temporal pole compared to mid- and ventral-region, and a significant modality x coherence interaction ((F(42) = 7.90, p =.007).

Ventral ATL vs. Lateral Temporal ROIs:

In order to examine whether the response in vATL differed to that of the lateral temporal polar region for the coherent vs, scrambled ROIs analyses were conducted within the same independent bilateral vATL and Temporal Pole ROIs as above.

In the left hemisphere, 2 x 2 ANOVA comparing the size of the coherence effect for both tasks in vATL and Temporal Pole ROI revealed a significant variation in the size of coherence effect across tasks ($F(42) = 12.36$, $p < .001$), a significant effect of ROI ($F(42) = 65.31$, $p < .001$), and as a significant coherence effect x ROI interaction ($F(42) = 14.36$, $p < .001$). Similarly, in the right hemisphere, a 2 x 2 ANOVA showed size of coherence effect across tasks ($F(42) = 4.10$, $p = .05$), a significant effect of ROI ($F(42) = 50.99$, $p < .001$), and as a significant coherence effect x ROI interaction ($F(42) = 10.32$, $p < .005$). Follow-up t-test showed that the temporal pole showed significantly stronger activation for the combinatorial semantic compared to single concepts semantic tasks for both the language and picture task in the both the left- and right hemisphere (all ts > = 6.12 $p < .001$). In contrast, the vATL responded equally strongly to the coherent and scrambled conditions in neither the language or picture task, in either hemisphere (all ts < .84, all ps = .1) with the activation significantly above rest in all conditions (all ts > 2.16, all ps > .01) .

Experiment 2

*ToM and nTom task activation:* Compared to rest, the ToM narratives and nToM narratives activated an entirely overlapping network of regions, nearly identical to that engaged by the coherent narratives in study 1. That is a fully bilateral fronto-temporo-parietal network, as well as visual processing areas. Specifically, in the temporal lobe, activation spanned the length of the STS/TPJ, including the entirety of the temporal pole, as well as the posterior-anterior fusiform gyrus. Frontal activation included lateral frontal cortex (IFG, middle frontal, premotor cortex) as well as medial frontal areas (SMA, vMPFC, dMPFC, orbito-frontal cortex), and parietal activation included SMG, anterior AG, as well IPS. Despite the overlap across conditions, the Tom > nToM contrast revealed that the ToM task elicited significantly stronger activation throughout the entirety of the fronto-temporo-parietal network (with the exception of primary motor cortex, dorsal parietal areas, and visual cortex which did not differ between conditions). The bilateral precuneus was the only area to be revealed by the ToM > nToM contrast that was not present when contrasting task > rest. All peak MNI coordinates are reported in Table S3.

*Regression analyses:*

Factor 1: Factor 1 was associated with activation spanned the length of the STS/TPJ, including the entirety of the temporal pole, as well as the posterior-anterior fusiform gyrus. Frontal activation included lateral frontal cortex (IFG, middle frontal, premotor cortex) as well as medial frontal areas (SMA, vMPFC, dMPFC, orbito-frontal cortex). This pattern directly mirrored that of the ToM > nToM contrast, with the exception of the precuneus which was not significantly modulated by Factor 1.

Factor 2: Factor 2 was specifically associated with activation along the length of the STS/TPJ network bilaterally, as well as the bilateral precuneus.

Factor 1 > Factor 2: Direct contrasts of Factor 2 > Factor 1 showed that activation of the bilateral anterior temporal lobe, TPJ, and precuneus was stronger for Factor 2, whereas the reverse contrast revealed stronger activation for Factor 1 in left lateral prefrontal cortex, as well as left posterior fusiform gyrus.

Factor 3 and Factor 4: Factors 3 and 4 were not significantly associated with any neural activation.

All Factors > ToM contrast: In order to determine to what extent task activation could be explained by the PCA factors over and above the ToM > contrast we directly compared the activation for all Factors combined with the ToM contrast (ToM > nToM), and vice versa. All Factors combined were associated with significantly greater activation than the ToM contrast within the bilateral entire STS/TPJ network, as well as bilateral middle frontal gyrus, and midline areas within anterior medial prefrontal cortex and precuneus. Nevertheless, not all ToM contrast activation could be explained by the PCA Factors. Indeed, the ToM contrast revealed significantly greater activation compared to the combined Factors within bilateral insular and orbitofrontal cortex, bilateral anterior cingulate, and right frontal pole.

All peak MNI coordinates are reported in Table S3.

Table S3: The peak MNI coordinates from fMRI Experiment 1 and Experiment 2

| | p(FWE-corr) | p(FDR-corr) | equivk | T | equivZ | x,y,z {mm} | x,y,z {mm} | x,y,z {mm} |
|---|---|---|---|---|---|---|---|---|
| Experiment 1 | | | | | | | | |
| Language Coherent > rest | 0.00 | 0.00 | 18222 | 21.36 | 65535.00 | 28 | -95 | 2 |
| | | | | 15.04 | 65535.00 | 34 | -89 | -3 |
| | | | | 14.93 | 65535.00 | -7 | -31 | -5 |
| | | | | 14.93 | 65535.00 | 10 | -77 | -39 |
| | | | | 14.09 | 65535.00 | 24 | -29 | -1 |
| | | | | 14.00 | 65535.00 | 38 | -85 | -13 |
| | | | | 13.07 | 65535.00 | 8 | -75 | -21 |
| | | | | 12.97 | 65535.00 | 28 | -69 | -53 |
| | 0.00 | 0.00 | 20084 | 16.51 | 65535.00 | -29 | -91 | -11 |
| | | | | 16.37 | 65535.00 | -39 | -89 | -11 |
| | | | | 15.59 | 65535.00 | -39 | -47 | -19 |
| | | | | 15.18 | 65535.00 | -21 | -95 | -7 |
| | | | | 14.32 | 65535.00 | -43 | -81 | -13 |
| | | | | 14.00 | 65535.00 | -45 | -69 | -15 |
| | | | | 13.85 | 65535.00 | -49 | 2 | 52 |
| | | | | 13.84 | 65535.00 | -43 | -83 | -1 |
| | 0.02 | 0.00 | 161 | 13.95 | 65535.00 | -19 | -39 | -43 |
| | 0.02 | 0.00 | 164 | 12.99 | 65535.00 | 20 | -39 | -45 |
| | 0.00 | 0.00 | 4809 | 10.89 | 7.46 | -5 | -1 | 68 |
| | | | | 7.77 | 6.08 | 46 | -3 | 32 |
| | | | | 6.86 | 5.59 | 10 | -29 | 60 |
| | | | | 6.60 | 5.43 | 66 | -3 | 26 |
| | | | | 6.36 | 5.29 | -9 | -25 | 62 |
| | | | | 6.25 | 5.23 | 46 | 28 | 18 |
| | | | | 6.04 | 5.09 | 56 | -7 | 36 |
| | | | | 5.99 | 5.06 | 30 | -1 | 52 |
| | 0.04 | 0.01 | 143 | 6.60 | 5.43 | -33 | -69 | -53 |
| | 0.08 | 0.01 | 114 | 6.55 | 5.41 | -1 | -35 | -43 |
| | | | | 3.65 | 3.38 | -3 | -25 | -43 |
| | 0.09 | 0.01 | 113 | 5.70 | 4.88 | -29 | -13 | -17 |
| | | | | 5.58 | 4.80 | -19 | -11 | -23 |
| | 0.12 | 0.01 | 104 | 4.62 | 4.13 | 12 | -13 | 4 |
| | | | | 4.34 | 3.92 | 18 | -21 | 10 |
| | 0.09 | 0.01 | 113 | 4.24 | 3.85 | 2 | 48 | -21 |
| Language Scrambled > rest | 0.00 | 0.00 | 39416 | 21.92 | 65535.00 | 26 | -95 | -1 |
| | | | | 17.02 | 65535.00 | -29 | -91 | -11 |
| | | | | 16.69 | 65535.00 | -39 | -89 | -11 |
| | | | | 15.77 | 65535.00 | -19 | -95 | -7 |
| | | | | 15.64 | 65535.00 | 34 | -89 | -3 |

| | | | | | | | | |
|---|---|---|---|---|---|---|---|---|
| | | | | 15.03 | 65535.00 | -47 | -61 | -17 |
| | | | | 14.93 | 65535.00 | 10 | -81 | -43 |
| | | | | 14.80 | 65535.00 | -49 | 2 | 52 |
| | 0.00 | 0.00 | 1764 | 14.40 | 65535.00 | -7 | -29 | -5 |
| | | | | 14.15 | 65535.00 | 26 | -29 | -1 |
| | | | | 12.41 | 65535.00 | 8 | -31 | -3 |
| | | | | 11.30 | 7.62 | -23 | -29 | -3 |
| | | | | 8.27 | 6.34 | -11 | -15 | 6 |
| | | | | 7.68 | 6.03 | 6 | -23 | -9 |
| | | | | 7.32 | 5.84 | -3 | -21 | -11 |
| | | | | 7.08 | 5.71 | -13 | -19 | -13 |
| | 0.04 | 0.01 | 135 | 11.07 | 7.53 | -21 | -39 | -43 |
| | 0.00 | 0.00 | 935 | 10.90 | 7.47 | -5 | -1 | 68 |
| | | | | 7.85 | 6.13 | -9 | 12 | 54 |
| | | | | 6.21 | 5.20 | 8 | 12 | 52 |
| | 0.06 | 0.01 | 124 | 10.44 | 7.29 | 20 | -39 | -45 |
| | 0.02 | 0.00 | 158 | 8.88 | 6.63 | -33 | -69 | -53 |
| | 0.01 | 0.00 | 198 | 8.85 | 6.61 | 22 | -13 | -21 |
| Picture Coherent > rest | 0.00 | 0.00 | 53421 | 28.93 | 65535.00 | 24 | -31 | 2 |
| | | | | 24.68 | 65535.00 | 34 | -45 | -15 |
| | | | | 24.51 | 65535.00 | -21 | -31 | -3 |
| | | | | 23.40 | 65535.00 | 28 | -95 | -1 |
| | | | | 22.94 | 65535.00 | 46 | -75 | -7 |
| | | | | 22.89 | 65535.00 | 18 | -91 | -11 |
| | | | | 22.16 | 65535.00 | -19 | -101 | -3 |
| | | | | 22.11 | 65535.00 | 18 | -95 | -1 |
| | 0.00 | 0.00 | 3604 | 10.07 | 7.14 | 48 | 26 | 22 |
| | | | | 10.04 | 7.13 | 46 | 18 | 24 |
| | | | | 9.64 | 6.96 | 32 | -1 | 56 |
| | | | | 9.03 | 6.70 | 42 | 10 | 26 |
| | | | | 8.10 | 6.25 | 56 | 38 | 6 |
| | | | | 7.50 | 5.94 | 48 | -3 | 32 |
| | | | | 7.46 | 5.92 | 46 | 4 | 44 |
| | | | | 6.57 | 5.42 | 44 | 2 | 52 |
| | 0.00 | 0.00 | 2932 | 9.99 | 7.11 | -49 | 14 | 28 |
| | | | | 8.99 | 6.68 | -47 | -1 | 50 |
| | | | | 8.99 | 6.68 | -31 | -3 | 50 |
| | | | | 8.48 | 6.44 | -43 | 32 | -13 |
| | | | | 8.12 | 6.26 | -43 | 2 | 32 |
| | | | | 8.00 | 6.20 | -41 | -5 | 44 |
| | | | | 6.19 | 5.19 | -57 | -1 | 50 |
| | | | | 6.01 | 5.08 | -57 | 30 | 22 |
| | 0.00 | 0.00 | 867 | 9.90 | 7.07 | 2 | 52 | -19 |
| | | | | 7.67 | 6.03 | 2 | 38 | -23 |

| | | | | | | | | |
|---|---|---|---|---|---|---|---|---|
| | 0.00 | 0.00 | 301 | 6.70 | 5.50 | -7 | 6 | 62 |
| | | | | 6.06 | 5.11 | -11 | 12 | 54 |
| | 0.00 | 0.00 | 312 | 6.26 | 5.23 | -33 | -25 | 56 |
| | | | | 3.89 | 3.57 | -35 | -31 | 48 |
| | 0.04 | 0.01 | 140 | 6.24 | 5.22 | -3 | 20 | 10 |
| | | | | 4.54 | 4.07 | -13 | 28 | 8 |
| Picture Scrambled > rest | 0.00 | 0.00 | 51472 | 28.67 | 65535.00 | 22 | -31 | 2 |
| | | | | 24.33 | 65535.00 | 28 | -95 | -1 |
| | | | | 24.27 | 65535.00 | 34 | -45 | -15 |
| | | | | 23.36 | 65535.00 | 18 | -91 | -9 |
| | | | | 22.72 | 65535.00 | 18 | -95 | -1 |
| | | | | 21.93 | 65535.00 | 36 | -33 | -21 |
| | | | | 21.54 | 65535.00 | -19 | -101 | -3 |
| | | | | 21.12 | 65535.00 | -29 | -91 | -11 |
| | 0.00 | 0.00 | 3657 | 9.66 | 6.97 | 46 | 26 | 24 |
| | | | | 9.58 | 6.94 | 44 | 18 | 24 |
| | | | | 9.21 | 6.78 | 42 | 10 | 28 |
| | | | | 8.84 | 6.61 | 32 | 4 | 56 |
| | | | | 7.87 | 6.14 | 44 | 4 | 42 |
| | | | | 7.01 | 5.67 | 48 | -3 | 32 |
| | | | | 6.74 | 5.52 | 26 | 2 | 48 |
| | | | | 5.40 | 4.68 | 58 | 34 | 4 |
| | 0.00 | 0.00 | 3150 | 9.11 | 6.73 | -47 | 14 | 28 |
| | | | | 8.66 | 6.53 | -47 | -1 | 52 |
| | | | | 8.58 | 6.48 | -31 | -3 | 50 |
| | | | | 8.32 | 6.36 | -43 | 2 | 32 |
| | | | | 7.52 | 5.95 | -41 | -5 | 44 |
| | | | | 7.36 | 5.87 | -7 | 4 | 62 |
| | | | | 6.89 | 5.61 | -11 | 10 | 56 |
| | | | | 6.38 | 5.30 | -57 | -1 | 50 |
| | 0.09 | 0.01 | 109 | 8.58 | 6.49 | 8 | 2 | 30 |
| | | | | 5.86 | 4.98 | -5 | 2 | 28 |
| | 0.02 | 0.00 | 166 | 8.39 | 6.39 | 32 | -71 | -53 |
| | 0.00 | 0.00 | 826 | 7.69 | 6.04 | -1 | 60 | -15 |
| | | | | 7.54 | 5.96 | 4 | 48 | -19 |
| | | | | 7.00 | 5.67 | -7 | 52 | -17 |
| | | | | 6.05 | 5.10 | 4 | 38 | -23 |
| | | | | 3.65 | 3.38 | 2 | 50 | -31 |
| | 0.04 | 0.01 | 140 | 7.48 | 5.93 | -43 | 32 | -15 |
| | 0.11 | 0.01 | 103 | 6.70 | 5.50 | -5 | 22 | 8 |
| | | | | 4.21 | 3.82 | -13 | 28 | 6 |
| | 0.00 | 0.00 | 259 | 6.20 | 5.19 | -31 | -23 | 56 |
| Main effect of Coherence: All Normal > scrambled | 0.00 | 0.00 | 8373 | 11.30 | 65535.00 | -65 | -33 | 32 |
| | | | | 9.24 | 65535.00 | -51 | -67 | 2 |

| | | | | | | | | |
|---|---|---|---|---|---|---|---|---|
| | | | | 8.81 | 7.75 | -13 | -57 | 66 |
| | | | | 8.78 | 7.73 | 12 | -55 | 68 |
| | | | | 7.09 | 6.49 | -13 | -27 | 38 |
| | | | | 6.88 | 6.33 | -3 | -5 | 40 |
| | | | | 6.71 | 6.20 | -65 | -47 | 34 |
| | | | | 6.70 | 6.19 | 28 | -51 | 74 |
| | 0.00 | 0.00 | 5262 | 9.53 | 65535.00 | 60 | -27 | 38 |
| | | | | 9.18 | 65535.00 | 68 | -29 | 34 |
| | | | | 7.90 | 7.11 | 56 | -59 | 2 |
| | | | | 6.71 | 6.19 | 48 | -63 | -7 |
| | | | | 6.32 | 5.88 | 68 | -37 | 26 |
| | | | | 6.01 | 5.63 | 60 | -47 | 12 |
| | | | | 5.86 | 5.50 | 48 | -33 | -3 |
| | | | | 5.61 | 5.29 | 58 | -23 | -7 |
| | 0.00 | 0.00 | 4888 | 8.97 | 65535.00 | 8 | 58 | 38 |
| | | | | 8.45 | 7.50 | -9 | 34 | 58 |
| | | | | 8.29 | 7.39 | 6 | 56 | 30 |
| | | | | 8.06 | 7.22 | 8 | 62 | 16 |
| | | | | 7.52 | 6.82 | 10 | 36 | 58 |
| | | | | 7.33 | 6.68 | -11 | 54 | 38 |
| | | | | 7.30 | 6.65 | -11 | 54 | 16 |
| | | | | 7.16 | 6.54 | -11 | 42 | 50 |
| | 0.00 | 0.00 | 891 | 7.65 | 6.92 | -31 | -81 | -33 |
| | | | | 4.76 | 4.56 | -17 | -77 | -47 |
| | 0.00 | 0.00 | 1711 | 7.63 | 6.91 | 56 | 12 | -29 |
| | | | | 6.98 | 6.41 | 50 | 20 | -31 |
| | | | | 6.93 | 6.37 | 52 | 14 | -39 |
| | | | | 6.75 | 6.22 | 48 | 6 | -41 |
| | | | | 6.71 | 6.20 | 38 | 24 | -29 |
| | | | | 5.92 | 5.55 | 40 | 26 | -37 |
| | | | | 5.68 | 5.35 | 34 | 26 | -17 |
| | | | | 5.56 | 5.25 | 28 | 18 | -17 |
| | 0.00 | 0.00 | 1080 | 6.98 | 6.41 | -51 | 4 | -37 |
| | | | | 5.68 | 5.35 | -57 | 10 | -31 |
| | | | | 5.28 | 5.01 | -51 | 14 | -37 |
| | | | | 5.05 | 4.81 | -43 | 22 | -35 |
| | | | | 4.97 | 4.74 | -39 | 24 | -17 |
| | | | | 4.53 | 4.35 | -31 | 18 | -17 |
| | | | | 3.99 | 3.86 | -57 | -11 | -27 |
| | | | | 3.45 | 3.37 | -41 | -1 | -41 |
| | 0.00 | 0.00 | 1109 | 6.98 | 6.41 | 24 | -3 | -19 |
| | | | | 5.81 | 5.46 | 36 | -11 | -1 |
| | | | | 5.51 | 5.20 | -3 | -21 | 18 |
| | | | | 5.46 | 5.16 | 14 | -5 | 16 |

| | | | | | | | | |
|---|---|---|---|---|---|---|---|---|
| | | | | 4.78 | 4.57 | 26 | -13 | -9 |
| | | | | 4.50 | 4.32 | 18 | -17 | 14 |
| | | | | 4.35 | 4.19 | 24 | -3 | 8 |
| | | | | 4.20 | 4.05 | 6 | 6 | 2 |
| | 0.00 | 0.00 | 377 | 6.80 | 6.26 | -53 | 4 | 24 |
| | 0.00 | 0.00 | 296 | 6.40 | 5.95 | -21 | -1 | 68 |
| | | | | 4.51 | 4.33 | -23 | -11 | 64 |
| | 0.00 | 0.00 | 596 | 6.30 | 5.86 | -21 | -5 | -13 |
| | | | | 6.10 | 5.70 | -31 | -11 | -5 |
| | | | | 4.79 | 4.59 | -39 | 2 | 2 |
| | | | | 4.00 | 3.87 | -21 | 4 | -7 |
| | | | | 3.68 | 3.58 | -27 | -1 | -23 |
| | 0.00 | 0.00 | 699 | 6.15 | 5.74 | 28 | -79 | -33 |
| | | | | 5.85 | 5.50 | 14 | -75 | -49 |
| | 0.01 | 0.00 | 196 | 5.59 | 5.27 | -49 | -47 | -19 |
| | 0.00 | 0.00 | 252 | 4.99 | 4.76 | 56 | 38 | 4 |
| | | | | 4.83 | 4.62 | 50 | 32 | -7 |
| | | | | 3.57 | 3.48 | 54 | 30 | 8 |
| | 0.01 | 0.00 | 225 | 4.90 | 4.68 | -19 | -19 | 16 |
| | | | | 4.27 | 4.12 | -13 | -5 | 16 |
| | | | | 4.12 | 3.99 | -15 | -27 | 6 |
| | | | | 3.45 | 3.37 | -21 | -23 | 24 |
| | 0.12 | 0.02 | 112 | 4.77 | 4.56 | -35 | 30 | 40 |
| | | | | 3.99 | 3.87 | -39 | 34 | 46 |
| | 0.01 | 0.00 | 208 | 4.74 | 4.54 | 56 | 8 | 24 |
| | | | | 4.17 | 4.03 | 54 | 10 | 14 |
| | | | | 3.92 | 3.80 | 60 | 18 | 18 |
| | 0.02 | 0.00 | 184 | 4.26 | 4.11 | 30 | -43 | -43 |
| Main Effect of Coherence: All scrambled> Normal | 0.00 | 0.00 | 7766 | 11.67 | 65535.00 | -7 | -27 | 30 |
| | | | | 11.65 | 65535.00 | 6 | -29 | 30 |
| | | | | 11.60 | 65535.00 | -7 | -35 | 28 |
| | | | | 10.18 | 65535.00 | -11 | -67 | 34 |
| | | | | 9.84 | 65535.00 | 8 | -37 | 26 |
| | | | | 9.57 | 65535.00 | -13 | -69 | 22 |
| | | | | 9.41 | 65535.00 | 8 | -75 | 44 |
| | | | | 8.44 | 7.50 | -11 | -49 | 10 |
| | 0.00 | 0.00 | 1230 | 10.04 | 65535.00 | -39 | -55 | 42 |
| | | | | 7.39 | 6.72 | -33 | -71 | 44 |
| | 0.00 | 0.00 | 1317 | 9.50 | 65535.00 | 36 | -63 | 46 |
| | 0.01 | 0.00 | 226 | 7.41 | 6.74 | -39 | -73 | -53 |
| | 0.00 | 0.00 | 239 | 7.31 | 6.66 | 34 | -75 | -53 |
| | 0.00 | 0.00 | 1298 | 7.24 | 6.61 | 48 | 34 | 28 |
| | | | | 7.01 | 6.43 | 36 | 54 | 6 |
| | | | | 4.49 | 4.32 | 30 | 66 | 4 |

| | | | | | | | | |
|---|---|---|---|---|---|---|---|---|
| | | | | 4.09 | 3.95 | 24 | 56 | 2 |
| | 0.00 | 0.00 | 1114 | 7.04 | 6.45 | -43 | 54 | 10 |
| | | | | 6.59 | 6.10 | -35 | 50 | -1 |
| | | | | 5.70 | 5.37 | -21 | 56 | 2 |
| | 0.00 | 0.00 | 897 | 6.98 | 6.41 | -7 | 14 | 50 |
| | | | | 6.64 | 6.13 | 2 | 16 | 48 |
| | | | | 4.76 | 4.56 | 8 | 28 | 38 |
| | 0.06 | 0.01 | 135 | 6.49 | 6.02 | -21 | -17 | -19 |
| | 0.00 | 0.00 | 1132 | 6.33 | 5.88 | -45 | 24 | 26 |
| | | | | 6.24 | 5.82 | -43 | 16 | 34 |
| | | | | 5.85 | 5.49 | -33 | 18 | 2 |
| | | | | 5.76 | 5.42 | -41 | 24 | 18 |
| | | | | 3.97 | 3.84 | -39 | 2 | 32 |
| | | | | 3.55 | 3.45 | -33 | 20 | 14 |
| | 0.00 | 0.00 | 307 | 6.25 | 5.82 | 30 | 38 | -9 |
| | 0.00 | 0.00 | 313 | 6.24 | 5.81 | 64 | -43 | -15 |
| | 0.01 | 0.00 | 222 | 6.22 | 5.80 | 34 | 16 | 56 |
| | 0.00 | 0.00 | 280 | 5.79 | 5.44 | 36 | 20 | -1 |
| | 0.00 | 0.00 | 361 | 5.55 | 5.24 | -39 | -29 | 8 |
| | | | | 4.88 | 4.66 | -51 | -17 | 4 |
| | | | | 4.83 | 4.61 | -59 | -13 | 2 |
| | | | | 4.36 | 4.20 | -57 | -3 | -3 |
| | | | | 4.08 | 3.95 | -53 | -27 | 6 |
| | | | | 3.96 | 3.84 | -45 | -37 | 10 |
| | | | | 3.82 | 3.71 | -51 | 4 | -9 |
| | 0.00 | 0.00 | 364 | 5.53 | 5.23 | 8 | -85 | -37 |
| | | | | 5.47 | 5.17 | 8 | -71 | -25 |
| | | | | 4.20 | 4.05 | -5 | -83 | -37 |
| | | | | 4.08 | 3.95 | -9 | -71 | -27 |
| | 0.00 | 0.00 | 267 | 5.50 | 5.20 | -33 | 12 | 58 |
| | 0.00 | 0.00 | 300 | 5.35 | 5.07 | -15 | -91 | -3 |
| | | | | 4.32 | 4.16 | -15 | -87 | -11 |
| | | | | 3.76 | 3.66 | -23 | -79 | -11 |
| | 0.07 | 0.01 | 129 | 5.06 | 4.82 | -61 | -47 | -15 |
| | 0.00 | 0.00 | 373 | 4.66 | 4.47 | 54 | 6 | -7 |
| | | | | 4.60 | 4.42 | 40 | -23 | 8 |
| | | | | 4.54 | 4.36 | 50 | -15 | 2 |
| | | | | 4.53 | 4.35 | 38 | -31 | 12 |
| | | | | 4.06 | 3.93 | 56 | -9 | 2 |
| Main effect Modality: All Language > All Pictures | 0.00 | 0.00 | 37166 | 16.32 | 65535.00 | -61 | -35 | 6 |
| | | | | 13.95 | 65535.00 | -5 | -1 | 66 |
| | | | | 13.22 | 65535.00 | -61 | -3 | -5 |
| | | | | 12.89 | 65535.00 | -51 | 16 | -11 |
| | | | | 12.25 | 65535.00 | -57 | 10 | -9 |

| | | | | | | | | |
|---|---|---|---|---|---|---|---|---|
| | | | | 11.33 | 65535.00 | -51 | -1 | 50 |
| | | | | 11.21 | 65535.00 | -47 | -41 | -1 |
| | | | | 10.49 | 65535.00 | 6 | -19 | 32 |
| | 0.00 | 0.00 | 6080 | 14.91 | 65535.00 | 26 | -67 | -57 |
| | | | | 11.57 | 65535.00 | 18 | -69 | -27 |
| | | | | 11.48 | 65535.00 | 6 | -75 | 4 |
| | | | | 9.76 | 65535.00 | 8 | -89 | 20 |
| | | | | 9.41 | 65535.00 | 28 | -59 | -27 |
| | | | | 8.37 | 7.45 | -7 | -75 | 2 |
| | | | | 8.12 | 7.27 | 14 | -85 | -47 |
| | | | | 7.45 | 6.77 | 16 | -57 | -21 |
| | 0.00 | 0.00 | 884 | 7.23 | 6.60 | 56 | -57 | 48 |
| | | | | 6.24 | 5.81 | 54 | -65 | 42 |
| | 0.00 | 0.00 | 244 | 6.84 | 6.29 | -29 | -45 | 58 |
| | 0.02 | 0.00 | 175 | 6.34 | 5.90 | 32 | -15 | 56 |
| | | | | 4.81 | 4.60 | 20 | -19 | 66 |
| | 0.01 | 0.00 | 201 | 6.20 | 5.78 | 24 | -43 | 64 |
| | | | | 5.08 | 4.83 | 32 | -43 | 56 |
| | 0.00 | 0.00 | 240 | 6.09 | 5.69 | -45 | 2 | -45 |
| | 0.01 | 0.00 | 223 | 5.86 | 5.50 | -21 | -67 | -57 |
| | 0.00 | 0.00 | 441 | 5.72 | 5.38 | -45 | -59 | -39 |
| | | | | 3.88 | 3.76 | -51 | -71 | -37 |
| | | | | 3.75 | 3.64 | -35 | -67 | -35 |
| | | | | 3.61 | 3.51 | -47 | -81 | -37 |
| | 0.02 | 0.00 | 176 | 5.18 | 4.92 | 44 | 20 | 44 |
| | 0.05 | 0.01 | 144 | 4.92 | 4.70 | -19 | -35 | 60 |
| | | | | 4.74 | 4.54 | -29 | -31 | 60 |
| Main effect of Modality: All pictures > All language | 0.00 | 0.00 | 55283 | 38.63 | 65535.00 | 30 | -51 | -11 |
| | | | | 37.90 | 65535.00 | -31 | -51 | -11 |
| | | | | 34.34 | 65535.00 | 30 | -75 | -11 |
| | | | | 30.23 | 65535.00 | 32 | -59 | -13 |
| | | | | 29.43 | 65535.00 | -27 | -67 | -11 |
| | | | | 28.79 | 65535.00 | 20 | -43 | -11 |
| | | | | 28.51 | 65535.00 | 34 | -39 | -15 |
| | | | | 27.97 | 65535.00 | -33 | -89 | 22 |
| | 0.00 | 0.00 | 3382 | 10.53 | 65535.00 | 42 | 10 | 30 |
| | | | | 9.62 | 65535.00 | 32 | -1 | 56 |
| | | | | 9.26 | 65535.00 | 42 | 26 | 22 |
| | | | | 8.61 | 7.62 | 48 | 34 | 16 |
| | | | | 3.80 | 3.69 | 14 | 8 | 72 |
| | 0.00 | 0.00 | 809 | 6.35 | 5.90 | 2 | 50 | -19 |
| | | | | 5.54 | 5.23 | -3 | 42 | -23 |
| | | | | 4.36 | 4.20 | -5 | 70 | -5 |
| | 0.00 | 0.00 | 312 | 5.62 | 5.30 | -29 | -1 | 52 |

| | | | | | | | | |
|---|---|---|---|---|---|---|---|---|
| | | | | 5.35 | 5.07 | -25 | 4 | 60 |
| | 0.07 | 0.02 | 130 | 5.31 | 5.03 | 30 | 34 | -13 |
| | | | | 3.94 | 3.81 | 26 | 38 | -23 |
| Interaction: Coherence (normal and scrambled) and Modality (language and pictures) | 0.00 | 0.00 | 4302 | 67.63 | 7.25 | -13 | -69 | 24 |
| | | | | 63.60 | 7.07 | -11 | -59 | 18 |
| | | | | 43.81 | 6.01 | 6 | -31 | 32 |
| | | | | 43.77 | 6.01 | -7 | -43 | 22 |
| | | | | 39.11 | 5.71 | 14 | -61 | 26 |
| | | | | 36.14 | 5.51 | -25 | -39 | -5 |
| | | | | 34.42 | 5.38 | -11 | -51 | 10 |
| | | | | 31.31 | 5.15 | -5 | -21 | 30 |
| | 0.00 | 0.00 | 2701 | 55.93 | 6.69 | -69 | -31 | 34 |
| | | | | 37.21 | 5.58 | -49 | -69 | 2 |
| | | | | 29.60 | 5.01 | -61 | -61 | 8 |
| | | | | 27.17 | 4.81 | -57 | -73 | 8 |
| | | | | 25.62 | 4.68 | -59 | -61 | 20 |
| | | | | 25.61 | 4.68 | -39 | -49 | 14 |
| | | | | 25.37 | 4.66 | -43 | -59 | 12 |
| | | | | 23.97 | 4.53 | -67 | -47 | 34 |
| | 0.00 | 0.00 | 843 | 42.68 | 5.94 | -43 | 28 | -9 |
| | | | | 26.26 | 4.74 | -39 | 22 | 14 |
| | | | | 22.46 | 4.39 | -55 | 26 | 14 |
| | | | | 18.66 | 4.00 | -57 | 34 | 8 |
| | | | | 15.33 | 3.62 | -57 | 18 | 6 |
| | 0.00 | 0.00 | 836 | 37.92 | 5.63 | -43 | -63 | 48 |
| | | | | 32.40 | 5.23 | -39 | -73 | 44 |
| | | | | 29.35 | 4.99 | -37 | -71 | 52 |
| | 0.00 | 0.00 | 2837 | 34.83 | 5.41 | 52 | -31 | 38 |
| | | | | 31.46 | 5.16 | 54 | -27 | -5 |
| | | | | 31.35 | 5.15 | 32 | -75 | 2 |
| | | | | 31.28 | 5.15 | 54 | -67 | -3 |
| | | | | 30.04 | 5.05 | 58 | -41 | 22 |
| | | | | 29.10 | 4.97 | 60 | -29 | 38 |
| | | | | 24.07 | 4.54 | 62 | -21 | 42 |
| | | | | 23.19 | 4.46 | 54 | -57 | 12 |
| | 0.04 | 0.01 | 129 | 34.78 | 5.41 | 42 | -33 | 12 |
| | 0.00 | 0.00 | 403 | 33.82 | 5.34 | -25 | 66 | -1 |
| | | | | 19.14 | 4.05 | -39 | 60 | -11 |
| | 0.00 | 0.00 | 1026 | 33.13 | 5.29 | 10 | -55 | 66 |
| | | | | 30.48 | 5.08 | -13 | -57 | 68 |
| | | | | 26.38 | 4.75 | -15 | -51 | 56 |
| | | | | 18.17 | 3.95 | 10 | -47 | 54 |
| | | | | 13.12 | 3.34 | 20 | -59 | 60 |

| | | | | | | | | |
|---|---|---|---|---|---|---|---|---|
| | 0.00 | 0.00 | 480 | 30.32 | 5.07 | -59 | -9 | 12 |
| | | | | 26.71 | 4.77 | -33 | -19 | 18 |
| | | | | 23.42 | 4.48 | -49 | -15 | 22 |
| | | | | 23.06 | 4.45 | -41 | -13 | 22 |
| | | | | 13.02 | 3.32 | -35 | -9 | 14 |
| | 0.00 | 0.00 | 370 | 29.31 | 4.99 | 34 | -37 | -7 |
| | | | | 25.85 | 4.70 | 34 | -29 | -15 |
| | | | | 24.32 | 4.56 | 22 | -31 | -17 |
| | | | | 21.44 | 4.29 | 30 | -47 | -1 |
| | | | | 19.51 | 4.09 | 32 | -37 | -17 |
| | | | | 14.79 | 3.55 | 20 | -41 | -13 |
| | 0.00 | 0.00 | 316 | 29.22 | 4.98 | -45 | -47 | 64 |
| | | | | 25.33 | 4.65 | -35 | -51 | 72 |
| | | | | 18.79 | 4.02 | -39 | -39 | 48 |
| | | | | 13.55 | 3.39 | -43 | -37 | 38 |
| | 0.00 | 0.00 | 297 | 27.53 | 4.84 | 50 | 32 | -7 |
| | | | | 24.78 | 4.60 | 42 | 38 | -3 |
| | | | | 15.67 | 3.66 | 56 | 38 | 6 |
| | 0.00 | 0.00 | 725 | 26.76 | 4.78 | -9 | 56 | 36 |
| | | | | 22.22 | 4.37 | 8 | 56 | 30 |
| | | | | 16.96 | 3.81 | 12 | 62 | 14 |
| | | | | 16.88 | 3.80 | -1 | 60 | 24 |
| | | | | 16.63 | 3.78 | 12 | 52 | 18 |
| | | | | 12.18 | 3.21 | -9 | 54 | 16 |
| | 0.00 | 0.00 | 337 | 26.63 | 4.77 | 14 | 20 | -23 |
| | | | | 21.55 | 4.30 | 10 | 38 | -25 |
| | | | | 20.89 | 4.23 | 14 | 8 | -21 |
| | | | | 20.09 | 4.15 | 18 | 36 | -21 |
| | | | | 13.91 | 3.44 | 16 | 42 | -13 |
| | 0.00 | 0.00 | 305 | 23.11 | 4.45 | 64 | -37 | -23 |
| | | | | 20.64 | 4.21 | 64 | -41 | -15 |
| | | | | 18.37 | 3.97 | 62 | -21 | -35 |
| | | | | 14.14 | 3.47 | 60 | -15 | -27 |
| | | | | 12.24 | 3.22 | 70 | -39 | -9 |
| | 0.01 | 0.00 | 161 | 22.37 | 4.38 | -47 | -41 | -13 |
| | 0.05 | 0.01 | 119 | 22.32 | 4.38 | -59 | -47 | -11 |
| | | | | 16.66 | 3.78 | -61 | -41 | -17 |
| | 0.01 | 0.00 | 165 | 20.00 | 4.14 | 12 | 58 | -11 |
| | | | | 18.38 | 3.97 | 20 | 64 | -1 |
| | | | | 12.19 | 3.21 | 14 | 68 | -15 |
| | 0.03 | 0.01 | 134 | 19.55 | 4.10 | -57 | -21 | 6 |
| | | | | 18.48 | 3.98 | -47 | -37 | 12 |
| | | | | 14.01 | 3.45 | -41 | -31 | 14 |
| | 0.01 | 0.00 | 168 | 19.35 | 4.08 | -3 | -65 | -19 |

| | | | | | | | | |
|---|---|---|---|---|---|---|---|---|
| | | | | 19.27 | 4.07 | 2 | -57 | -15 |
| | | | | 12.98 | 3.32 | -7 | -59 | -11 |
| | 0.00 | 0.00 | 370 | 19.30 | 4.07 | 38 | -69 | 44 |
| | | | | 19.28 | 4.07 | 48 | -63 | 40 |
| | | | | 14.91 | 3.57 | 44 | -57 | 48 |
| | 0.00 | 0.00 | 589 | 19.01 | 4.04 | 58 | -7 | 8 |
| | | | | 18.73 | 4.01 | 56 | -27 | 12 |
| | | | | 18.67 | 4.00 | 62 | -13 | 4 |
| | | | | 16.21 | 3.73 | 52 | -13 | 10 |
| | | | | 16.01 | 3.70 | 66 | -23 | 10 |
| | | | | 15.63 | 3.66 | 54 | -1 | -3 |
| | | | | 12.91 | 3.31 | 50 | 8 | -7 |
| | 0.02 | 0.00 | 151 | 17.51 | 3.88 | 44 | 54 | -7 |
| | | | | 15.61 | 3.66 | 38 | 62 | 8 |
| | 0.09 | 0.02 | 104 | 15.70 | 3.67 | 36 | 24 | 16 |
| | | | | 15.61 | 3.65 | 44 | 22 | 14 |
| | | | | 14.42 | 3.51 | 50 | 20 | 24 |
| | | | | 13.97 | 3.45 | 42 | 32 | 8 |
| | | | | 13.43 | 3.38 | 50 | 30 | 12 |
| Language Coherent > Language Scrambled | 0.00 | 0.00 | 1375 | 11.35 | 7.63 | -55 | 8 | -31 |
| | | | | 6.50 | 5.38 | -55 | -3 | -25 |
| | | | | 6.03 | 5.09 | -49 | 16 | -39 |
| | | | | 4.99 | 4.40 | -41 | 24 | -33 |
| | | | | 3.86 | 3.55 | -61 | -19 | -29 |
| | 0.00 | 0.00 | 2772 | 10.12 | 7.16 | -7 | 30 | 58 |
| | | | | 7.88 | 6.14 | -3 | 48 | 32 |
| | | | | 7.73 | 6.06 | -11 | 38 | 58 |
| | | | | 7.61 | 6.00 | 10 | 46 | 50 |
| | | | | 7.38 | 5.87 | -11 | 48 | 50 |
| | | | | 6.76 | 5.53 | 8 | 28 | 60 |
| | | | | 6.06 | 5.11 | -9 | 54 | 40 |
| | | | | 5.98 | 5.06 | 10 | 62 | 32 |
| | 0.00 | 0.00 | 3603 | 8.69 | 6.54 | 54 | 12 | -25 |
| | | | | 8.25 | 6.33 | 50 | 10 | -33 |
| | | | | 7.77 | 6.09 | 56 | -29 | -7 |
| | | | | 7.47 | 5.92 | 48 | 10 | -41 |
| | | | | 7.17 | 5.76 | 60 | -55 | 34 |
| | | | | 6.82 | 5.56 | 52 | -51 | 22 |
| | | | | 6.39 | 5.31 | 60 | -39 | -1 |
| | | | | 6.39 | 5.31 | 62 | -19 | -11 |
| | 0.00 | 0.00 | 2937 | 8.30 | 6.35 | -53 | -25 | -7 |
| | | | | 8.20 | 6.30 | -61 | -53 | 30 |
| | | | | 7.98 | 6.19 | -53 | -55 | 28 |
| | | | | 7.83 | 6.12 | -55 | -51 | 36 |

| | | | | | | | | |
|---|---|---|---|---|---|---|---|---|
| | | | | 7.71 | 6.05 | -55 | -61 | 34 |
| | | | | 7.41 | 5.89 | -61 | -31 | -5 |
| | | | | 6.88 | 5.60 | -41 | -53 | 30 |
| | | | | 6.20 | 5.19 | -65 | -39 | -1 |
| | 0.00 | 0.00 | 628 | 7.19 | 5.78 | -25 | -81 | -33 |
| | 0.00 | 0.00 | 743 | 7.10 | 5.72 | 24 | -75 | -29 |
| | | | | 5.77 | 4.92 | 24 | -85 | -43 |
| | | | | 3.48 | 3.24 | 34 | -61 | -29 |
| | 0.00 | 0.00 | 352 | 6.40 | 5.32 | -39 | 24 | 48 |
| | 0.03 | 0.00 | 146 | 6.35 | 5.29 | -7 | -57 | -41 |
| | | | | 4.70 | 4.19 | 6 | -57 | -41 |
| | 0.00 | 0.00 | 214 | 5.93 | 5.03 | -11 | 8 | 14 |
| | | | | 4.36 | 3.94 | -15 | -1 | 22 |
| | 0.00 | 0.00 | 262 | 5.74 | 4.90 | 10 | 6 | 18 |
| | | | | 5.07 | 4.45 | 8 | 14 | 6 |
| | 0.00 | 0.00 | 321 | 5.39 | 4.67 | -55 | 24 | 4 |
| | | | | 5.25 | 4.57 | -61 | 20 | 14 |
| | | | | 5.21 | 4.55 | -53 | 28 | 12 |
| | | | | 4.69 | 4.18 | -61 | 28 | 10 |
| | 0.00 | 0.00 | 292 | 5.20 | 4.54 | -53 | 38 | -11 |
| | 0.05 | 0.01 | 134 | 5.14 | 4.50 | 50 | 38 | -13 |
| | 0.00 | 0.00 | 257 | 4.97 | 4.38 | -43 | -59 | -7 |
| | | | | 4.74 | 4.22 | -45 | -69 | -11 |
| Picture Coherent > Picture Scrambled | 0.00 | 0.00 | 7938 | 12.44 | 65535.00 | 50 | -59 | 10 |
| | | | | 10.94 | 7.48 | 54 | -69 | -3 |
| | | | | 10.34 | 7.25 | 62 | -25 | 40 |
| | | | | 9.79 | 7.03 | 48 | -75 | 4 |
| | | | | 9.58 | 6.94 | 50 | -51 | 6 |
| | | | | 9.27 | 6.80 | 34 | -69 | 2 |
| | | | | 8.74 | 6.56 | 46 | -63 | -7 |
| | | | | 8.64 | 6.51 | 54 | -37 | 10 |
| | 0.00 | 0.00 | 12566 | 12.34 | 65535.00 | -53 | -67 | 8 |
| | | | | 11.87 | 7.81 | -59 | -61 | 8 |
| | | | | 11.60 | 7.72 | -51 | -75 | 4 |
| | | | | 10.86 | 7.45 | -67 | -33 | 32 |
| | | | | 10.67 | 7.38 | -63 | -29 | 40 |
| | | | | 9.79 | 7.03 | 12 | -55 | 66 |
| | | | | 9.45 | 6.88 | -13 | -57 | 68 |
| | | | | 8.89 | 6.63 | -45 | -43 | -15 |
| | 0.00 | 0.00 | 477 | 9.65 | 6.97 | 28 | -1 | -23 |
| | | | | 7.61 | 6.00 | 22 | -5 | -13 |
| | | | | 4.96 | 4.37 | 36 | -9 | -3 |
| | | | | 4.75 | 4.22 | 32 | -17 | -3 |
| | | | | 4.41 | 3.98 | 16 | -15 | -7 |

| | | | | | | | | |
|---|---|---|---|---|---|---|---|---|
| | 0.00 | 0.00 | 4175 | 9.48 | 6.89 | 6 | 60 | 34 |
| | | | | 9.24 | 6.79 | -3 | 56 | 32 |
| | | | | 7.78 | 6.09 | -9 | 34 | 58 |
| | | | | 7.18 | 5.77 | 10 | 60 | 16 |
| | | | | 6.81 | 5.56 | -9 | 52 | 14 |
| | | | | 6.65 | 5.47 | 10 | 34 | 58 |
| | | | | 6.50 | 5.38 | 24 | -3 | 64 |
| | | | | 6.43 | 5.33 | 6 | 34 | 2 |
| | 0.00 | 0.00 | 632 | 8.98 | 6.67 | 14 | -77 | -47 |
| | | | | 6.67 | 5.48 | 24 | -79 | -35 |
| | | | | 4.93 | 4.36 | 24 | -73 | -25 |
| | | | | 3.39 | 3.17 | 34 | -89 | -31 |
| | 0.00 | 0.00 | 829 | 8.19 | 6.30 | -23 | -79 | -35 |
| | | | | 6.89 | 5.60 | -15 | -79 | -47 |
| | | | | 5.19 | 4.53 | -29 | -63 | -49 |
| | 0.00 | 0.00 | 1669 | 7.35 | 5.86 | -57 | 30 | 6 |
| | | | | 6.94 | 5.63 | -55 | 6 | 26 |
| | | | | 5.92 | 5.02 | -61 | 14 | 26 |
| | | | | 5.85 | 4.98 | -43 | 28 | -9 |
| | | | | 5.26 | 4.58 | -45 | 28 | 4 |
| | | | | 4.91 | 4.34 | -45 | 36 | -5 |
| | | | | 4.49 | 4.03 | -31 | 22 | -17 |
| | 0.00 | 0.00 | 2115 | 7.20 | 5.78 | 50 | 22 | -29 |
| | | | | 7.09 | 5.72 | 52 | 12 | -31 |
| | | | | 6.83 | 5.57 | 56 | 38 | 4 |
| | | | | 6.69 | 5.49 | 48 | 4 | -41 |
| | | | | 6.46 | 5.35 | 50 | 30 | -5 |
| | | | | 6.45 | 5.35 | 56 | 8 | 24 |
| | | | | 5.62 | 4.83 | 44 | 24 | -23 |
| | | | | 5.57 | 4.79 | 28 | 18 | -17 |
| | 0.00 | 0.00 | 348 | 7.09 | 5.72 | -21 | -5 | -15 |
| | | | | 6.15 | 5.16 | -37 | -13 | -1 |
| | 0.00 | 0.00 | 506 | 6.34 | 5.28 | -55 | 10 | -31 |
| | | | | 5.05 | 4.44 | -49 | 6 | -37 |
| | | | | 4.43 | 3.99 | -57 | 8 | -21 |
| | | | | 4.23 | 3.84 | -57 | 8 | -41 |
| | 0.00 | 0.00 | 374 | 6.19 | 5.19 | -23 | -1 | 66 |
| | | | | 5.56 | 4.79 | -21 | -1 | 74 |
| | | | | 4.66 | 4.16 | -27 | -9 | 62 |
| | 0.04 | 0.00 | 138 | 6.08 | 5.12 | 26 | -79 | 40 |
| | 0.00 | 0.00 | 213 | 6.01 | 5.08 | -53 | -27 | -3 |
| | 0.05 | 0.00 | 129 | 5.65 | 4.84 | -17 | -55 | -49 |
| | | | | 4.19 | 3.80 | -19 | -45 | -51 |
| | | | | 4.07 | 3.72 | -7 | -57 | -43 |

| | | | | | | | | |
|---|---|---|---|---|---|---|---|---|
| | 0.00 | 0.00 | 718 | 5.58 | 4.80 | -17 | -17 | 10 |
| | | | | 5.50 | 4.74 | -15 | -31 | 4 |
| | | | | 5.08 | 4.46 | -7 | -7 | 10 |
| | | | | 4.90 | 4.34 | 12 | -5 | 14 |
| | | | | 4.61 | 4.12 | -3 | 8 | -1 |
| | | | | 4.52 | 4.06 | 16 | -27 | 6 |
| | | | | 4.38 | 3.95 | 18 | -19 | 14 |
| | | | | 4.07 | 3.71 | 12 | -13 | 10 |
| | 0.01 | 0.00 | 168 | 5.48 | 4.73 | 14 | -53 | -51 |
| | 0.03 | 0.00 | 147 | 4.97 | 4.38 | 2 | 18 | 24 |
| | | | | 4.09 | 3.73 | -3 | 14 | 30 |
| | 0.75 | 0.10 | 40 | 4.79 | 4.25 | -37 | 32 | 38 |
| | | | | 3.54 | 3.29 | -39 | 34 | 46 |
| | 0.02 | 0.00 | 154 | 4.63 | 4.14 | 34 | -45 | -45 |
| | | | | 4.49 | 4.04 | 42 | -45 | -39 |
| | 0.83 | 0.12 | 35 | 4.48 | 4.02 | -1 | -37 | 2 |
| | 0.22 | 0.02 | 82 | 4.28 | 3.88 | -39 | -45 | -39 |
| | | | | 3.84 | 3.54 | -29 | -43 | -45 |
| | 0.98 | 0.24 | 21 | 4.15 | 3.78 | 24 | 24 | 20 |
| | | | | 3.52 | 3.28 | 18 | 30 | 20 |
| | 1.00 | 0.38 | 13 | 4.13 | 3.76 | -23 | -79 | 34 |
| | 0.72 | 0.10 | 42 | 4.08 | 3.72 | -39 | 22 | -33 |
| | 0.87 | 0.14 | 32 | 3.89 | 3.57 | -21 | -85 | 48 |
| | 1.00 | 0.39 | 12 | 3.85 | 3.54 | 20 | 8 | 8 |
| | 1.00 | 0.68 | 3 | 3.81 | 3.51 | -7 | -5 | -3 |
| | 1.00 | 0.65 | 4 | 3.71 | 3.43 | -1 | -15 | -33 |
| | 1.00 | 0.65 | 4 | 3.57 | 3.32 | -37 | 4 | -1 |
| | 1.00 | 0.68 | 3 | 3.56 | 3.31 | -13 | -47 | -59 |
| | 1.00 | 0.65 | 4 | 3.50 | 3.26 | -3 | 50 | -19 |
| | 1.00 | 0.74 | 2 | 3.50 | 3.26 | 28 | -31 | -35 |
| | 1.00 | 0.65 | 5 | 3.48 | 3.24 | 30 | -61 | -49 |
| | 1.00 | 0.79 | 1 | 3.46 | 3.22 | -41 | -5 | -39 |
| | 1.00 | 0.79 | 1 | 3.36 | 3.15 | 44 | -81 | 34 |
| | 1.00 | 0.79 | 1 | 3.30 | 3.09 | -19 | 40 | 28 |
| Experiment 2 | | | | | | | | |
| ToM > rest | 0.00 | 0.00 | 49590 | 22.31 | 65535.00 | 22 | -95 | -5 |
| | | | | 17.78 | 65535.00 | -29 | -91 | -11 |
| | | | | 17.71 | 65535.00 | -39 | -89 | -11 |
| | | | | 17.30 | 65535.00 | -61 | -11 | -9 |
| | | | | 16.40 | 65535.00 | 10 | -79 | -41 |
| | | | | 16.22 | 65535.00 | -7 | -29 | -5 |
| | | | | 15.08 | 65535.00 | -47 | -61 | -17 |
| | | | | 15.03 | 65535.00 | -49 | 2 | 52 |
| | 0.00 | 0.00 | 1468 | 10.87 | 7.46 | -5 | -1 | 68 |

| | | | | | | | | |
|---|---|---|---|---|---|---|---|---|
| | | | | 8.09 | 6.25 | -11 | 46 | 48 |
| | | | | 7.95 | 6.17 | -9 | 54 | 38 |
| | | | | 5.38 | 4.67 | -9 | 34 | 58 |
| | 0.07 | 0.01 | 126 | 9.65 | 6.97 | -21 | -39 | -43 |
| | 0.00 | 0.00 | 917 | 8.05 | 6.23 | -3 | 62 | -17 |
| | | | | 7.46 | 5.92 | -3 | 48 | -19 |
| | | | | 5.56 | 4.79 | -3 | 32 | -21 |
| | | | | 4.77 | 4.24 | -3 | 22 | -19 |
| | | | | 4.19 | 3.81 | -5 | 12 | -15 |
| | 0.02 | 0.00 | 169 | 7.23 | 5.79 | -3 | -35 | -45 |
| | 0.14 | 0.02 | 102 | 6.55 | 5.41 | -21 | 2 | 6 |
| nToM >rest | 0.00 | 0.00 | 47984 | 22.93 | 65535.00 | 22 | -95 | -5 |
| | | | | 17.30 | 65535.00 | -29 | -91 | -11 |
| | | | | 17.12 | 65535.00 | -39 | -89 | -11 |
| | | | | 16.36 | 65535.00 | -47 | -61 | -17 |
| | | | | 15.81 | 65535.00 | -7 | -29 | -5 |
| | | | | 15.34 | 65535.00 | -43 | -47 | -21 |
| | | | | 15.01 | 65535.00 | 10 | -77 | -39 |
| | | | | 14.65 | 65535.00 | 34 | -91 | -3 |
| | 0.04 | 0.02 | 143 | 10.14 | 7.17 | 20 | -39 | -45 |
| | 0.04 | 0.02 | 148 | 10.09 | 7.15 | -5 | 2 | 28 |
| | 0.13 | 0.03 | 104 | 9.24 | 6.79 | -21 | -37 | -45 |
| | 0.13 | 0.03 | 105 | 5.11 | 4.48 | -3 | 64 | -19 |
| Tom > nToM | 0.00 | 0.00 | 9044 | 15.47 | 65535.00 | -51 | -55 | 24 |
| | | | | 14.31 | 65535.00 | -53 | 10 | -27 |
| | | | | 12.61 | 65535.00 | -57 | 4 | -19 |
| | | | | 12.49 | 65535.00 | -55 | -19 | -9 |
| | | | | 12.38 | 65535.00 | -63 | -13 | -9 |
| | | | | 11.77 | 7.78 | -55 | -5 | -17 |
| | | | | 10.94 | 7.48 | -45 | 12 | -39 |
| | | | | 10.11 | 7.16 | -55 | -39 | 4 |
| | 0.00 | 0.00 | 9091 | 12.93 | 65535.00 | 48 | 20 | -29 |
| | | | | 12.77 | 65535.00 | 52 | -19 | -13 |
| | | | | 12.36 | 65535.00 | 52 | 10 | -23 |
| | | | | 12.13 | 65535.00 | 56 | 2 | -21 |
| | | | | 11.71 | 7.76 | 50 | 18 | -37 |
| | | | | 11.70 | 7.75 | 52 | -53 | 22 |
| | | | | 10.84 | 7.45 | 52 | -27 | -7 |
| | | | | 9.43 | 6.87 | 48 | -47 | 28 |
| | 0.00 | 0.00 | 955 | 12.36 | 65535.00 | 22 | -77 | -29 |
| | | | | 7.54 | 5.96 | 14 | -87 | -43 |
| | 0.00 | 0.00 | 1447 | 12.28 | 65535.00 | -25 | -79 | -31 |
| | | | | 7.65 | 6.02 | -13 | -83 | -43 |
| | 0.00 | 0.00 | 4122 | 10.89 | 7.47 | -9 | 54 | 42 |

| | | | | | | | | |
|---|---|---|---|---|---|---|---|---|
| | | | | 10.08 | 7.15 | -9 | 46 | 46 |
| | | | | 9.91 | 7.08 | -11 | 62 | 30 |
| | | | | 9.47 | 6.89 | -9 | 32 | 58 |
| | | | | 8.59 | 6.49 | 8 | 48 | 50 |
| | | | | 8.43 | 6.41 | -7 | 12 | 68 |
| | | | | 8.30 | 6.35 | 12 | 42 | 54 |
| | | | | 7.71 | 6.05 | 8 | 54 | 32 |
| | 0.00 | 0.00 | 2376 | 10.77 | 7.42 | -5 | -55 | 38 |
| | | | | 10.77 | 7.42 | 6 | -57 | 40 |
| | | | | 10.06 | 7.14 | -13 | -53 | 36 |
| | | | | 6.05 | 5.10 | 8 | -49 | 30 |
| | | | | 4.87 | 4.31 | -5 | -67 | 26 |
| | | | | 3.35 | 3.13 | -23 | -41 | 34 |
| | 0.00 | 0.00 | 670 | 9.27 | 6.80 | -7 | -53 | -41 |
| | | | | 8.62 | 6.50 | 6 | -55 | -43 |
| | | | | 3.86 | 3.55 | 18 | -41 | -45 |
| | 0.00 | 0.00 | 739 | 7.28 | 5.82 | 2 | 48 | -19 |
| | | | | 6.97 | 5.65 | -1 | 60 | -15 |
| | | | | 4.49 | 4.04 | -3 | 30 | -23 |
| | 0.00 | 0.00 | 589 | 7.19 | 5.77 | -49 | 2 | 52 |
| | | | | 6.32 | 5.27 | -43 | 8 | 56 |
| | | | | 5.96 | 5.04 | -57 | -3 | 52 |
| | 0.00 | 0.00 | 1027 | 6.53 | 5.39 | 10 | -31 | 4 |
| | | | | 6.45 | 5.35 | -9 | -29 | 6 |
| | | | | 6.08 | 5.12 | 12 | 4 | 22 |
| | | | | 6.00 | 5.07 | -13 | 10 | 16 |
| | | | | 5.31 | 4.62 | -5 | -23 | -3 |
| | | | | 5.20 | 4.54 | -13 | 2 | 20 |
| | | | | 4.75 | 4.23 | -5 | -31 | -5 |
| | | | | 4.40 | 3.97 | 10 | -1 | 12 |
| | 0.01 | 0.00 | 191 | 6.21 | 5.20 | -31 | -7 | -21 |
| | | | | 5.73 | 4.90 | -21 | -7 | -13 |
| | | | | 4.35 | 3.93 | -11 | -13 | -11 |
| | 0.00 | 0.00 | 1323 | 6.17 | 5.18 | 38 | -87 | -9 |
| | | | | 5.76 | 4.92 | 40 | -59 | -9 |
| | | | | 5.45 | 4.71 | 34 | -91 | -3 |
| | | | | 5.13 | 4.50 | 38 | -43 | -21 |
| | | | | 4.82 | 4.27 | 36 | -83 | 12 |
| | | | | 4.57 | 4.10 | 26 | -97 | -1 |
| | | | | 4.56 | 4.09 | 32 | -65 | -23 |
| | 0.00 | 0.00 | 789 | 5.78 | 4.93 | -33 | -95 | -7 |
| | | | | 5.59 | 4.81 | -41 | -69 | -7 |
| | | | | 4.88 | 4.32 | -43 | -81 | -7 |
| | | | | 4.68 | 4.18 | -31 | -87 | 6 |

| | | | | | | | | |
|---|---|---|---|---|---|---|---|---|
| | 0.04 | 0.00 | 137 | 4.37 | 3.95 | 26 | -59 | 54 |
| | | | | 4.04 | 3.69 | 32 | -63 | 66 |
| | | | | 3.95 | 3.62 | 20 | -67 | 70 |
| | | | | 3.50 | 3.26 | 14 | -61 | 74 |
| F1 | 0.00 | 0.00 | 9892 | 10.74 | 7.41 | -51 | 12 | -21 |
| | 0.00 | 0.00 | | 10.42 | 7.28 | -59 | -13 | -11 |
| | 0.00 | 0.00 | | 10.16 | 7.18 | -55 | -27 | -5 |
| | 0.00 | 0.00 | | 8.89 | 6.63 | -49 | 30 | -7 |
| | 0.00 | 0.00 | | 8.74 | 6.56 | -49 | -23 | -11 |
| | 0.00 | 0.00 | | 8.63 | 6.51 | -53 | -7 | -19 |
| | 0.00 | 0.00 | | 8.59 | 6.49 | -51 | -39 | 6 |
| | 0.00 | 0.00 | | 8.41 | 6.40 | -53 | -59 | 24 |
| | 0.00 | 0.00 | 3677 | 9.25 | 6.79 | 50 | 20 | -27 |
| | 0.00 | 0.00 | | 8.65 | 6.52 | 48 | 10 | -29 |
| | 0.00 | 0.00 | | 7.82 | 6.11 | 50 | -31 | -3 |
| | 0.00 | 0.00 | | 7.33 | 5.85 | 50 | -19 | -7 |
| | 0.00 | 0.00 | | 6.79 | 5.55 | 50 | -11 | -15 |
| | 0.00 | 0.00 | | 6.57 | 5.42 | 48 | 16 | -41 |
| | 0.01 | 0.00 | | 6.42 | 5.33 | 54 | -51 | 26 |
| | 0.17 | 0.02 | | 5.28 | 4.60 | 64 | -3 | -13 |
| | 0.00 | 0.00 | 475 | 8.08 | 6.24 | 6 | -53 | -41 |
| | 0.00 | 0.00 | | 7.82 | 6.11 | -5 | -57 | -43 |
| | 0.00 | 0.00 | 1199 | 7.78 | 6.09 | -11 | 58 | 38 |
| | 0.00 | 0.00 | | 7.38 | 5.88 | -9 | 46 | 52 |
| | 0.22 | 0.02 | | 5.17 | 4.52 | -11 | 68 | 22 |
| | 0.36 | 0.04 | | 4.95 | 4.37 | -7 | 14 | 66 |
| | 0.91 | 0.16 | | 4.29 | 3.89 | -9 | 32 | 62 |
| | 0.00 | 0.00 | 985 | 6.81 | 5.56 | 56 | 28 | 22 |
| | 0.53 | 0.06 | | 4.75 | 4.23 | 38 | -1 | 40 |
| | 0.00 | 0.00 | 854 | 6.73 | 5.51 | 14 | -85 | -43 |
| | 0.00 | 0.00 | | 6.64 | 5.46 | 22 | -83 | -37 |
| | 0.03 | 0.00 | | 5.91 | 5.01 | 18 | -73 | -27 |
| | 0.99 | 0.25 | | 4.06 | 3.71 | 8 | -73 | -29 |
| | 0.00 | 0.00 | 1806 | 6.71 | 5.50 | -47 | -73 | -11 |
| | 0.01 | 0.00 | | 6.42 | 5.33 | -37 | -47 | -21 |
| | 0.01 | 0.00 | | 6.32 | 5.27 | -43 | -53 | -9 |
| | 0.02 | 0.00 | | 6.16 | 5.17 | -47 | -63 | -17 |
| | 0.03 | 0.00 | | 5.94 | 5.03 | -39 | -65 | -7 |
| | 0.03 | 0.00 | | 5.93 | 5.02 | -43 | -81 | -13 |
| | 0.06 | 0.01 | | 5.65 | 4.85 | -45 | -35 | -19 |
| | 0.53 | 0.06 | | 4.75 | 4.22 | -37 | -87 | -11 |
| | 0.00 | 0.00 | 681 | 6.63 | 5.45 | 30 | -73 | 34 |
| | 0.14 | 0.02 | | 5.36 | 4.65 | 40 | -87 | 20 |
| | 0.30 | 0.03 | | 5.04 | 4.43 | 28 | -61 | 50 |

| | | | | | | | | |
|---|---|---|---|---|---|---|---|---|
| | 0.91 | 0.16 | | 4.29 | 3.88 | 26 | -67 | 58 |
| | 0.94 | 0.19 | | 4.22 | 3.83 | 26 | -67 | 42 |
| | 0.02 | 0.00 | 538 | 6.14 | 5.16 | -3 | 46 | -19 |
| | 0.02 | 0.00 | | 6.12 | 5.14 | -5 | 54 | -17 |
| | 0.42 | 0.04 | | 4.88 | 4.32 | -5 | 12 | -17 |
| | 0.52 | 0.06 | | 4.76 | 4.23 | -1 | 30 | -21 |
| | 0.96 | 0.20 | | 4.18 | 3.80 | -1 | 66 | -13 |
| | 0.02 | 0.00 | 184 | 6.02 | 5.08 | -27 | -73 | 30 |
| | 0.03 | 0.00 | 225 | 5.89 | 5.00 | 50 | 34 | -7 |
| | 0.05 | 0.01 | 238 | 5.77 | 4.92 | 44 | -83 | -11 |
| | 0.09 | 0.01 | 700 | 5.52 | 4.76 | -21 | -81 | -35 |
| | 0.15 | 0.02 | | 5.33 | 4.63 | -7 | -79 | -37 |
| | 0.16 | 0.02 | | 5.31 | 4.61 | -11 | -83 | -45 |
| | 0.45 | 0.05 | | 4.84 | 4.29 | -5 | -77 | -29 |
| | 0.64 | 0.07 | | 4.64 | 4.14 | -17 | -75 | -25 |
| | 0.12 | 0.01 | 226 | 5.41 | 4.69 | -21 | 6 | 2 |
| | 0.12 | 0.01 | | 5.41 | 4.69 | -13 | -1 | 20 |
| | 0.60 | 0.07 | | 4.68 | 4.17 | -15 | 6 | 14 |
| | 0.18 | 0.02 | 414 | 5.26 | 4.58 | 34 | -47 | -17 |
| | 0.22 | 0.02 | | 5.18 | 4.53 | 48 | -61 | -15 |
| | 0.44 | 0.05 | | 4.85 | 4.30 | 38 | -39 | -23 |
| | 0.97 | 0.22 | 129 | 4.15 | 3.77 | 20 | -93 | 4 |
| | 0.98 | 0.23 | | 4.11 | 3.75 | 28 | -101 | -1 |
| | 0.99 | 0.27 | | 4.03 | 3.68 | 42 | -93 | 2 |
| | 1.00 | 0.68 | | 3.54 | 3.29 | 30 | -101 | 8 |
| F2 | 0.00 | 0.00 | 5431 | 10.39 | 7.27 | -53 | 10 | -25 |
| | 0.00 | 0.00 | | 9.07 | 6.71 | -49 | -53 | 24 |
| | 0.00 | 0.00 | | 8.86 | 6.62 | -53 | 4 | -31 |
| | 0.00 | 0.00 | | 8.24 | 6.32 | -55 | -59 | 24 |
| | 0.00 | 0.00 | | 7.76 | 6.08 | -39 | -51 | 24 |
| | 0.00 | 0.00 | | 7.43 | 5.91 | -49 | 16 | -31 |
| | 0.00 | 0.00 | | 6.95 | 5.64 | -41 | -57 | 32 |
| | 0.00 | 0.00 | | 6.70 | 5.50 | -55 | -17 | -11 |
| | 0.00 | 0.00 | 8005 | 10.04 | 7.13 | 48 | 10 | -31 |
| | 0.00 | 0.00 | | 9.10 | 6.73 | 8 | -59 | 48 |
| | 0.00 | 0.00 | | 8.98 | 6.67 | 44 | -45 | 30 |
| | 0.00 | 0.00 | | 8.55 | 6.47 | 52 | -53 | 22 |
| | 0.00 | 0.00 | | 7.97 | 6.18 | 50 | -9 | -19 |
| | 0.00 | 0.00 | | 7.84 | 6.12 | -1 | -61 | 46 |
| | 0.00 | 0.00 | | 7.84 | 6.12 | 54 | -1 | -19 |
| | 0.00 | 0.00 | | 7.76 | 6.08 | -9 | -51 | 48 |
| | 0.00 | 0.00 | 193 | 9.51 | 6.91 | 14 | -85 | 2 |
| | 0.00 | 0.00 | 579 | 8.02 | 6.21 | -11 | -57 | -41 |
| | 0.09 | 0.01 | | 5.55 | 4.78 | 12 | -49 | -51 |

| | | | | | | | | |
|---|---|---|---|---|---|---|---|---|
| | 0.29 | 0.04 | | 5.06 | 4.44 | 6 | -53 | -45 |
| | 1.00 | 0.70 | | 3.53 | 3.28 | -9 | -55 | -59 |
| | 0.00 | 0.00 | 233 | 7.04 | 5.69 | 14 | -87 | -43 |
| | 0.40 | 0.06 | | 4.91 | 4.34 | 24 | -79 | -29 |
| | 0.91 | 0.19 | | 4.30 | 3.89 | 16 | -71 | -29 |
| | 0.02 | 0.00 | 106 | 6.15 | 5.17 | -9 | -85 | 2 |
| | 0.03 | 0.01 | 607 | 5.90 | 5.01 | -25 | -81 | -33 |
| | 0.04 | 0.01 | | 5.83 | 4.96 | -11 | -83 | -41 |
| | 0.13 | 0.02 | 141 | 5.39 | 4.67 | -39 | 2 | 56 |
| | 0.82 | 0.16 | | 4.44 | 4.00 | -43 | 10 | 54 |
| | 1.00 | 0.60 | | 3.63 | 3.36 | -53 | -5 | 56 |
| | 0.14 | 0.02 | 123 | 5.35 | 4.65 | 38 | 12 | 48 |
| | 0.26 | 0.04 | 100 | 5.12 | 4.48 | 12 | 44 | 52 |
| | 0.37 | 0.05 | 119 | 4.95 | 4.37 | 52 | -69 | -11 |
| | 0.96 | 0.23 | | 4.17 | 3.79 | 44 | -67 | -3 |
| | 1.00 | 0.42 | | 3.81 | 3.51 | 50 | -83 | -9 |
| | 0.86 | 0.17 | 146 | 4.39 | 3.96 | 34 | -91 | -1 |
| | 1.00 | 0.59 | | 3.64 | 3.38 | 40 | -89 | 16 |
| | 0.98 | 0.26 | 110 | 4.11 | 3.75 | -23 | -59 | 22 |
| | 1.00 | 0.61 | | 3.61 | 3.35 | -9 | -65 | 24 |
| ToM contrast > all factors | 0.00 | 0.01 | 562 | 6.79 | 5.55 | 4 | -17 | 32 |
| | 0.74 | 0.24 | | 4.53 | 4.06 | -5 | -33 | 24 |
| | 0.98 | 0.36 | | 4.10 | 3.74 | 8 | -45 | 18 |
| | 0.00 | 0.01 | 3465 | 6.73 | 5.51 | 36 | 16 | -5 |
| | 0.02 | 0.03 | | 6.10 | 5.13 | 10 | 40 | 22 |
| | 0.02 | 0.03 | | 6.05 | 5.10 | 42 | 48 | 8 |
| | 0.07 | 0.06 | | 5.64 | 4.84 | 10 | 36 | 30 |
| | 0.08 | 0.06 | | 5.58 | 4.80 | 12 | 28 | 26 |
| | 0.08 | 0.06 | | 5.55 | 4.78 | 32 | 12 | 8 |
| | 0.09 | 0.06 | | 5.54 | 4.77 | 30 | 18 | 2 |
| | 0.14 | 0.09 | | 5.34 | 4.64 | 28 | 48 | 10 |
| | 0.19 | 0.09 | 599 | 5.23 | 4.56 | -41 | 14 | -3 |
| | 0.83 | 0.28 | | 4.42 | 3.98 | -29 | 16 | 8 |
| | 0.93 | 0.32 | | 4.25 | 3.86 | -45 | 8 | 2 |
| | 0.96 | 0.35 | | 4.19 | 3.80 | -31 | 28 | 6 |
| | 0.97 | 0.36 | | 4.14 | 3.77 | -33 | 22 | 16 |
| | 0.21 | 0.09 | 170 | 5.20 | 4.54 | 8 | -91 | 18 |
| | 0.22 | 0.09 | 391 | 5.16 | 4.51 | 10 | -77 | 34 |
| | 0.41 | 0.13 | | 4.89 | 4.32 | 16 | -67 | 38 |
| | 0.47 | 0.15 | 100 | 4.82 | 4.27 | -45 | -59 | -31 |
| | 1.00 | 0.46 | | 3.92 | 3.60 | -43 | -63 | -39 |
| | 0.68 | 0.22 | 336 | 4.59 | 4.11 | 16 | -57 | -21 |
| | 0.88 | 0.29 | | 4.34 | 3.92 | -9 | -63 | -15 |
| | 0.93 | 0.32 | | 4.25 | 3.85 | 4 | -59 | -15 |

| | | | | | | | | |
|---|---|---|---|---|---|---|---|---|
| | 0.99 | 0.44 | | 3.97 | 3.64 | 16 | -57 | -11 |
| | 1.00 | 0.45 | | 3.95 | 3.62 | -9 | -65 | -23 |
| | 1.00 | 0.59 | | 3.71 | 3.43 | 12 | -67 | -15 |
| | 0.83 | 0.28 | 206 | 4.42 | 3.98 | -9 | -87 | 20 |
| | 0.93 | 0.32 | | 4.25 | 3.85 | -15 | -105 | 6 |
| | 1.00 | 0.49 | | 3.88 | 3.57 | -9 | -95 | 12 |
| | 0.91 | 0.32 | 173 | 4.28 | 3.88 | -5 | -83 | -7 |
| | 0.98 | 0.37 | | 4.08 | 3.72 | -9 | -67 | -1 |
| | 1.00 | 0.56 | | 3.80 | 3.50 | 10 | -73 | 2 |
| | 1.00 | 0.57 | | 3.77 | 3.48 | -7 | -75 | 2 |
| | 1.00 | 0.88 | | 3.40 | 3.18 | -9 | -77 | -15 |
| All Factors > ToM contrast | 0.01 | 0.02 | 452 | 6.56 | 5.41 | 30 | -71 | 32 |
| | 0.01 | 0.02 | 226 | 6.53 | 5.39 | 8 | -51 | -39 |
| | 0.46 | 0.35 | | 4.83 | 4.28 | 14 | -53 | -51 |
| | 0.03 | 0.05 | 404 | 5.96 | 5.05 | -33 | -73 | 28 |
| | 0.91 | 0.47 | | 4.29 | 3.88 | -37 | -93 | 28 |
| | 0.99 | 0.58 | | 4.00 | 3.66 | -33 | -83 | 48 |
| | 0.16 | 0.25 | 342 | 5.30 | 4.61 | -55 | -15 | -11 |
| | 0.39 | 0.35 | | 4.91 | 4.34 | -61 | -9 | -11 |
| | 0.26 | 0.33 | 1334 | 5.09 | 4.47 | -49 | -57 | 28 |
| | 0.49 | 0.35 | | 4.80 | 4.26 | -55 | -65 | 18 |
| | 0.79 | 0.39 | | 4.47 | 4.02 | -65 | -59 | 8 |
| | 0.94 | 0.50 | | 4.22 | 3.83 | -65 | -53 | 2 |
| | 1.00 | 0.67 | | 3.85 | 3.54 | -65 | -63 | 24 |
| | 1.00 | 0.67 | | 3.85 | 3.54 | -39 | -59 | 32 |
| | 1.00 | 0.71 | | 3.67 | 3.40 | -49 | -39 | 6 |
| | 0.29 | 0.33 | 175 | 5.05 | 4.44 | -49 | -59 | -11 |
| | 0.44 | 0.35 | 246 | 4.85 | 4.29 | -55 | -5 | -27 |
| | 0.91 | 0.47 | | 4.30 | 3.89 | -49 | 4 | -27 |
| | 1.00 | 0.67 | | 3.83 | 3.53 | -57 | 4 | -19 |
| | 1.00 | 0.75 | | 3.57 | 3.32 | -53 | 14 | -29 |
| | 1.00 | 0.86 | | 3.48 | 3.24 | -63 | -9 | -31 |
| | 0.61 | 0.35 | 101 | 4.66 | 4.16 | -11 | -29 | 12 |
| | 0.98 | 0.55 | | 4.09 | 3.73 | -11 | -27 | -3 |
| | 0.64 | 0.35 | 107 | 4.64 | 4.14 | 26 | -53 | 48 |
| | 1.00 | 0.73 | | 3.62 | 3.35 | 30 | -63 | 54 |
| | 0.68 | 0.35 | 144 | 4.59 | 4.11 | 54 | 8 | -27 |
| | 1.00 | 0.71 | | 3.67 | 3.40 | 54 | 2 | -17 |
| | 0.70 | 0.35 | 111 | 4.57 | 4.09 | -27 | -39 | -15 |
| | 0.96 | 0.51 | | 4.18 | 3.80 | -35 | -35 | -17 |
| | 0.70 | 0.35 | 176 | 4.57 | 4.09 | -9 | 56 | -13 |
| | 0.93 | 0.47 | | 4.26 | 3.86 | -1 | 64 | -7 |
| | 0.78 | 0.39 | 154 | 4.48 | 4.03 | -35 | 8 | 28 |
| | 1.00 | 0.68 | | 3.82 | 3.51 | -41 | -1 | 36 |

| | | | | | | | | |
|---|---|---|---|---|---|---|---|---|
| | 0.84 | 0.43 | 314 | 4.40 | 3.97 | 2 | -29 | 64 |
| | 0.91 | 0.47 | | 4.30 | 3.89 | -7 | -23 | 60 |
| | 1.00 | 0.71 | | 3.77 | 3.47 | -11 | -23 | 70 |
| | 1.00 | 0.71 | | 3.67 | 3.40 | 10 | -25 | 76 |
| | 1.00 | 0.86 | | 3.47 | 3.24 | 8 | -23 | 58 |
| | 0.91 | 0.47 | 106 | 4.30 | 3.89 | -15 | 58 | 32 |
| | 0.98 | 0.55 | | 4.07 | 3.72 | -9 | 48 | 44 |
| | 1.00 | 0.67 | | 3.87 | 3.56 | -9 | 42 | 52 |
| | 0.96 | 0.51 | 354 | 4.16 | 3.79 | 52 | -43 | 20 |
| | 0.98 | 0.55 | | 4.07 | 3.72 | 42 | -43 | 24 |
| | 0.98 | 0.55 | | 4.07 | 3.71 | 50 | -55 | 26 |
| | 0.99 | 0.55 | | 4.06 | 3.71 | 46 | -57 | 14 |
| | 1.00 | 0.72 | | 3.65 | 3.38 | 52 | -63 | 8 |
| | 1.00 | 0.73 | | 3.63 | 3.37 | 58 | -51 | 22 |
| F1>F2 | 0.00 | 0.00 | 695 | 8.03 | 6.21 | -53 | 38 | 14 |
| | 0.15 | 0.03 | | 5.35 | 4.65 | -45 | 26 | 20 |
| | 0.99 | 0.35 | | 4.03 | 3.68 | -35 | 28 | 10 |
| | 1.00 | 0.71 | | 3.59 | 3.33 | -39 | 24 | 2 |
| | 0.00 | 0.00 | 2262 | 7.99 | 6.19 | -37 | -41 | -19 |
| | 0.00 | 0.00 | | 7.94 | 6.17 | -29 | -7 | -13 |
| | 0.00 | 0.00 | | 7.49 | 5.93 | -25 | -15 | -13 |
| | 0.05 | 0.02 | | 5.77 | 4.92 | -17 | -11 | -19 |
| | 0.05 | 0.02 | | 5.76 | 4.92 | -23 | -33 | -19 |
| | 0.05 | 0.02 | | 5.73 | 4.90 | -21 | -3 | -15 |
| | 0.13 | 0.03 | | 5.38 | 4.67 | -33 | 2 | -17 |
| | 0.13 | 0.03 | | 5.38 | 4.67 | -51 | -49 | -19 |
| | 0.00 | 0.00 | 501 | 7.57 | 5.98 | -31 | 32 | -13 |
| | 0.86 | 0.19 | | 4.40 | 3.97 | -49 | 24 | -15 |
| | 0.91 | 0.20 | | 4.32 | 3.91 | -51 | 16 | -11 |
| | 0.00 | 0.00 | 578 | 6.64 | 5.46 | 24 | -1 | -15 |
| | 0.15 | 0.03 | | 5.34 | 4.64 | 20 | -15 | -23 |
| | 0.24 | 0.05 | | 5.15 | 4.51 | 22 | 14 | -23 |
| | 0.49 | 0.10 | | 4.81 | 4.27 | 32 | 8 | -21 |
| | 0.05 | 0.02 | 343 | 5.74 | 4.91 | -5 | -51 | 4 |
| | 0.19 | 0.04 | | 5.25 | 4.58 | -5 | -55 | 12 |
| | 0.79 | 0.16 | | 4.48 | 4.03 | -17 | -49 | 6 |
| | 1.00 | 0.55 | | 3.77 | 3.48 | -7 | -41 | -1 |
| | 0.20 | 0.04 | 149 | 5.23 | 4.56 | -33 | -5 | -31 |
| | 0.62 | 0.11 | | 4.67 | 4.17 | -29 | 2 | -43 |
| | 0.30 | 0.06 | 105 | 5.05 | 4.44 | -3 | 4 | 28 |
| | 0.54 | 0.10 | 118 | 4.75 | 4.23 | -39 | 6 | 24 |
| | 0.57 | 0.10 | 191 | 4.72 | 4.20 | 2 | -73 | -29 |
| | 0.87 | 0.19 | | 4.37 | 3.95 | -5 | -75 | -19 |
| | 1.00 | 0.71 | | 3.58 | 3.33 | 10 | -83 | -33 |

| | | | | | | | | |
|---|---|---|---|---|---|---|---|---|
| | 0.74 | 0.15 | 120 | 4.54 | 4.07 | 26 | -29 | -21 |
| | 0.90 | 0.20 | | 4.32 | 3.91 | 30 | -37 | -23 |
| F2 > F1 | 0.00 | 0.00 | 2398 | 9.09 | 6.72 | 10 | -55 | 48 |
| | 0.01 | 0.01 | | 6.32 | 5.27 | -3 | -63 | 44 |
| | 0.04 | 0.02 | | 5.84 | 4.97 | -7 | -53 | 50 |
| | 0.14 | 0.05 | | 5.37 | 4.66 | 16 | -47 | 40 |
| | 0.93 | 0.27 | | 4.26 | 3.86 | -21 | -83 | 28 |
| | 0.98 | 0.32 | | 4.08 | 3.72 | 2 | -55 | 62 |
| | 0.99 | 0.33 | | 4.06 | 3.70 | -7 | -83 | 40 |
| | 0.99 | 0.36 | | 4.00 | 3.66 | -1 | -87 | 24 |
| | 0.00 | 0.00 | 467 | 8.45 | 6.42 | 12 | -85 | 4 |
| | 0.00 | 0.00 | | 7.15 | 5.75 | 12 | -75 | -7 |
| | 0.00 | 0.00 | 381 | 7.56 | 5.97 | 52 | 10 | -33 |
| | 0.01 | 0.01 | | 6.20 | 5.19 | 56 | 2 | -35 |
| | 0.01 | 0.00 | 352 | 6.55 | 5.40 | -9 | -85 | 2 |
| | 0.19 | 0.06 | | 5.24 | 4.57 | -15 | -79 | -9 |
| | 0.15 | 0.05 | 917 | 5.33 | 4.63 | 50 | -57 | 30 |
| | 0.16 | 0.05 | | 5.33 | 4.63 | 48 | -59 | 18 |
| | 0.27 | 0.07 | | 5.09 | 4.47 | 40 | -53 | 12 |
| | 0.40 | 0.09 | | 4.91 | 4.34 | 58 | -55 | 26 |
| | 0.75 | 0.18 | | 4.53 | 4.06 | 40 | -45 | 26 |
| | 0.75 | 0.18 | | 4.53 | 4.06 | 52 | -51 | 16 |
| | 0.96 | 0.29 | | 4.20 | 3.81 | 58 | -55 | 38 |
| | 1.00 | 0.38 | | 3.93 | 3.61 | 50 | -43 | 30 |
| | 0.20 | 0.06 | 727 | 5.22 | 4.56 | -49 | -61 | 34 |
| | 0.26 | 0.06 | | 5.11 | 4.48 | -51 | -51 | 24 |
| | 0.50 | 0.11 | | 4.79 | 4.26 | -41 | -63 | 14 |
| | 0.56 | 0.12 | | 4.73 | 4.21 | -37 | -51 | 22 |
| | 0.72 | 0.17 | | 4.56 | 4.09 | -55 | -61 | 24 |
| | 1.00 | 0.49 | | 3.80 | 3.50 | -45 | -71 | 4 |
| | 0.32 | 0.08 | 643 | 5.02 | 4.41 | 20 | -85 | 28 |
| | 0.35 | 0.08 | | 4.98 | 4.39 | 34 | -85 | 36 |
| | 0.94 | 0.27 | | 4.25 | 3.85 | 22 | -79 | 34 |
| | 0.94 | 0.27 | | 4.24 | 3.84 | 26 | -79 | 44 |
| | 1.00 | 0.75 | | 3.47 | 3.24 | 22 | -75 | 52 |
| | 0.83 | 0.21 | 145 | 4.44 | 3.99 | -61 | -9 | -35 |
| | 0.98 | 0.32 | | 4.10 | 3.74 | -55 | -1 | -31 |
| | 1.00 | 0.45 | | 3.84 | 3.53 | -57 | 6 | -23 |
| | 0.91 | 0.25 | 107 | 4.32 | 3.91 | 42 | -67 | -3 |
| | 1.00 | 0.38 | | 3.93 | 3.61 | 46 | -73 | 4 |

References

Chiou, R., Humphreys, G. F., & Lambon Ralph, M. A. (2020). Bipartite Functional Fractionation within the Default Network Supports Disparate Forms of Internally Oriented Cognition. *Cerebral Cortex (New York, N.Y.: 1991)*, *30*(10), 5484–5501. https://doi.org/10.1093/cercor/bhaa130

Deen, B., Koldewyn, K., Kanwisher, N., & Saxe, R. (2015). Functional Organization of Social Perception and Cognition in the Superior Temporal Sulcus. *Cerebral Cortex*, *25*(11), 4596–4609. https://doi.org/10.1093/cercor/bhv111

Dodell-Feder, D., Koster-Hale, J., Bedny, M., & Saxe, R. (2011). fMRI item analysis in a theory of mind task. *NeuroImage*, *55*(2), 705–712. https://doi.org/10.1016/j.neuroimage.2010.12.040

Hodgson, V. J., Lambon Ralph, M. A., & Jackson, R. L. (2023). The cross-domain functional organization of posterior lateral temporal cortex: Insights from ALE meta-analyses of 7 cognitive domains spanning 12,000 participants. *Cerebral Cortex*, *33*(8), 4990–5006. https://doi.org/10.1093/cercor/bhac394

Humphreys, G. F., Halai, A. D., Branzi, F. M., & Lambon Ralph, M. A. (2024). The left posterior angular gyrus is engaged by autobiographical recall not object-semantics, or event-semantics: Evidence from contrastive propositional speech production. *Imaging Neuroscience*, *2*, 1–19. https://doi.org/10.1162/imag_a_00116

Humphreys, G. F., Jung, J., & Lambon Ralph, M. L. R. (2022). The convergence and divergence of episodic and semantic functions across lateral parietal cortex. *Cerebral Cortex*, *32*(24), 5664–5681.

Humphreys, G. F., & Lambon Ralph, M. A. (2025). Mapping the task-general and task-specific neural correlates of speech production: Meta-analysis and fMRI direct comparisons of category fluency and picture naming. *Imaging Neuroscience*, *3*, IMAG.a.154. https://doi.org/10.1162/IMAG.a.154

Jackson, R. L. (2021). The neural correlates of semantic control revisited. *NeuroImage*, *224*, 117444. https://doi.org/10.1016/j.neuroimage.2020.117444

Shain, C., Paunov, A., Chen, X., Lipkin, B., & Fedorenko, E. (2023). No evidence of theory of mind reasoning in the human language network. *Cerebral Cortex*, *33*(10), 6299–6319. https://doi.org/10.1093/cercor/bhac505
